# Protenix - Advancing Structure Prediction Through a Comprehensive AlphaFold3 Reproduction

**DOI:** 10.1101/2025.01.08.631967

**Authors:** ByteDance AML AI4Science Team, Xinshi Chen, Yuxuan Zhang, Chan Lu, Wenzhi Ma, Jiaqi Guan, Chengyue Gong, Jincai Yang, Hanyu Zhang, Ke Zhang, Shenghao Wu, Kuangqi Zhou, Yanping Yang, Zhenyu Liu, Lan Wang, Bo Shi, Shaochen Shi, Wenzhi Xiao

## Abstract

In this technical report, we present Protenix, a comprehensive reproduction of AlphaFold3 (AF3), aimed at advancing the field of biomolecular structure prediction. Protenix tackles the challenges of predicting complex interactions involving proteins, ligands, and nucleic acids, while enhancing accessibility and reproducibility. Across diverse benchmarks, including PoseBusters V2, low-homology PDB sets, and CASP15 RNA, Protenix achieves state-of-the-art performance in protein-ligand, protein-protein, and protein-nucleic acid predictions. We also address limitations, such as potential memorization effects, and outline future directions for improvement. By open-sourcing Protenix, we aim to democratize advanced structure prediction tools and accelerate interdisciplinary research in computational biology and drug discovery.

## 1 Introduction

Understanding the three-dimensional structures of biological molecules has revolutionized our ability to study and manipulate molecular functions. Over recent decades, significant progress in protein structure prediction has been made through various contributions [1, 2, 3, 4, 5, 6, 7, 8, 9, 10]. However, predicting complex structures that involve multiple biomolecules—such as protein-ligand, protein-nucleic acid, and antibody-antigen interactions—poses additional challenges. Recent works like NeuralPLexer [11], Umol [12], and RoseTTAFold-AA [13] have targeted these complex prediction tasks. Among these, AlphaFold 3 (AF3) [4] has set a new milestone, representing a significant leap forward in this domain.

Despite the impressive advances of AF3, limited accessibility has constrained its broader adoption within the research community. The absence of code and certain ambiguities and typographical errors in the AF3 paper present additional challenges for machine learning and computational biology researchers seeking to reproduce or improve the model. Open-source initiatives like HelixFold3 [14] and Chai-1 [15] have made significant strides in democratizing access to these advanced models. However, the lack of comprehensive training code and preprocessed data remains a barrier for researchers aiming to fully reconstruct and utilize these models. To address these challenges, we introduce *Protenix*, designed to lower these barriers and better support the interdisciplinary research community.

Below, we outline our key contributions:

- **Model performance.** We benchmark *Protenix* against AF3 [4], Alphafold-Multimer 2.3 (AF2.3) [3], and RoseTTAFold2NA (RF2NA) [16]. *Protenix* achieves strong performance in predicting structures across different molecular types (Figure 1). As a fully open-source model, it empowers researchers to generate novel predictions and fine-tune the model for specialized applications.
- **Methodology.** We implement *Protenix* based on the descriptions provided in AF3 as part of our reproduction effort. We refine several ambiguous steps, correct typographical errors, and make targeted adjustments based on our observations of the model’s behavior. By sharing our reproduction experience, we aim to support the community’s efforts to build on these improvements and drive further advancements in the field.
- **Accessibility.** We have open-sourced *Protenix*, providing model weights, inference code, and trainable code for research purposes, as detailed in Section 4.

**Figure 1:**
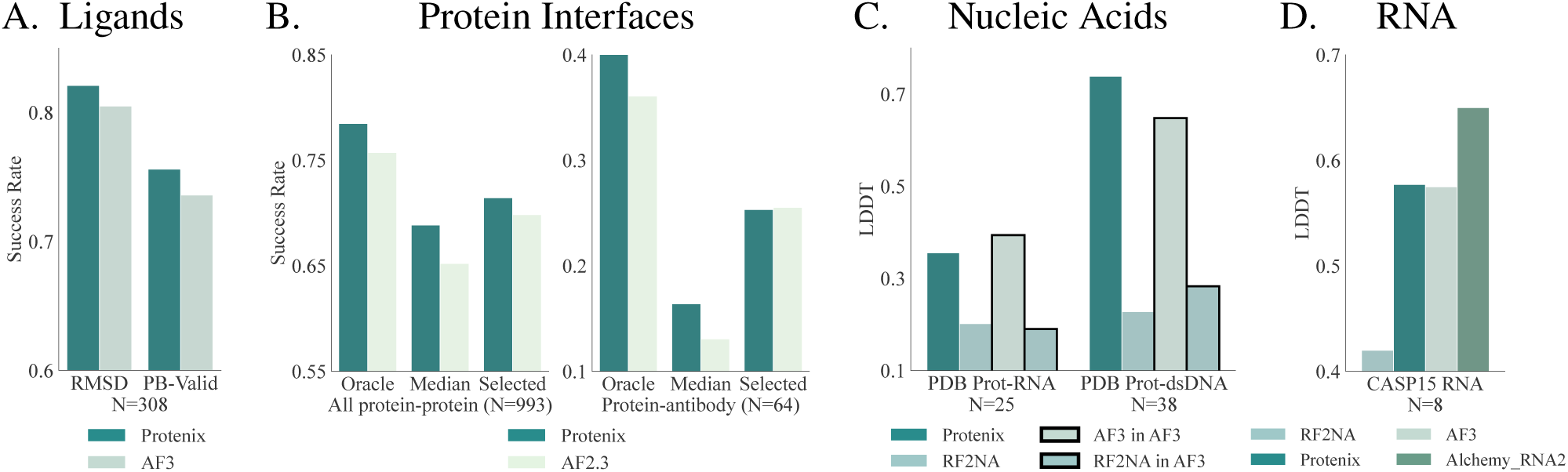
Model performance across several benchmarks. Unless specified otherwise, performances are evaluated on the top-ranked prediction of each model based on its ranking score. **A.** Performance of *Protenix* and AF3 on PoseBusters V2. The RMSD success rate is the percentage of predictions where the pocket-aligned ligand RMSD is 2 Å or lower. The PB-Valid success rate is the percentage of predictions that meet all 18 validity criteria specified by the PoseBusters test-suite. *N* is the number of structures. **B & C.** Performance of *Protenix*, RF2NA, AF2.3, and AF3 on filtered subsets of the Low Homology Recent PDB Set across different interface types. For protein interfaces, the success rate is defined as the percentage of interfaces with a DockQ score above 0.23, while for protein-nucleotide interfaces, the interface LDDT score is reported. Metrics are averaged within and across interface clusters; *N* represents the number of clusters. “AF3 in AF3” and “RF2NA in AF3” indicate metrics of AF3 and RF2NA as reported by AF3, respectively. **D.** Performance of *Protenix*, AIChemy_RNA2, RF2NA, and AF-3 on CASP15 RNA. *N* represents the number of structures.

## 2 Results

### 2.1 Overview

*Protenix* is trained using experimental structures curated from the Protein Data Bank (PDB) [17] with a cutoff date of September 30, 2021 and predicted structures of protein monomers with AlphaFold2 [2] and OpenFold [10] (See details in Table 3 and Appendix A.1). We benchmark *Protenix* on different evaluation datasets to assess its performance on different types of molecules. Details of the evaluation datasets are provided in Appendix A.2. We compare *Protenix*’s performance with that of AF3, AF2.3, and RF2NA.

In contrast to AF3, which trains two separate models with different data cutoffs (September 30, 2019, for Posebusters, and September 30, 2021, for other datasets), we opt to train a single model due to resource limitations. Targets from PoseBusters Benchmark Set Verison 2 (PoseBusters V2) are excluded to avoid data leakage. We acknowledge the differences in the cutoff dates and present an analysis in Section 2.2 to distinguish genuine improvements on Posebusters from potential data overlap.

For each PDB entry, we follow AF3’s inference setup, generating 25 predictions using 5 model seeds, with each seed producing 5 diffusion samples. The predictions are ranked using confidence scores.

In the following sections, we present detailed benchmark results, along with in-depth analysis of low-performing cases to highlight potential directions for further improvement.

### 2.2 Ligands

We evaluate *Protenix* on the PoseBusters Version 2 benchmark set [18] and compare its performance to that of AF3 [4]. We follow the evaluation procedures outlined in AF3 paper to ensure consistency and fairness of comparisons. Details of the evaluation procedure are provided in Appendix B.

**RMSD success rate.** The success rate is defined as the percentage of predictions for which the pocket aligned ligand root-mean-square deviation (RMSD), as defined in [4], between the ground truth and the prediction is no greater than 2 Å. As shown in Figure 2[A], *Protenix* outperforms AF3-2019 in terms of success rate, achieving slightly higher rates for both RMSD and PB-Valid metrics. This suggests that *Protenix* represents the current state-of-the-art (SOTA) model for the protein-ligand cofolding task.

**Figure 2:**
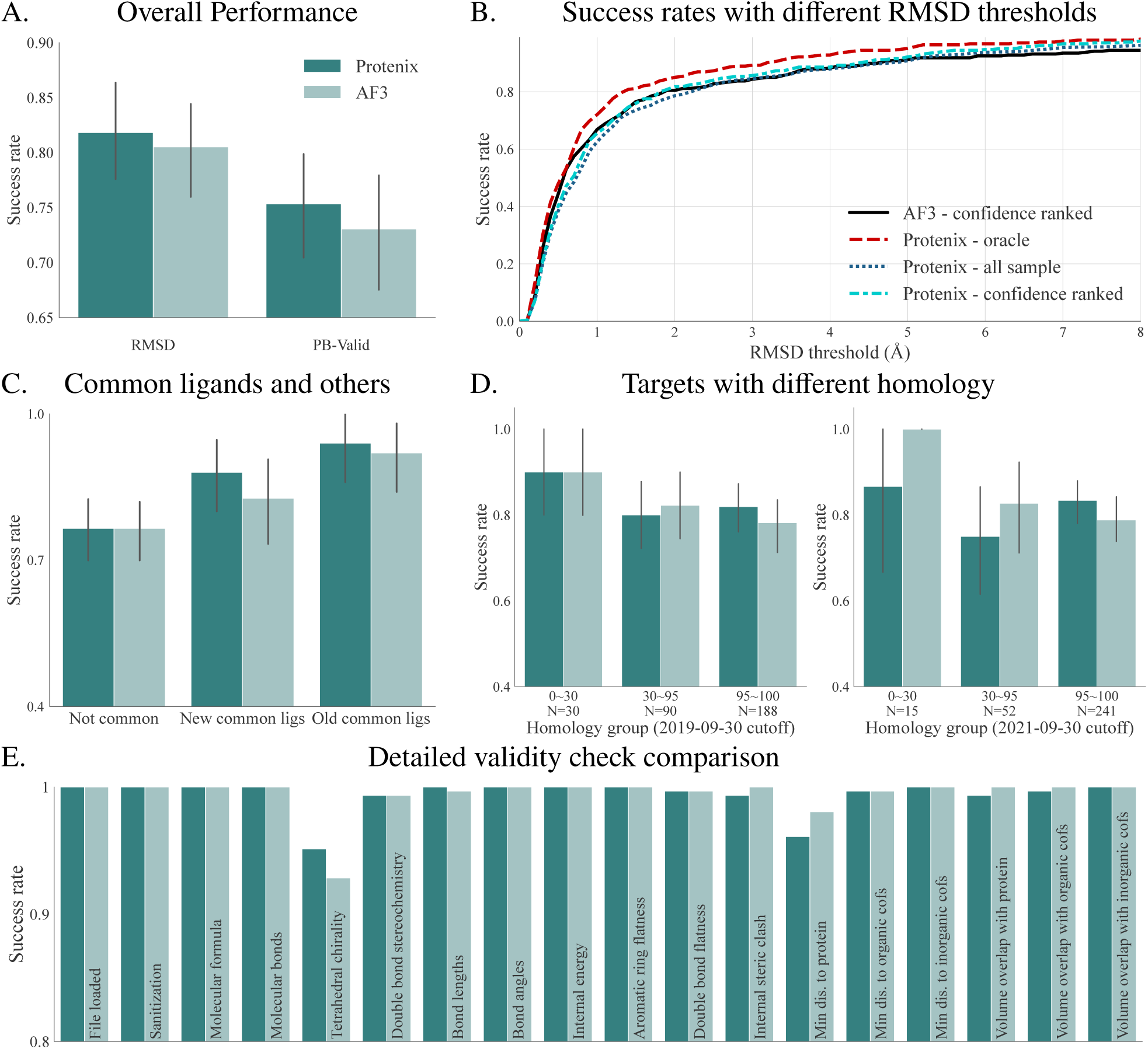
PoseBusters evaluation. **A.** Comparison of protein-ligand binding success rate and the structure validity between *Protenix* and AF3. The metrics of AF3 are computed using the PoseBusters test suite and the predictions provided by AF3. These metrics are consistent with the results reported by AF3. **B.** A comparison of RMSD success rates under different thresholds. In addition to the results for AF3, we present the corresponding success rates of our model under various sample selection methods. “Oracle” refers to the samples with the lowest RMSD; “all sample” presents the metrics computed using all 25 samples generated by *Protenix*; and “confidence ranked” provides the metrics for the samples chosen based on the confidence head. **C.** Comparison of results on ligands split into different categories. We follow AF3’s definition [4] to identify 50 AF3 defined Natural Common Ligands (NCL), which occur over 100 times in PDB and are visually identified as natural, and call them “old natural common ligands”. Furthermore, we select other 75 NCL, referred as “new natural common ligands”, in PoseBusters V2 that meet the same criteria within our training set. **D.** Comparison of results on targets with low, moderate, and high protein sequence homology. We display the homology computed with the 2019-cutoff training set (left) and the 2021-cutoff training set (right). Note that the sample size in each homology group is slightly different from AF3 report [4], due to the potential differences in alignment methods used. **E.** Comparison of detailed validity check on PoseBusters V2.

Figure 2[B] shows success rates across varying RMSD thresholds, offering a more detailed view of model performance. The *Protenix* - oracle configuration (red dashed line) achieves the highest success rate at all RMSD thresholds, consistently outperforming the AF3-2019 confidence ranked (black solid line). The *Protenix* - confidence ranked and *Protenix* - all sample configurations closely match AF3’s performance, highlighting *Protenix*’s ability to generate high-quality protein-ligand cofolding predictions. It’s worth noting that the selected results still lag behind the best candidates among all samples, suggesting room for improvement with a better sample ranker.

**Similarity analysis.** We note that *Protenix* is trained with PDB cut-off 2021-09-30, hence the improvement of performance may be attributed to additional training data. To avoid potential data leakage issue, we remove identical PDB IDs from our training set. Additionally, we perform a similarity analysis as shown in Figure 2[C] and [D]. These figures illustrate that compared to AF3-2019, *Protenix* performs better on common ligands, likely benefiting from the additional training data. However, since they exhibit the same performance on non-common ligands, we can conclude that our model generalizes as well as AF3-2019. On the target side, *Protenix* outperforms AF3-2019 on high homology group but performs slightly worse on other targets.

**Physical plausibility.** We also conduct the PoseBusters V2 plausibility test, using the PoseBusters test-suite version 0.2.7 [18]. Apart from RMSD metrics, PB-valid reflects the ability of a model to generate physically valid structures. As shown in Figure 2[E], we check on 18 validity criteria, which can primarily be categorized into intramolecular validity, intermolecular validity, chemical validity and consistency, etc. Similar to AF3, the model cannot consistently generate samples with correct chirality and without steric clashes, hence we follow AF3 and incorporate a penalty term to mitigate such stereochemistry violations in our score function when we perform sample ranking. This penalty term has successfully improved the PB-valid success rate by over 10%, though there still exists a roughly 5% chance that we fail to predict the correct chirality.

**Ligand case studies.** We inspect some challenging cases to assess *Protenix*’s ability to predict protein-ligand interactions. In cases where there are no close structural neighbors in the training set, *Protenix* demonstrates highly accurate predictions, as shown in Figure 3. Specifically, Figure 3[A] shows precise binding interactions with endogenous ligands like sulfooxy-heptanoic acid (7WUX), while Figure 3[B] illustrates accurate predictions for novel inhibitors (7URD), even when the ligands are larger and have tight binding conformations. These examples highlight *Protenix*’s ability to predict diverse and complex interactions with high accuracy.

**Figure 3:**
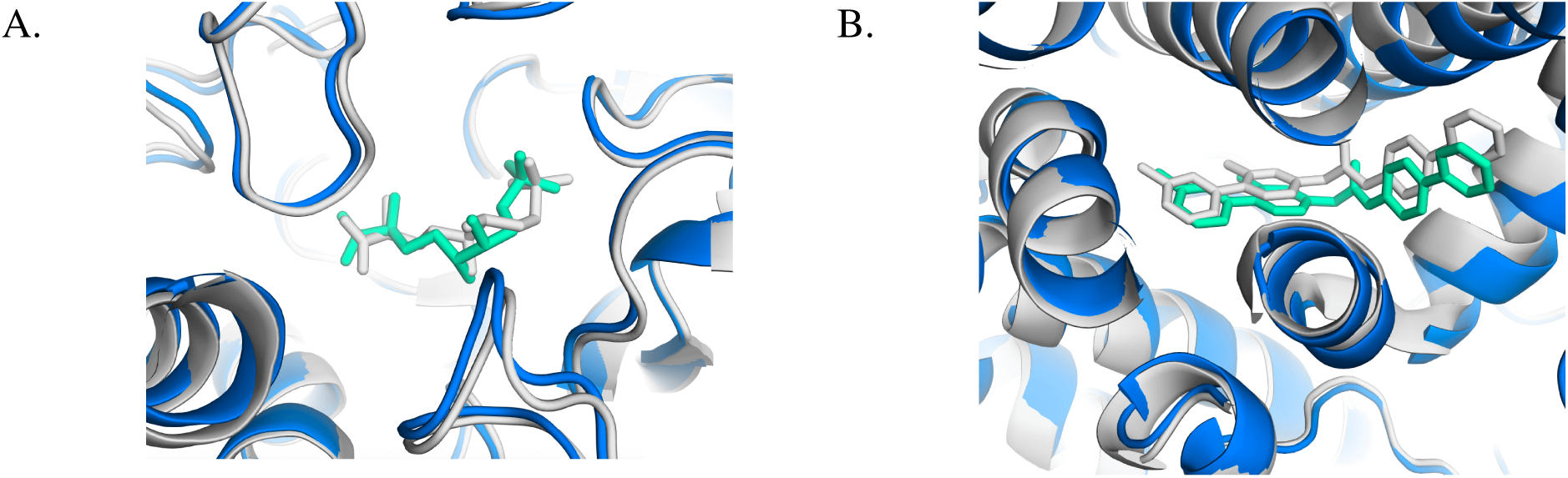
Novel protein-ligand interactions. Predicted protein chains are shown in blue, predicted ligands show in turquoise, and the ground truth is shown in gray. **A.** AziU3/U2 complexed with (5S,6S)-O7-sulfo DADH from Streptomyces sahachiroi (PDB: 7WUX). **B.** Human PORCN in complex with LGK974, a clinical-stage inhibitor with 30 heavy atoms (PDB: 7URD).

We also inspect the cases where the RMSD of predicted ligands exceeds 2 Å and find that, in some instances, the predictions are reasonable. For example, in Figure 4[A], *Protenix* predicts an alternative ligand conformation with the fluorine atom oriented oppositely (7LOE), though both conformations share similar occupancies (42% vs. 58%). In Figure 4[B], *Protenix*’s prediction results in a larger RMSD, but the predicted binding mode still maintains a key interaction between the ligand and two aspartates (5SAK). Interestingly, the predicted binding mode resembles another fragment in the PDB for the same target (5SAN). These cases reveal areas for improvement in the evaluation metrics. They also show that *Protenix* is able to capture key binding features even in more challenging predictions.

**Figure 4:**
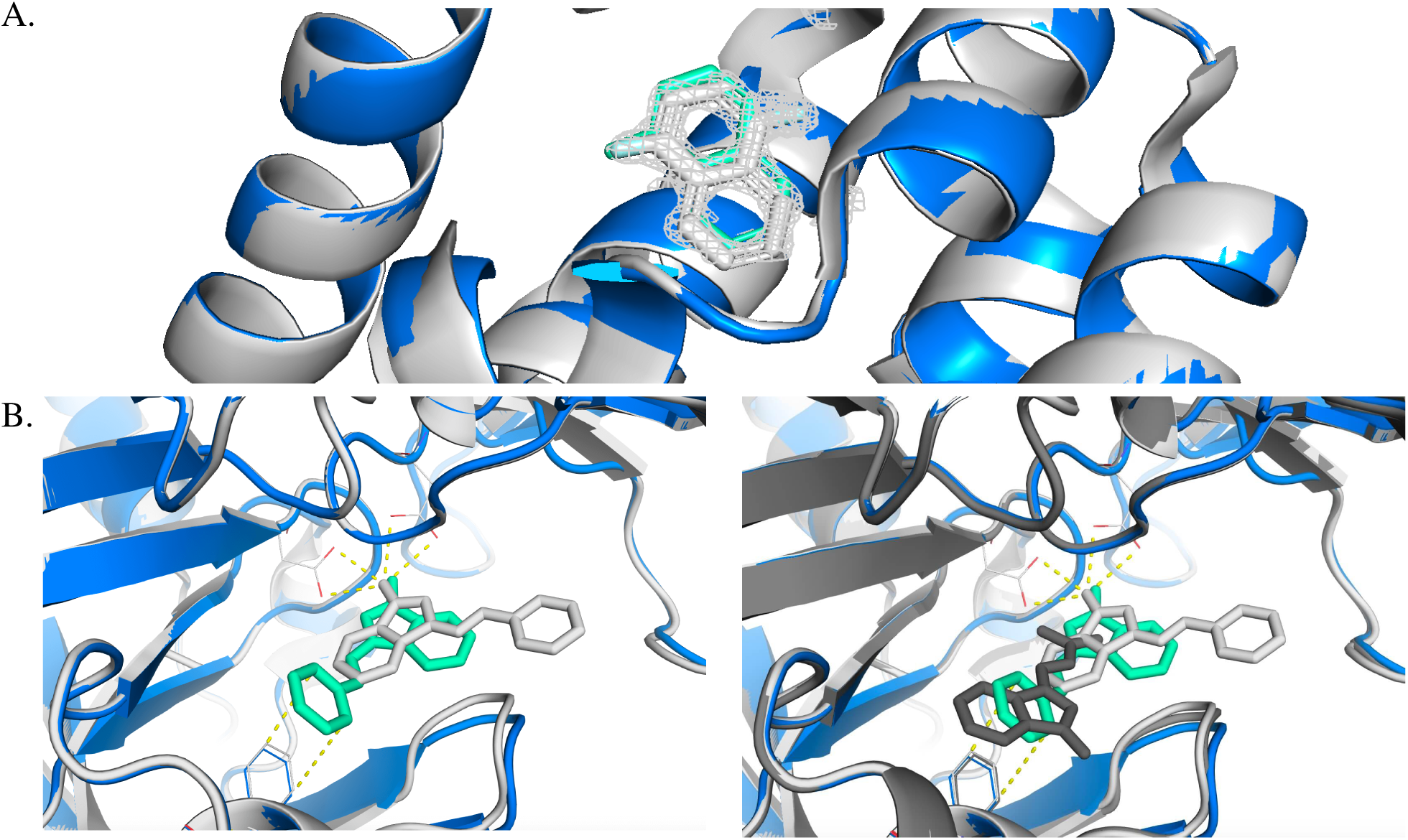
Sensible failure cases. Predicted protein chains are shown in blue, predicted ligands show in turquoise, and the ground truth is shown in gray. **A.** T4 lysozyme mutant L99A in complex with 1-fluoranylnaphthalene alternative ligand conformations (PDB: 7LOE). **B.** Endothiapepsin in complex with compound FU5-1 (PDB: 5SAK) and similar fragment in the PDB for the same target (PDB: 5SAN).

### 2.3 Proteins

We benchmark our model against AF2.3, the top-performing open-source model available for comparison. Both models predict the first bioassembly of each structure. For AF2.3, non-protein chains are excluded, and non-standard residues are mapped to standard ones, as the model does not support these components. Our model predicts the complete complex, though we do not assess whether including additional chains in the input influences performance. The results are summarized in Figure 1[B]. We report the DockQ success rate for different types of interfaces: all protein-protein interfaces and protein-antibody interfaces. We report results from our model with 10 recycles and from AF2.3(5×5) with 20 recycles, including separate scores for best (oracle), median, and top-1 ranked predictions. For *Protenix*, the ranker used for protein interfaces is the “chain pair ipTM” confidence. Overall, our model achieves higher DockQ success rate compared to AF2.3, indicating improved prediction accuracy. The only exception occurs in protein-antibody cases, where our top-ranked predictions are on par with those of AF2.3, indicating potential for improvement in the sample ranker.

**Case study.** Beyond the performance metrics, we also identify an interesting case, as shown in Figure 5. It highlights the potential applicability of *Protenix* as a tool for biology and pharmacology research, helping with hypothesis testing and uncovering novel mechanisms which can serve as the starting point for the development of new therapeutics.

**Figure 5:**
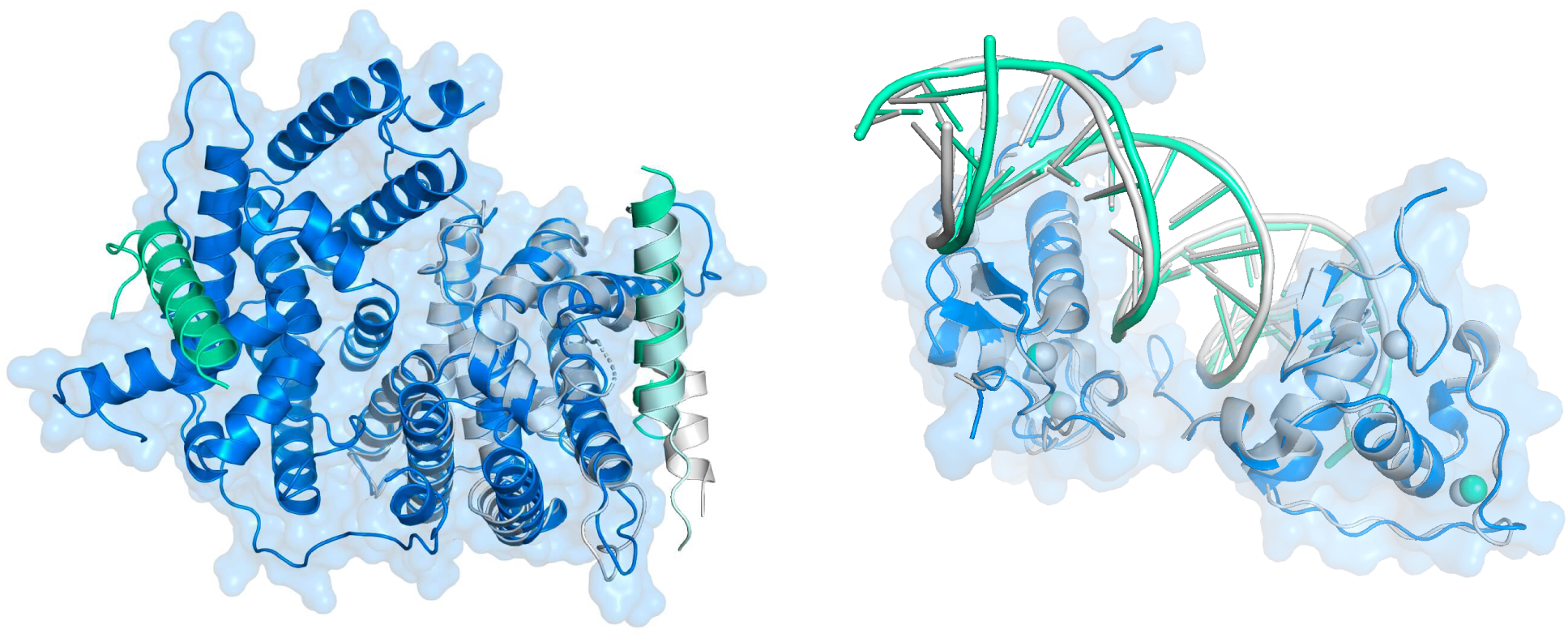
Potential applications. **Left**: *Protenix* prediction (blue and turquoise) on a novel protein-protein interaction from PDB: 7XVA (gray), only very recently characterized as novel regulation mechanism for the TR4 transcription factor binding to JAZF1, a regulation mechanism involved in different cancer types, whose understanding might lead to novel therapeutic developments. **Right**: Notably, *Protenix* also predicts correctly the TR4 DBD binding to DNA as homodimer from PDB: 7XV8, highlighting integrative modeling capabilities and ability to model complex biological mechanisms [19].

### 2.4 Nucleic Acids

We extend our evaluations to include RNA and DNA targets, where *Protenix* demonstrates performance comparable to AF3 while achieving higher accuracy than RF2NA. *Protenix* does not use MSA for nucleic chains.

**CASP15.** We assess *Protenix* on CASP15 RNA targets, following the approach in the AF3 paper, focusing on eight targets that were publicly available as of December 1, 2023. We compare the results of *Protenix*, AF3, AIchemy_RNA2, and RF2NA. The AIchemy_RNA2 predicted structures are obtained from the CASP website, with those exhibiting the highest LDDT scores selected for analysis. The RF2NA predicted structures are downloaded from the Zenodo repository [20], alongside the original paper. The metrics for AF3 are taken from the preprint by Bernard et al. [21], where the authors systematically evaluated AF3’s RNA prediction performance and reported LDDT and TM-score values computed using OpenStructure [22, 23]. Since the authors evaluted AF3’s performance based on a single seed, we also report the performance of *Protenix* using a single seed instead of five seeds. The top-ranked sample of each structure, selected based on pLDDT, is used for evaluation. For RF2NA, AIchemy_RNA2, and *Protenix*, the LDDT and TM-score metrics are recomputed consistently using OpenStructure [22, 23]. As shown in Figure 1[C] and Figure 6, the average LDDT and TM-score of *Protenix* are similar to those of AF3, significantly outperforming RF2NA, but still lagging behind AIchemy_RNA2 [24], which benefits from human input.

**Figure 6:**
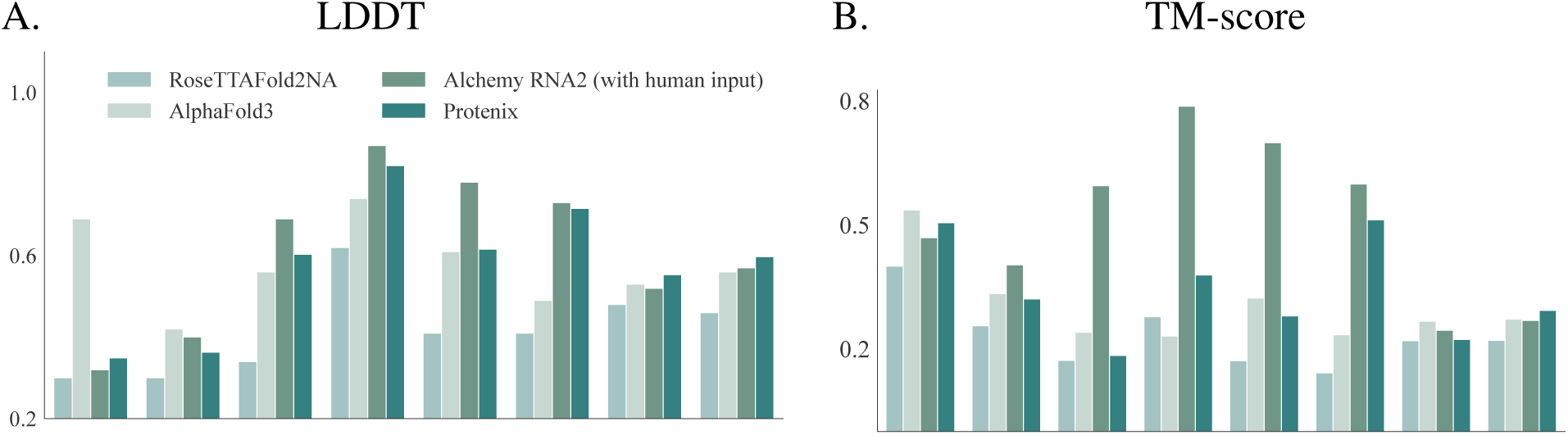
CASP15 RNA prediction accuracy. The results from AlChemy_RNA2, RF2NA, AF3, and *Protenix* are reported.

**Protein-nucleic acid complexes.** As illustrated in Figure 1[C][D], *Protenix* predicts protein-nucleic complexes with higher accuracy than RF2NA. To ensure a fair comparison, we apply the same evaluation approach to both *Protenix* and RF2NA (details in Appendix B). For protein-dsDNA interfaces, the two DNA chains in the double-stranded DNA (dsDNA) are treated as a single entity. The interface LDDT is then calculated between this entity and the protein chain. The dataset used for our comparison is curated based on the description provided in AF3 (See Appendix A.2 for details). The number of structures in our curated dataset matches the count reported in AF3’s Figure 1.c, although we cannot guarantee the datasets are identical. Notably, the performance of RF2NA differs from that reported in AF3, which may be attributed to discrepancies in the dataset or the evaluation tools used. For each structure, the top-ranked sample based on the complex ranking score (0.8 · ipTM + 0.2 · pTM − 100 · has_clash) is selected for evaluation. This is because, for dsDNA-protein interfaces, three chains are involved, making it difficult to use chain pair scores directly.

**Case study.** We evaluate *Protenix*’s ability to predict protein-nucleic acid complexes, with a particular focus on cases with novel structures. One such example is shown in Figure 7, where *Protenix* successfully predicts a novel protein-DNA complex that reveals the mechanism by which a monomeric repressor promotes transcriptional silencing.

**Figure 7:**
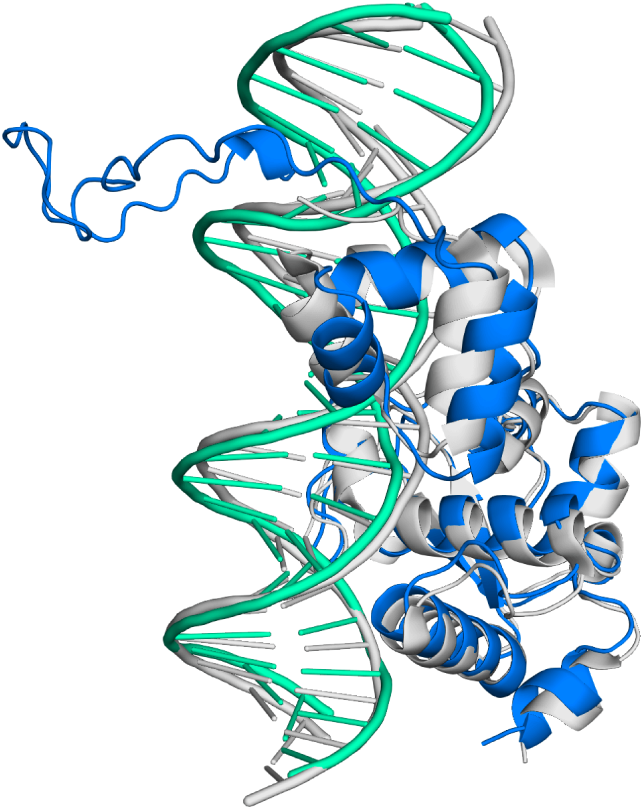
Novel protein-nucleic acid complex. Accurate prediction (blue and turquoise) by *Protenix* on a novel protein-nucleic acid complex (mycobacteriophage immunity repressor protein bound to double stranded DNA) without close homologs in the training set. (PDB: 7R6R (gray))

### 2.5 Discussion

**Confidence score.** We make slight modifications to the confidence head architecture in contrast to AF3, which will be discussed in Section 3. After training the confidence head, we observe that predicted scores can be used to select interfaces on low homology recentPDB dataset. As illustrated in Figure 8, a positive Pearson correlation (*ρ*>0.5) is observed between predicted scores and interface DockQ scores for diverse interface types, including protein-protein and protein-nucleic acid interactions. Given that our PAE head has been trained for only 4K steps, there is still huge potential for further improvement.

**Figure 8:**
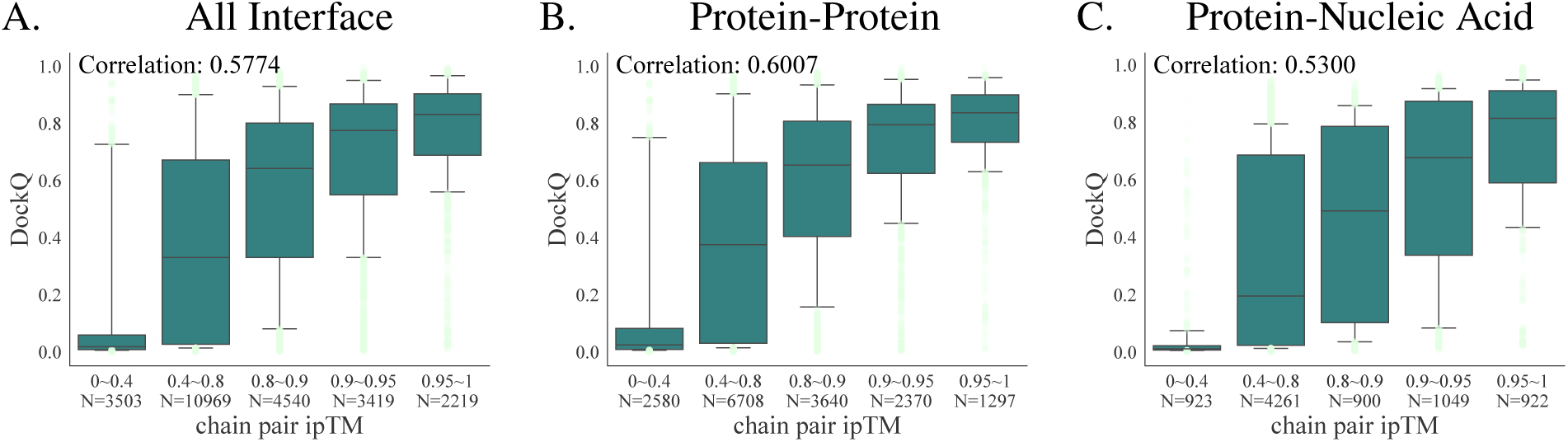
Relationship between predicted chain pair ipTM and the DockQ score. **A**. A box-and-whisker plot of DockQ against binned chain pair ipTM on both protein-protein and protein-nucleic acid interfaces. **B.** The subset of figure **A** on the protein-protein interfaces. **C**. The protein-nucleic acid subset of figure A.

**Limitations.** We review the showcase examples presented in the AF3 paper to evaluate how our model performs on these cases. We find that our model also predicts these structures with high accuracy. Upon observing similar results, we further investigated the training set and noted that some of these complexes share significant similarities with certain training samples. For instance, as shown in Figure 9, the protein-DNA complex 7PZB has a close counterpart in the training set (3MZH), as does the protein-ligand complex 7XFA, which is similar to 5OAX. This suggests that the accuracy of both *Protenix* and AF3 may partially result from memorization, indicating the need for more out-of-distribution (OOD) test sets to better assess its generalizability.

**Figure 9:**
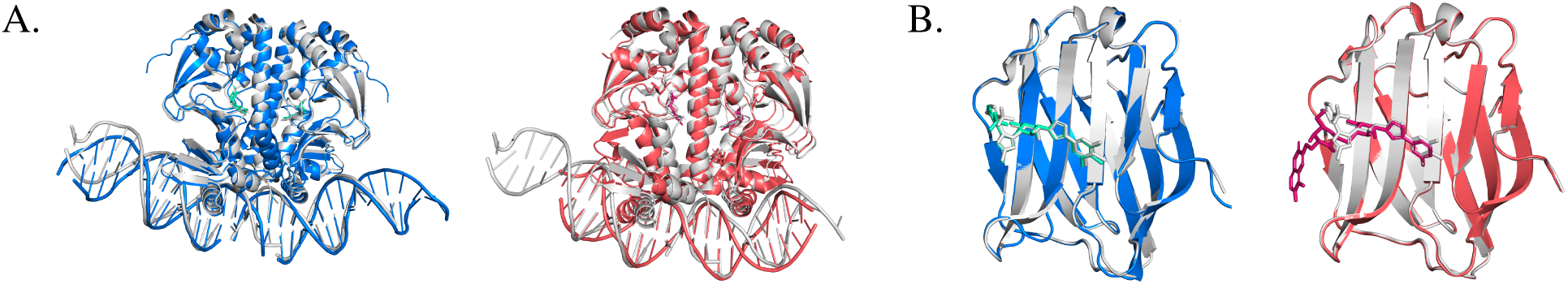
**A. Left**: Example of good prediction of protein-nucleic acids interactions by *Protenix* (blue) and PDB crystal structure 7PZB (gray). **Right**: Close sample in the training set (3MZH) in salmon. **B. Left**: Example of good prediction of protein-ligand interactions by *Protenix* in PoseBusters and PDB crystal structure (7XFA). **Right**: Close sample in the training set (5OAX).

## 3 Methods

### 3.1 Data Pipeline

Inclusion of various biomolecules creates exceptional challenges for data curation and featurization. We dive into the details of AF3 supplementary materials to reproduce the data pipeline. Below we list the major differences in our implementation.

**Parser.** In selecting alternative locations, we use the first occupancy rather than the largest one. This is because we found that using the largest occupancy may result in some adjacent residues adopting different conformations, preventing the formation of covalent bonds and leading to chain breaks.

**MSA.** We search MSAs using MMSEQS2 [25] and ColabFold [7] MSA pipeline, and use MSAs from the Uniref100 [26] database for pairing (species are identified with taxonomy IDs). We do not use MSA for nucleic chains.

**Templates.** We do not use templates.

**Featurization.**

- **Reference features.** We use the version of CCD [27] downloaded on 2024-06-08 and generate RDKit conformers using pdbeccdutils [28] v0.8.5. When RDKit conformer generation fails, we use the CCD ideal coordinates and do not use the representative coordinates to avoid potential data leakage.
- **Bond features.** Bond features between tokens only include bonds within ligands, bonds between ligands and polymers, and bonds within non-standard residues.

**Cropping.** Our implementation of the cropping method takes the following aspects into consideration.

- In our contiguous cropping method, we discard metals and ions, as this approach cannot guarantee that cropped atoms remain close to each other. When metals or ions are cropped in isolation from other chains, it becomes challenging for the model to accurately predict their positions, potentially leading to large training losses. Therefore, we exclude metals and ions from contiguous cropping but retain them in spatial cropping.
- Since ligands and non-standard amino acids are represented by a single atom per token, they may be cropped into fragments. To prevent this, we ensure that ligands and non-standard amino acids remain intact and are not split into incomplete segments during cropping.
- We account for the impact of cropping on chain permutation during training. For spatial cropping, we restrict permutation to chains within the cropped region to ensure predictions focus on spatially close structures. In contrast, for contiguous cropping, where cropped chains may be distant but equivalent chains may be closer to each other, we allow equivalent permutations and adjust the cropped region based on model predictions. Details on permutation handling are provided in Appendix C.

We believe this is a typo because the per-sample weighting of the diffusion loss, which is different from the original form in AF3, is carefully designed in a principled way by the authors of EDM [29].

### 3.2 Model and Training

We refer readers to the supplementary materials of AF3 for detailed methodology, as our implementation is based on it. In this section, we primarily highlight the differences we have introduced in our implementation and explain the underlying reasons for making these modifications.

**Corrections and adjustments.** In our analysis of the algorithms presented in the original paper, we identify several errors and ambiguities. Some of the adjustments are key to performance. These corrections and their corresponding explanations are shown in Table 1. Additionally, we make slight modifications to the confidence head by incorporating LayerNorm and adding a few linear layers (see our released code). We find that the confidence loss does not converge as effectively when implemented exactly as described in the original paper.

**Table 1:**
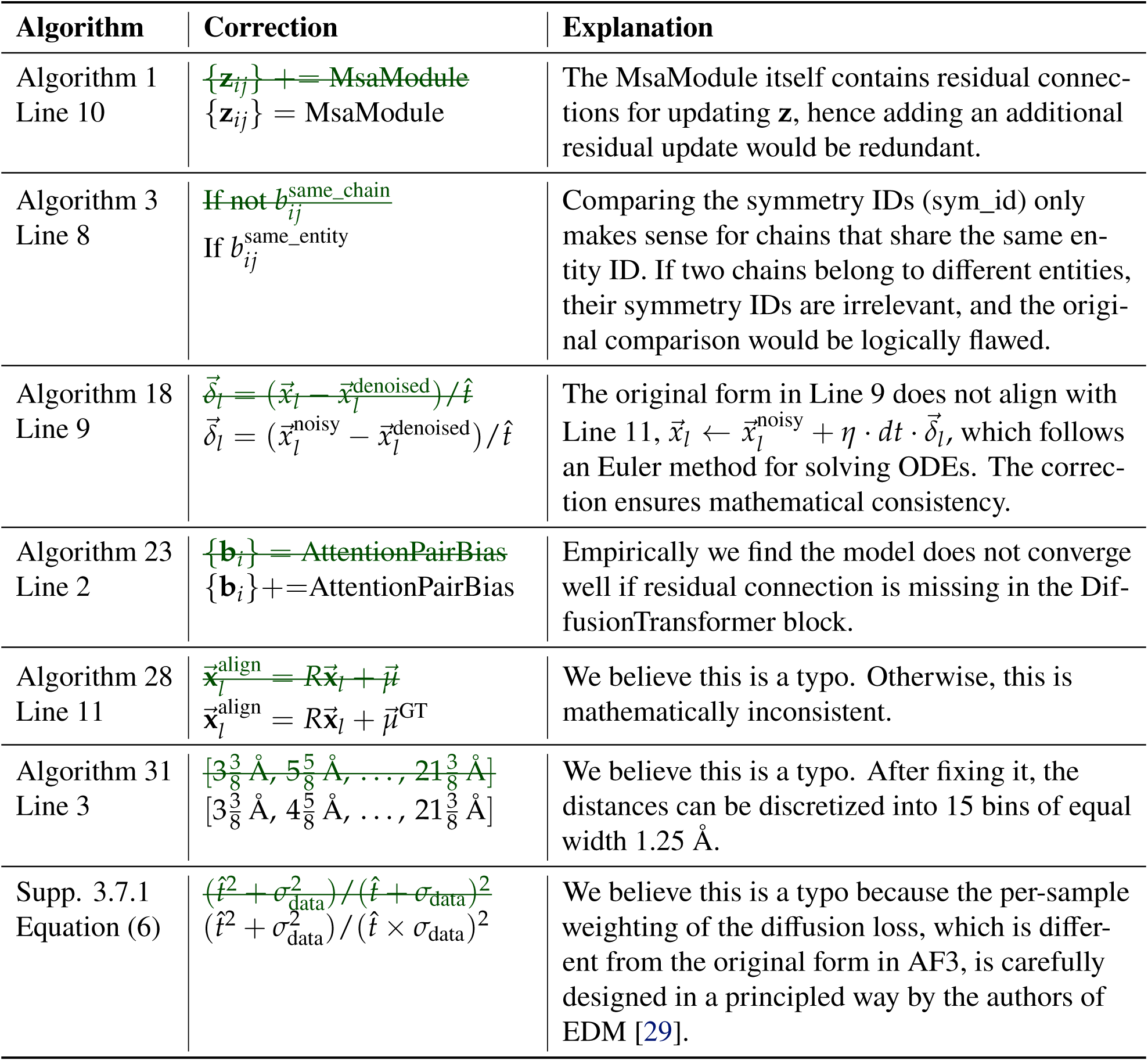
Corrections. We provide a description and explanation of the introduced corrections in our implementation. The algorithm numbers correspond to those in the AF3 Supplementary materials.

**Parameter initialization.** We do not conduct an extensive analysis of the impact of different initialization strategies. However, we apply zero-initialization to several modules. Specifically, some linear layers are zero-initialized to ensure that each residual layer behaves as an identity operation at initialization. Following recent work on conditional diffusion models [30, 31], we zero-initialize two linear layers in the AdaptiveLayerNorm. Additionally, we apply zero initialization to the N-cycle loop connection blocks in the Pairformer (see Algorithm 1, line 8 in AF3 report). Empirically, we find that these zero-initializations help prevent dimensional collapse in the network weights and mitigate the explosion of hidden values.

**Scalability.** Despite having only about 386 million parameters, AF3 requires an enormous amount of computation. Especially, its Pairformer blocks lead to numerous intermediate activations. Additionally, during training, the diffusion module has a large diffusion batch size and applies activations for each atom pair, thereby creating memory bottlenecks for both training and inference. To tackle these challenges and enhance training efficiency, we have introduced several approaches as follows:

- **BF16 mixed precision training.** Compared to FP32 training, BF16 training significantly reduces peak memory usage and nearly doubles the actual training speed. However, not all GPU architectures support BF16, with compatibility limited to specific models like NVIDIA’s Ampere [32] and Hopper [33].
- **Custom CUDA Kernels.** A naive implementation of LayerNorm in PyTorch [34] often encounters memory limitations. Building on optimizations from FastFold [35] and OneFlow [36], we develop a custom LayerNorm implementation that adaptively adjusts block and grid sizes while leveraging shared memory to optimize GPU utilization. This approach yields a 30%-50% improvement in end-to-end training speed compared to PyTorch’s native LayerNorm across various training stages.
- **DS4Sci_EvoformerAttention.** By utilizing the DS4Sci_EvoformerAttention kernels from Deep-Speed4Science [37], a 10%-20% improvement in end-to-end training speed across various training stages is achieved.
- **Other Considerations.** A direct implementation of the model update steps, as outlined in the AF3 paper, leads to the creation of unnecessarily large tensors. To avoid this, we use alternative implementations that prevent the formation of these large tensors. Additionally, to address memory bottlenecks during both training and inference, we employ several optimization techniques, including gradient checkpointing, in-place operations, tensor offloading, and chunking, similar to the approach used in OpenFold [10].

**Training.** *Protenix* is trained on 192 GPUs over approximately two weeks. Table 2 provides a summary of our training stages. We also simplified the hyperparameters considerably, as our training utilizes the Protein Monomer Distillation dataset only, excluding other distillation sets. Although we implement a multi-stage training setup similar to AF3, our model is trained with significantly fewer steps. We train the model for 75K steps during the initial stage and 15K steps during fine-tuning stage 1. We run only 4K steps for fine-tuning stage 2, which may be insufficient for adequately training the confidence head.

**Table 2:** Training Stages Configurations. We list the key hyper-parameters about the multiple training stages.

| Details | Initial Training | Fine-tuning Stage 1 | Fine-tuning Stage 2 |
| --- | --- | --- | --- |
| Training Crop Size | 384 | 640 | 640 |
| Train PAE Head | ✗ | ✗ | ✓ |
| Train Structure and Distogram | ✓ | ✓ | ✗ |
| Diffusion Batch Size | 48 | 32 | - |
| Train on Distillation Sets | ✓ | ✓ | ✗ |
| Bond Loss Weight | 0 | 1 | - |

## 4 Accessibility

Source code and datasets are available at the Protenix repository^*^. We encourage readers to refer to the source code for detailed use cases.

**Data release.** Our data release aims to provide a comprehensive foundation for researchers to reproduce results, conduct further analysis, or leverage the datasets for new applications. We highlight our contributions as follows:

- **Significant computational efforts**: The processed data includes time-consuming MSA (multiple sequence alignment) search results, covering all instances in the wwPDB. This is especially beneficial for researchers with limited access to computing resources, saving them considerable time and effort in data processing.
- **PDB ID to processed data mapping**: To facilitate ease of use, we curate the dataset and provide a mapping from Protein Data Bank (PDB) IDs to the processed data format. This allows users to quickly integrate the data into their pipelines without extensive adjustments, streamlining their research process.

**Code release.** Our code release is designed to maximize usability and encourage experimentation, offering researchers a robust set of tools for extending our work:

- **Inference code**: We provide code for model inference, along with a notebook that allows users to customize and model their inputs.
- **Trainable code**: We release trainable code, allowing users to retrain or fine-tune models on additional datasets. This is beneficial for those interested in adapting the model to other applications or improving model performance on specific tasks.
- **Documentation**: Comprehensive documentation is included in our codebase to guide users through setting up, running, and modifying the code, making it easy for users at all skill levels to get started.

## 5 Future Plan

We believe the first version of *Protenix* delivers promising results, but there is still considerable room for improvement. Moving forward, we plan to continue enhancing the model and introduce additional features and functionalities for users.

- **Next improved version.** Continuously improving the model’s performance will remain a priority on our to-do list.
- **Extra features.** Certain features may be particularly useful for specific applications, such as predicting structures with incorporated priors on partial structures or interactions, or making predictions without utilizing the MSA feature.
- **Evaluation and benchmark.** In our reproduction process, we observe that the evaluation procedure is highly complex, demanding close attention to numerous details, particularly given the diversity of molecular types involved. We plan to continue enhancing our evaluation tools and may eventually open-source a general-purpose evaluation platform to support fair model comparisons.

## Supporting information

figures and result summary

## Acknowledgements and Thanks

We would like to express our sincere gratitude to the authors and contributors of the open-source tools and datasets. Their efforts in making such valuable resources publicly available have been instrumental in advancing our work. In particular, we thank the developers of AlphaFold, OpenFold, and ColabFold, whose contributions provided a strong foundation for our study.

### A Dataset

#### A.1 Training Dataset

In addition to WeightedPDB, we also use AlphaFold 2 predictions to train our model. We use representative sequences from MGnify [38] with greater than 200 residues as the distillation data source. Due to resource limitations, we run 5 models on a small part of the data and only 1 model (model-3) on the rest. As a supplement, we also use a subset of the OpenProteinSet [39]. All distillation datasets are filtered based on pLDDT to obtain high-quality training data.

**Table 3:** Datasets used to train *Protenix*. We list the descriptions for the training datsets.

| Name | Description | Sampl. strategy | Weight |
| --- | --- | --- | --- |
| Weighted PDB | Ground truth PDB structures | weighted | 0.60 |
| Protein monomer distillation (Set1) | Protein monomer predictions from MGnify using 1 model | uniform | 0.35 |
| Protein monomer distillation (Set2) | Protein monomer predictions from MGnify using 5 models | uniform | 0.03 |
| OpenProteinSet [39] subset | Protein monomer predictions from Uniclust30 | uniform | 0.02 |

#### A.2 Evaluation Dataset

##### A.2.1 PoseBusters V2

We use the PoseBusters Benchmark Set Verison 2 (PoseBusters V2) [18] for evaluating protein-ligand interactions. This dataset contains 308 structures. To facilitate a fair comparison with AF3, we follow their procedure [4] to remove chains clashing with the ligand of interest (specifically, files: 7O1T, 7PUV, 7SCW, 7WJB, 7ZXV, 8AIE). Similarly, for PDB entry 8F4J, we retain only the protein chains within 20 Å of the ligand. We follow the AF3 practice, in which the model inference is performed on the *asymmetric unit* of each structure, to allow for direct comparability with both the numbers reported in AF3 and their released predictions.

##### A.2.2 Low Homology Recent PDB Set

We follow the Supplementary Information of AF3 to curate the Low Homology (LowH) Recent PDB set for evaluating the prediction of protein and nucleic acid interfaces. This set consists of PDB targets released between 2022-05-01 and 2023-01-12, filtered for sequence similarity below 40% to better assess model generalization. We apply the same token cutoff to evaluate models on structures with fewer than 2560 tokens. However, due to some missing details in the paper and various possible differences related to clustering, low homology filtering, and other factors, our dataset still differs in PDB and cluster numbers compared to the AF3 paper, despite our efforts with different tools and settings. The number of clusters used for each evaluation task is specified in the figures.

##### A.2.3 Nucleic Acids Evaluation Set

**CASP15.** Eight targets that were publicly available as of December 1, 2023 are used for the evaluation: R1116/8S95, R1117/8FZA, R1126 (https://predictioncenter.org/casp15/TARGETS_PDB/R1126.pdb), R1128/8BTZ, R1136/7ZJ4, R1138/[7PTK/7PTL], R1189/7YR7 and R1190/7YR6.

**Protein-Nucleic Acids Complexes.** The dataset used for the results in Figure 1 is curated by the following procedure. The nucleic acid evaluation set is screened from the RecentPDB dataset’s PDB IDs provided in Table 14 of the AF3 supplementary materials. RoseTTAFold2NA can only input up to 1000 residues, hence the dataset does not include systems with a total residue count exceeding 1000. Additionally, it does not contain systems with both DNA and RNA present simultaneously. Subsequently, we retain structures with only one RNA chain and one protein chain, or with two DNA chains and one protein chain. We then remove all systems containing non-standard residues in the DNA. The remaining structures coincided with the number in AF3 Figure 1, 25 RNA-Protein structures and 38 dsDNA-Protein structures. Although the number here matches that in AF3, it does not necessarily mean the evaluation was conducted on an identical evaluation set, which may lead to discrepancies in the metrics.

### B Evaluation Workflow

For each PDB entry, we perform the following steps:

1. **Inference and Sampling.**
2. Unless specified, we generate 25 samples using 5 model seeds. Each sample is exported as both a CIF file and a JSON file, with the JSON file containing confidence ranking scores. For targets in PoseBusters V2, we predict the asymmetric unit of the structure; for targets in all other datasets, we predict the first bioassembly of the structure. No cropping is applied during inference, and we generally use 10 recycles unless otherwise noted. AF3 specifies its configuration for recycles only in Appendix 5.10, where inference times are reported using 10 trunk recycles on 16 GPUs. Therefore, we infer that AF3 likely used at least 10 recycles in their report.
3. **Chain and Atom Permutation.** For each predicted structure (CIF), we perform chain mapping to establish a one-to-one correspondence between predicted and true entities. Once equivalent chains are identified, we permute the chains and atoms to better align the prediction with the true structure (details in Appendix C).
4. **Metrics Computation.** Different types of interactions are evaluated with specific methods. For proteins, we use the DockQ Python package to calculate DockQ scores. For predictions on PoseBusters V2, we follow these steps:

a. *Exporting Structures*: The ligand of interest is exported to an SDF file, while all other chains with at least one atom within 5 Å of any atom in the predicted ligand are exported to a PDB file.
b. *PoseBusters Analysis*: Finally, we use the PoseBusters Python package v.0.2.7 to calculate the RMSD of the ligands and evaluate any structural violations using the exported files.

### C Chain and Atom Permutation

#### Algorithm 1 Chain Permutation (Training)

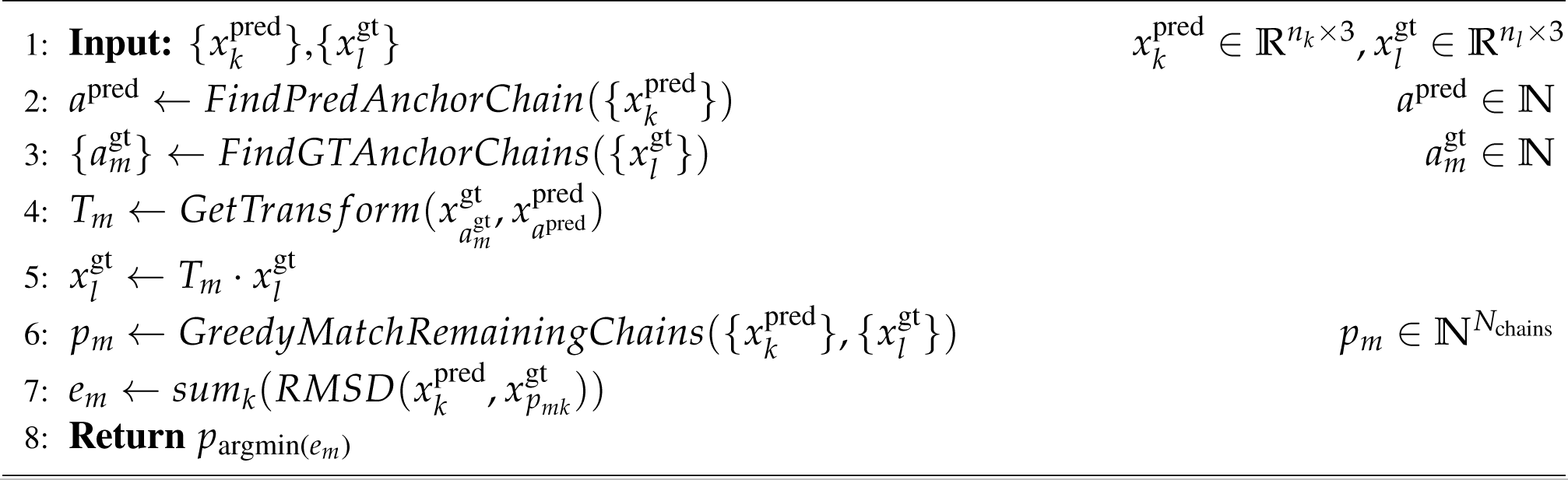

#### Algorithm 2 Chain Permutation (Inference)

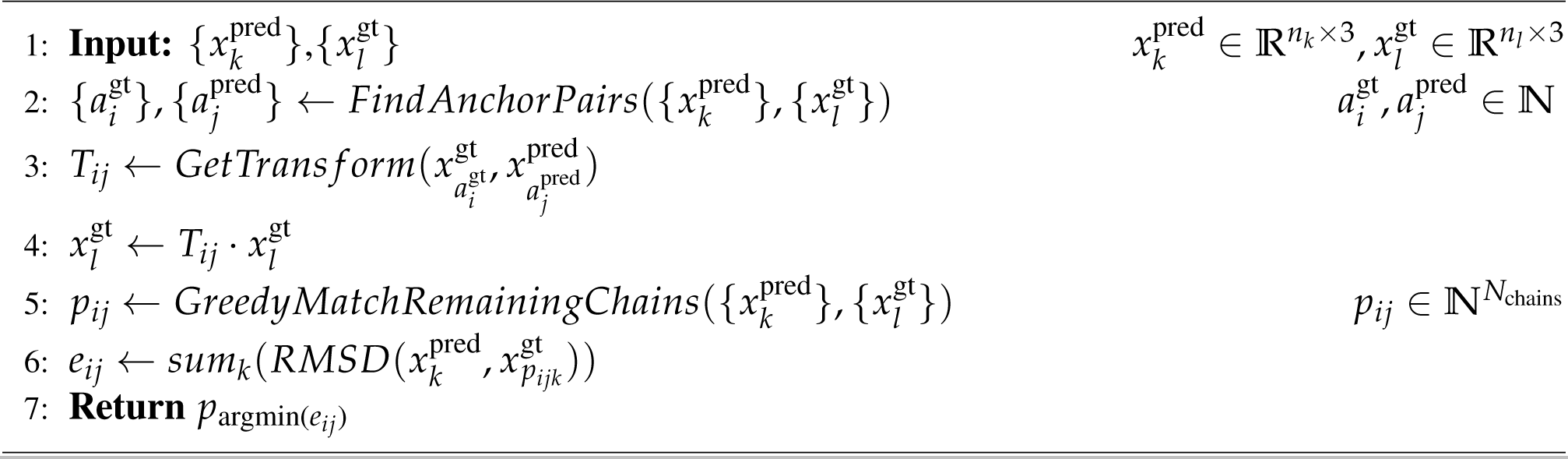

To address ambiguities in chain and atom naming for accurate loss and metric calculations, we introduce chain and atom permutations at various stages. The methods are detailed below.

**Training.** During training, we permute the chains and atoms of the ground truth structure to align with the mini-rollout predictions before the diffusion sampling stage. Our chain permutation approach largely builds on the heuristic algorithms used in AF3 [4] and AF2.3 [40]. Initially, anchor chains in the ground truth and predicted structures are identified for alignment. This is followed by a greedy matching process for the remaining chains (Algorithm 1, Line 6). The permutation that minimizes the average RMSD between the prediction and ground truth chains is selected. For better efficiency and applicability, we also introduce several adaptations: we only consider entities with at least four resolved tokens for anchor chains and prioritize polymer chains to reduce permutation complexity. Additionally, to account for cropping during training, we first identify an anchor chain in the predicted structure, then enumerate candidate anchor chains in the ground truth (Algorithm 1, Line 2 and 3). Specifically, for samples using spatial cropping, we restrict the ground truth anchor chain to chains included in the cropped region, as this cropping method may exclude certain chains during training. In contrast, for contiguous cropping, we lift this restriction, as all chains, even if cropped, remain included throughout the process.

We then perform atom permutation on the ground-truth structure within each residue/ligand to correct atom-level computation. This process begin by first aligning the global structure of the prediction to that of the ground truth, followed by residue-level permutation to minimize within-residue RMSD.

**Inference.** During inference, we permute the chains and atoms in the structure predicted by the diffusion module, aligning them to the ground truth independently for each diffusion sample. The chain permutation approach here differs slightly from the training stage. First, we identify anchor chains in both the predicted and ground truth structures, requiring a minimum of four resolved tokens (Algorithm 2, Line 2). We then enumerate all anchor chain pairs between the prediction and ground truth and greedily matched the remaining chains for each pair, selecting the permutation which minimizes the global RMSD.

When applying chain permutation to ligand complexes (e.g., PoseBusters V2), we generate all possible combinations of pockets and ligands, and then select the permutation resulted in the lowest ligand-specific RMSD. For atom permutation, we align the predicted structure to the ground truth based specifically on the coordinates of pockets and ligands, rather than using global coordinates.

## Footnotes

* https://github.com/bytedance/Protenix

