## Supplementary material for "Protenix - Advancing Structure Prediction Through a Comprehensive AlphaFold3 Reproduction": figures and result summary: bibfile.bib

@article{jumper2021af2,
  title={Highly accurate protein structure prediction with AlphaFold},
  author={Jumper, John and Evans, Richard and Pritzel, Alexander and Green, Tim and Figurnov, Michael and Ronneberger, Olaf and Tunyasuvunakool, Kathryn and Bates, Russ and {\v{Z}}{\'\i}dek, Augustin and Potapenko, Anna and others},
  journal={nature},
  volume={596},
  number={7873},
  pages={583--589},
  year={2021},
  publisher={Nature Publishing Group}
}

@article{lin2023esmfold,
  title={Evolutionary-scale prediction of atomic-level protein structure with a language model},
  author={Lin, Zeming and Akin, Halil and Rao, Roshan and Hie, Brian and Zhu, Zhongkai and Lu, Wenting and Smetanin, Nikita and Verkuil, Robert and Kabeli, Ori and Shmueli, Yaniv and others},
  journal={Science},
  volume={379},
  number={6637},
  pages={1123--1130},
  year={2023},
  publisher={American Association for the Advancement of Science}
}

@article{baek2021rosettafold,
  title={Accurate prediction of protein structures and interactions using a three-track neural network},
  author={Baek, Minkyung and DiMaio, Frank and Anishchenko, Ivan and Dauparas, Justas and Ovchinnikov, Sergey and Lee, Gyu Rie and Wang, Jue and Cong, Qian and Kinch, Lisa N and Schaeffer, R Dustin and others},
  journal={Science},
  volume={373},
  number={6557},
  pages={871--876},
  year={2021},
  publisher={American Association for the Advancement of Science}
}

@article{ahdritz2024openfold,
  title={OpenFold: Retraining AlphaFold2 yields new insights into its learning mechanisms and capacity for generalization},
  author={Ahdritz, Gustaf and Bouatta, Nazim and Floristean, Christina and Kadyan, Sachin and Xia, Qinghui and Gerecke, William and O’Donnell, Timothy J and Berenberg, Daniel and Fisk, Ian and Zanichelli, Niccol{\`o} and others},
  journal={Nature Methods},
  pages={1--11},
  year={2024},
  publisher={Nature Publishing Group US New York}
}

@article{mirdita2022colabfold,
  title={ColabFold: making protein folding accessible to all},
  author={Mirdita, Milot and Sch{\"u}tze, Konstantin and Moriwaki, Yoshitaka and Heo, Lim and Ovchinnikov, Sergey and Steinegger, Martin},
  journal={Nature methods},
  volume={19},
  number={6},
  pages={679--682},
  year={2022},
  publisher={Nature Publishing Group US New York}
}


@article{senior2020alphafold,
  title={Improved protein structure prediction using potentials from deep learning},
  author={Senior, Andrew W and Evans, Richard and Jumper, John and Kirkpatrick, James and Sifre, Laurent and Green, Tim and Qin, Chongli and {\v{Z}}{\'\i}dek, Augustin and Nelson, Alexander WR and Bridgland, Alex and others},
  journal={Nature},
  volume={577},
  number={7792},
  pages={706--710},
  year={2020},
  publisher={Nature Publishing Group UK London}
}

@article{evans2021afmultier,
  title={Protein complex prediction with AlphaFold-Multimer},
  author={Evans, Richard and O’Neill, Michael and Pritzel, Alexander and Antropova, Natasha and Senior, Andrew and Green, Tim and {\v{Z}}{\'\i}dek, Augustin and Bates, Russ and Blackwell, Sam and Yim, Jason and others},
  journal={biorxiv},
  pages={2021--10},
  year={2021},
  publisher={Cold Spring Harbor Laboratory}
}

@misc{ligobiosciences_github,
  author       = {Ligo Biosciences},
  title        = {GitHub repository -  AlphaFold3 -Ligo Biosciences},
  year         = {2024},
  url          = {https://github.com/Ligo-Biosciences/AlphaFold3/tree/main},
  note         = {Accessed: 2024-10-25}
}
@misc{lucidrians_github,
  author       = {},
  title        = {GitHub repository - Implementation of Alphafold 3 in Pytorch - lucidrains},
  year         = {2024},
  url          = {https://github.com/lucidrains/alphafold3-pytorch/tree/main
},
  note         = {Accessed: 2024-10-25}
}
@article{abramson2024af3,
  title={Accurate structure prediction of biomolecular interactions with AlphaFold 3},
  author={Abramson, Josh and Adler, Jonas and Dunger, Jack and Evans, Richard and Green, Tim and Pritzel, Alexander and Ronneberger, Olaf and Willmore, Lindsay and Ballard, Andrew J and Bambrick, Joshua and others},
  journal={Nature},
  pages={1--3},
  year={2024},
  publisher={Nature Publishing Group UK London}
}


@article{bertoline2023overview,
  title={Before and after AlphaFold2: An overview of protein structure prediction},
  author={Bertoline, Let{\'\i}cia MF and Lima, Ang{\'e}lica N and Krieger, Jose E and Teixeira, Samantha K},
  journal={Frontiers in bioinformatics},
  volume={3},
  pages={1120370},
  year={2023},
  publisher={Frontiers Media SA}
}


@article{richardson2023mgnify,
  title={MGnify: the microbiome sequence data analysis resource in 2023},
  author={Richardson, Lorna and Allen, Ben and Baldi, Germana and Beracochea, Martin and Bileschi, Maxwell L and Burdett, Tony and Burgin, Josephine and Caballero-P{\'e}rez, Juan and Cochrane, Guy and Colwell, Lucy J and others},
  journal={Nucleic Acids Research},
  volume={51},
  number={D1},
  pages={D753--D759},
  year={2023},
  publisher={Oxford University Press}
}


@misc{harris2022nvidiahopper,
  title={NVIDIA Hopper GPU Architecture Accelerates Dynamic Programming Up to 40x Using New DPX Instructions},
  author={Harris, Dion},
  year={2022}
}

@article{krashinsky2020nvidiaampere,
  title={Nvidia ampere architecture in-depth},
  author={Krashinsky, Ronny and Giroux, Olivier and Jones, Stephen and Stam, Nick and Ramaswamy, Sridhar},
  journal={NVIDIA blog: https://devblogs. nvidia. com/nvidia-ampere-architecture-in-depth},
  year={2020}
}

@inproceedings{esser2024sd3,
  title={Scaling rectified flow transformers for high-resolution image synthesis},
  author={Esser, Patrick and Kulal, Sumith and Blattmann, Andreas and Entezari, Rahim and M{\"u}ller, Jonas and Saini, Harry and Levi, Yam and Lorenz, Dominik and Sauer, Axel and Boesel, Frederic and others},
  booktitle={Forty-first International Conference on Machine Learning},
  year={2024}
}


@inproceedings{peebles2023dit,
  title={Scalable diffusion models with transformers},
  author={Peebles, William and Xie, Saining},
  booktitle={Proceedings of the IEEE/CVF International Conference on Computer Vision},
  pages={4195--4205},
  year={2023}
}

@article{karras2022edm,
  title={Elucidating the design space of diffusion-based generative models},
  author={Karras, Tero and Aittala, Miika and Aila, Timo and Laine, Samuli},
  journal={Advances in neural information processing systems},
  volume={35},
  pages={26565--26577},
  year={2022}
}

@article{paszke2019pytorch,
  title={Pytorch: An imperative style, high-performance deep learning library},
  author={Paszke, Adam and Gross, Sam and Massa, Francisco and Lerer, Adam and Bradbury, James and Chanan, Gregory and Killeen, Trevor and Lin, Zeming and Gimelshein, Natalia and Antiga, Luca and others},
  journal={Advances in neural information processing systems},
  volume={32},
  year={2019}
}

@article{yin2024evalantibody,
  title={Evaluation of AlphaFold antibody--antigen modeling with implications for improving predictive accuracy},
  author={Yin, Rui and Pierce, Brian G},
  journal={Protein Science},
  volume={33},
  number={1},
  pages={e4865},
  year={2024},
  publisher={Wiley Online Library}
}


@article{kingma2014adam,
  title={Adam: A method for stochastic optimization},
  author={Kingma, Diederik P},
  journal={arXiv preprint arXiv:1412.6980},
  year={2014}
}

@article{buttenschoen2024posebusters,
  title={PoseBusters: AI-based docking methods fail to generate physically valid poses or generalise to novel sequences},
  author={Buttenschoen, Martin and Morris, Garrett M and Deane, Charlotte M},
  journal={Chemical Science},
  volume={15},
  number={9},
  pages={3130--3139},
  year={2024},
  publisher={Royal Society of Chemistry}
}

@article{berman2000pdb,
  title={The protein data bank},
  author={Berman, Helen M and Westbrook, John and Feng, Zukang and Gilliland, Gary and Bhat, Talapady N and Weissig, Helge and Shindyalov, Ilya N and Bourne, Philip E},
  journal={Nucleic acids research},
  volume={28},
  number={1},
  pages={235--242},
  year={2000},
  publisher={Oxford University Press}
}

@article{helixfold3,
  title={Technical Report of HelixFold3 for Biomolecular Structure Prediction},
  author={PaddleHelix Team},
  journal = {arXiv},
  doi = {https://doi.org/10.48550/arXiv.2408.16975},
  year={2024}
}

@article{chai1,
	title        = {Chai-1: Decoding the molecular interactions of life},
	author       = {{Chai Discovery}},
	year         = 2024,
	journal      = {bioRxiv},
	publisher    = {Cold Spring Harbor Laboratory},
	doi          = {10.1101/2024.10.10.615955},
	url          = {https://www.biorxiv.org/content/early/2024/10/11/2024.10.10.615955},
	elocation-id = {2024.10.10.615955},
}

@article{qiao2024state,
  title={State-specific protein--ligand complex structure prediction with a multiscale deep generative model},
  author={Qiao, Zhuoran and Nie, Weili and Vahdat, Arash and Miller III, Thomas F and Anandkumar, Animashree},
  journal={Nature Machine Intelligence},
  volume={6},
  number={2},
  pages={195--208},
  year={2024},
  publisher={Nature Publishing Group UK London}
}

@article{krishna2024rosettafoldAA,
  title={Generalized biomolecular modeling and design with RoseTTAFold All-Atom},
  author={Krishna, Rohith and Wang, Jue and Ahern, Woody and Sturmfels, Pascal and Venkatesh, Preetham and Kalvet, Indrek and Lee, Gyu Rie and Morey-Burrows, Felix S and Anishchenko, Ivan and Humphreys, Ian R and others},
  journal={Science},
  volume={384},
  number={6693},
  pages={eadl2528},
  year={2024},
  publisher={American Association for the Advancement of Science}
}

@misc{nobel2024,
  title = {The Nobel Prize in Chemistry 2024},
  author = {{Nobel Prize}},
  year = {2024},
  url = {https://www.nobelprize.org/prizes/chemistry/2024/press-release/},
  note = {Accessed: 2024-10-24}
}
@article{baek2024accurate,
  title={Accurate prediction of protein--nucleic acid complexes using RoseTTAFoldNA},
  author={Baek, Minkyung and McHugh, Ryan and Anishchenko, Ivan and Jiang, Hanlun and Baker, David and DiMaio, Frank},
  journal={Nature methods},
  volume={21},
  number={1},
  pages={117--121},
  year={2024},
  publisher={Nature Publishing Group US New York}
}

@article{du2021trrosetta,
  title={The trRosetta server for fast and accurate protein structure prediction},
  author={Du, Zongyang and Su, Hong and Wang, Wenkai and Ye, Lisha and Wei, Hong and Peng, Zhenling and Anishchenko, Ivan and Baker, David and Yang, Jianyi},
  journal={Nature protocols},
  volume={16},
  number={12},
  pages={5634--5651},
  year={2021},
  publisher={Nature Publishing Group UK London}
}

@article{wu2022high,
  title={High-resolution de novo structure prediction from primary sequence},
  author={Wu, Ruidong and Ding, Fan and Wang, Rui and Shen, Rui and Zhang, Xiwen and Luo, Shitong and Su, Chenpeng and Wu, Zuofan and Xie, Qi and Berger, Bonnie and others},
  journal={BioRxiv},
  pages={2022--07},
  year={2022},
  publisher={Cold Spring Harbor Laboratory}
}

@article{bryant2024structure,
  title={Structure prediction of protein-ligand complexes from sequence information with Umol},
  author={Bryant, Patrick and Kelkar, Atharva and Guljas, Andrea and Clementi, Cecilia and No{\'e}, Frank},
  journal={Nature Communications},
  volume={15},
  number={1},
  pages={4536},
  year={2024},
  publisher={Nature Publishing Group UK London}
}


@misc{ahdritz2023openproteinset,
      title={{O}pen{P}rotein{S}et: {T}raining data for structural biology at scale}, 
      author={Gustaf Ahdritz and Nazim Bouatta and Sachin Kadyan and Lukas Jarosch and Daniel Berenberg and Ian Fisk and Andrew M. Watkins and Stephen Ra and Richard Bonneau and Mohammed AlQuraishi},
      year={2023},
      eprint={2308.05326},
      archivePrefix={arXiv},
}

@article{evans2021protein,
  title={Protein complex prediction with AlphaFold-Multimer},
  author={Evans, Richard and O’Neill, Michael and Pritzel, Alexander and Antropova, Natasha and Senior, Andrew and Green, Tim and {\v{Z}}{\'\i}dek, Augustin and Bates, Russ and Blackwell, Sam and Yim, Jason and others},
  journal={biorxiv},
  pages={2021--10},
  year={2021},
  publisher={Cold Spring Harbor Laboratory}
}

@article{liu2023structures,
  title={Structures of human TR4LBD--JAZF1 and TR4DBD--DNA complexes reveal the molecular basis of transcriptional regulation},
  author={Liu, Yunlong and Ma, Lulu and Li, Min and Tian, Zizi and Yang, Meiting and Wu, Xi and Wang, Xue and Shang, Guohui and Xie, Mengjia and Chen, Yiyun and others},
  journal={Nucleic Acids Research},
  volume={51},
  number={3},
  pages={1443--1457},
  year={2023},
  publisher={Oxford University Press}
}

@misc{cheng2022fastfold,
      title={FastFold: Reducing AlphaFold Training Time from 11 Days to 67 Hours}, 
      author={Shenggan Cheng and Ruidong Wu and Zhongming Yu and Binrui Li and Xiwen Zhang and Jian Peng and Yang You},
      year={2022},
      eprint={2203.00854},
      archivePrefix={arXiv},
}
@misc{DS4Sci_EvoformerAttention,
  author       = {DeepSpeed4Science},
  title        = {DS4Sci\_EvoformerAttention eliminates memory explosion problems for scaling Evoformer-centric structural biology models},
  howpublished = {\url{https://www.deepspeed.ai/tutorials/ds4sci_evoformerattention/}},
  year         = {2024},
  note         = {Accessed: 2024-11-06}
}

@article{steinegger2017mmseqs2,
  title={MMseqs2 enables sensitive protein sequence searching for the analysis of massive data sets},
  author={Steinegger, Martin and S{\"o}ding, Johannes},
  journal={Nature biotechnology},
  volume={35},
  number={11},
  pages={1026--1028},
  year={2017},
  publisher={Nature Publishing Group US New York}
}
@article{yuan2021oneflow,
  title={Oneflow: Redesign the distributed deep learning framework from scratch},
  author={Yuan, Jinhui and Li, Xinqi and Cheng, Cheng and Liu, Juncheng and Guo, Ran and Cai, Shenghang and Yao, Chi and Yang, Fei and Yi, Xiaodong and Wu, Chuan and others},
  journal={arXiv preprint arXiv:2110.15032},
  year={2021}
}
@article{uniprot2023uniprot,
  title={UniProt: the universal protein knowledgebase in 2023},
  journal={Nucleic acids research},
  volume={51},
  number={D1},
  pages={D523--D531},
  year={2023},
  publisher={Oxford University Press}
}
@article{chen2023rna,
  title={RNA tertiary structure modeling with BRiQ potential in CASP15},
  author={Chen, Ke and Zhou, Yaoqi and Wang, Sheng and Xiong, Peng},
  journal={Proteins: Structure, Function, and Bioinformatics},
  volume={91},
  number={12},
  pages={1771--1778},
  year={2023},
  publisher={Wiley Online Library}
}

@article {bernard24hasalphafold3,
        author = {Bernard, Cl{\'e}ment and Postic, Guillaume and Ghannay, Sahar and Tahi, Fariza},
        title = {Has AlphaFold 3 reached its success for RNAs?},
        elocation-id = {2024.06.13.598780},
        year = {2024},
        doi = {10.1101/2024.06.13.598780},
        publisher = {Cold Spring Harbor Laboratory},
        URL = {https://www.biorxiv.org/content/early/2024/07/22/2024.06.13.598780},
        % eprint = {https://www.biorxiv.org/content/early/2024/07/22/2024.06.13.598780.full.pdf},
        journal = {bioRxiv}
}

@dataset{baek23casp15,
  author       = {Baek, Minkyung and
                  McHugh, Ryan and
                  Anishchenko, Ivan and
                  Baker, David and
                  DiMaio, Frank},
  title        = {CASP15 RNA target predictions with RoseTTAFold-NA},
  month        = jan,
  year         = 2023,
  publisher    = {Zenodo},
  doi          = {10.5281/zenodo.7555957},
  url          = {https://doi.org/10.5281/zenodo.7555957}
}

@article{biasini2013openstructure,
  title={OpenStructure: an integrated software framework for computational structural biology},
  author={Biasini, Marco and Schmidt, Tobias and Bienert, Stefan and Mariani, Valerio and Studer, Gabriel and Haas, J{\"u}rgen and Johner, Niklaus and Schenk, Andreas Daniel and Philippsen, Ansgar and Schwede, Torsten},
  journal={Acta Crystallographica Section D: Biological Crystallography},
  volume={69},
  number={5},
  pages={701--709},
  year={2013},
  publisher={International Union of Crystallography}
}

@article{mariani2013lddt,
  title={lDDT: a local superposition-free score for comparing protein structures and models using distance difference tests},
  author={Mariani, Valerio and Biasini, Marco and Barbato, Alessandro and Schwede, Torsten},
  journal={Bioinformatics},
  volume={29},
  number={21},
  pages={2722--2728},
  year={2013},
  publisher={Oxford University Press}
}

@article{westbrook2015chemical,
  title={The chemical component dictionary: complete descriptions of constituent molecules in experimentally determined 3D macromolecules in the Protein Data Bank},
  author={Westbrook, John D and Shao, Chenghua and Feng, Zukang and Zhuravleva, Marina and Velankar, Sameer and Young, Jasmine},
  journal={Bioinformatics},
  volume={31},
  number={8},
  pages={1274--1278},
  year={2015},
  publisher={Oxford University Press}
}

@article{kunnakkattu2023pdbe,
  title={PDBe CCDUtils: an RDKit-based toolkit for handling and analysing small molecules in the Protein Data Bank},
  author={Kunnakkattu, Ibrahim Roshan and Choudhary, Preeti and Pravda, Lukas and Nadzirin, Nurul and Smart, Oliver S and Yuan, Qi and Anyango, Stephen and Nair, Sreenath and Varadi, Mihaly and Velankar, Sameer},
  journal={Journal of Cheminformatics},
  volume={15},
  number={1},
  pages={117},
  year={2023},
  publisher={Springer}
}

@article{mirabello2024dockq,
  title={DockQ v2: Improved automatic quality measure for protein multimers, nucleic acids, and small molecules},
  author={Mirabello, Claudio and Wallner, Bj{\"o}rn},
  journal={Bioinformatics},
  volume={40},
  number={10},
  pages={btae586},
  year={2024},
  publisher={Oxford University Press}
}
