## Supplementary material for "Protenix - Advancing Structure Prediction Through a Comprehensive AlphaFold3 Reproduction": figures and result summary: main.tex

\documentclass[
	a4paper, % Paper size, use either a4paper or letterpaper
	12pt, % Default font size, the template is designed to look good at 12pt so it's best not to change this
	%unnumberedsections, % Uncomment for no section numbering
]{CSSullivanBusinessReport}

\addbibresource{bibfile.bib} % BibLaTeX bibliography file
\usepackage{ragged2e}
\usepackage{hyperref}       % hyperlinks
\usepackage{url}            % simple URL typesetting
\usepackage{booktabs}       % professional-quality tables
\usepackage{amsfonts}       % blackboard math symbols
\usepackage{nicefrac}       % compact symbols for 1/2, etc.
\usepackage{microtype}      % microtypography
\usepackage{lipsum}
\usepackage{enumitem}
\usepackage{pgfplots}
% my inputs
\input{math_commands}
\usepackage{amsmath,bm,amsthm}
\newtheorem{theorem}{Theorem}[section]
\newtheorem{lemma}{Lemma}[section]
\newtheorem{corollary}{Corollary}[section]
\theoremstyle{definition}
\newtheorem{definition}{Definition}[section]

\usepackage{enumitem}
%\usepackage[vlined,ruled]{algorithm2e}
\usepackage{subfig}
\usepackage{xcolor,color}
\usepackage{graphicx}
\usepackage{wrapfig}
\usepackage{mathtools}
\usepackage{multirow,multicol}
\usepackage{booktabs}
\usepackage{lipsum}
\usepackage{sidecap}
\usepackage{cleveref}
\definecolor{deepmagenta}{rgb}{0.8, 0.0, 0.8}
\usepackage{letltxmacro}
\renewcommand{\eqref}{Eq.~\ref}
\usepackage{subcaption}
\usepackage{algorithm}
\usepackage{algpseudocode}
\usepackage{amssymb}% http://ctan.org/pkg/amssymb
\usepackage{pifont}% http://ctan.org/pkg/pifont
\newcommand{\cmark}{\ding{51}}%
\newcommand{\xmark}{\ding{55}}%
\definecolor{darkgreen}{rgb}{0.0, 0.3, 0.0} 
\definecolor{darkred}{rgb}{0.3, 0.0, 0.0}  % Dark red

\definecolor{mydarkblue}{rgb}{0,0.08,0.45}
\hypersetup{
 colorlinks=true,
 citecolor=mydarkblue,
 linkcolor=mydarkblue,
 urlcolor=blue}

% \captionsetup{
%     justification=centering,
%     format=hang
% }

\usepackage{tablefootnote}
\usepackage[export]{adjustbox}
\newcommand*{\Scale}[2][4]{\scalebox{#1}{$#2$}}%

\newcommand{\subfigcap}[2]{\fontfamily{phv}\selectfont {\textbf{#1} {\small #2}}}

\newcommand{\xs}[1]{{\color{red}{\bf\sf [xs: #1]}}}
\newcommand{\sw}[1]{{\color{brown}{\bf\sf [sw: #1]}}}
\newcommand{\hy}[1]{{\color{blue}{\bf\sf [hy: #1]}}}
\usepackage{xcolor}
\definecolor{myturquoise}{rgb}{0, 0.9, 0.6}
\definecolor{myblue}{rgb}{0, 0, 0.6}
\definecolor{mygray}{rgb}{0.6, 0.6, 0.6}
\definecolor{mysalmon}{RGB}{250, 128, 114}
\newcommand{\OurModelName}{{\it Protenix}}
\newcommand{\AFShort}{AF3}
\newcommand{\AFFull}{AlphaFold 3}
\newcommand{\chai}{Chai-1}
\newcommand{\AFmultierFull}{Alphafold-Multimer 2.3}
\newcommand{\AFmultierShort}{AF2.3}
\newcommand{\RFNAFull}{RoseTTAFold2NA}
\newcommand{\RFNAShort}{RF2NA}

\newcommand{\PbFull}{PoseBusters Benchmark Set Verison 2}
\newcommand{\PbShort}{PoseBusters V2}
\newcommand{\LddtFull}{Local Distance Difference Test}
\newcommand{\LddtShort}{LDDT}
\newcommand{\PlddtShort}{pLDDT}
\newcommand{\RmsdFull}{root-mean-square deviation}
\newcommand{\RmsdShort}{RMSD}
\newcommand{\LowhFull}{Low Homology}
\newcommand{\LowhShort}{LowH}

\usepackage{soul}
\usepackage{adjustbox}

\newcommand{\remove}[1]{{\color{gray}{\st{#1}}}}

\renewcommand*{\bibfont}{\small\rmfamily}  % Set font size and family for the bibliography

% Modify the URL appearance for online references
\AtEveryBibitem{
  \ifentrytype{online}{\def\UrlFont{\small\rmfamily}}{}  % Apply small and Times font for online URLs
}
\def\UrlFont{\small\rmfamily\color{mydarkblue}}

%----------------------------------------------------------------------------------------
%	REPORT INFORMATION
%----------------------------------------------------------------------------------------

\reporttitle{Protenix} % The report title to appear on the title page and page headers, do not create manual new lines here as this will carry over to page headers

\reportsubtitle{
%A Reproduction of AlphaFold3 \\
%and Open-Source Release
--- Advancing Structure Prediction Through a Comprehensive AlphaFold3 Reproduction}
%} % Report subtitle, include new lines if needed

\reportauthors{\it --- Advancing Structure Prediction Through a Comprehensive AlphaFold3 Reproduction}


\reportdate{\today} % Report date, include new lines for additional information if needed

\rightheadercontent{
\includegraphics[width=3cm]{figs/bytedance_logo_crop.png}
} % The content in the right header, you may want to add your own company logo or use your company/department name or leave this command empty for no right header content

%----------------------------------------------------------------------------------------

% \author{
%     Bytedance
%   %% \AND
%   %% Coauthor \\
%   %% Affiliation \\
%   %% Address \\
%   %% \texttt{email} \\
%   %% \And
%   %% Coauthor \\
%   %% Affiliation \\
%   %% Address \\
%   %% \texttt{email} \\
%   %% \And
%   %% Coauthor \\
%   %% Affiliation \\
%   %% Address \\
%   %% \texttt{email} \\
% }

\newcommand{\norm}[1]{\left\|#1\right\|}
\newcommand{\snorm}[1]{\|#1\|}

\newcommand{\figfont}[1]{{\fontfamily{phv}\selectfont #1}}


\newcommand{\keypoint}[1]{\textcolor{red}{\textbf{[#1 ]}}}
\newcommand{\outline}[1]{\textcolor{blue}{[#1 ]}}

\newcommand{\comment}[1]{\textcolor{purple}{[#1 ]}}

\newcommand{\cy}[1]{\textcolor{red}{[cy: #1 ]}}

\newcommand{\eric}[1]{\textcolor{green}{[eric: #1 ]}}

\definecolor{mycolor}{RGB}{219,90,107}
\newcommand{\kq}[1]{{\textcolor{mycolor}{#1}}}
\renewcommand{\thefootnote}{\fnsymbol{footnote}}

\begin{document}


%----------------------------------------------------------------------------------------
%	TITLE PAGE
%----------------------------------------------------------------------------------------

\thispagestyle{empty} % Suppress headers and footers on this page

\vspace*{-0.075\textheight} % Pull logo into the top margin
\vspace{1cm}
\hspace{0.4\textwidth}\includegraphics[width=0.6\textwidth]{figs/bytedance_logo.png} % Company logo
\vspace{0.1\textheight} % Vertical whitespace
 % \vspace{1cm}

\parbox{0.9\textwidth}{
        \fontsize{45pt}{50pt} %\fontfamily{fvm}\selectfont
        \raggedright
        \selectfont
        % \textbf{\reporttitle}
        \fontfamily{qag}\selectfont \textbf{\reporttitle}
        \par
    } % Report title, intentionally at less than full width for nice wrapping. Adjust the width of the \parbox and the font size as needed for your title to look good.
\vspace{0.08\textheight} % Vertical whitespace

{\Large \raggedleft
 \fontfamily{qag}\selectfont \textbf{\reportsubtitle}\par
 } % Subtitle
% {\Large \itshape \raggedleft --- Advancing Structure Prediction Through a Comprehensive AlphaFold3 Reproduction\par} % Report authors, group or 

\vspace{0.2\textheight} % Vertical whitespace
\subsection*{ByteDance AML AI4Science Team}

Xinshi Chen\footnotemark[1], 
Yuxuan Zhang\footnotemark[1], Chan Lu,  Wenzhi Ma, Jiaqi Guan, Chengyue Gong, Jincai Yang, Hanyu Zhang, Ke~Zhang, Shenghao Wu, Kuangqi Zhou, Yanping Yang, Zhenyu Liu, Lan Wang, Bo~Shi,  Shaochen Shi, Wenzhi Xiao%\footnotemark[2]
\footnotemark[4]
\footnotetext[1]{Contributed equally.}
\footnotetext[4]{Contact: {}}

	
% Push the following down to the bottom of the page} % Report authors/group/department, include new lines if needed
\vfill % Push the following down to the bottom of the page
\vfill\vfill\vfill % Vertical whitespace

% {\large\reportdate\par} % Report date
{\large November 9, 2024}
% \end{fullwidth}

\newpage

%----------------------------------------------------------------------------------------
%	DISCLAIMER/COPYRIGHT PAGE
%----------------------------------------------------------------------------------------

% \thispagestyle{empty} % Suppress headers and footers on this page

% \begin{twothirdswidth} % Content in this environment to be at two-thirds of the whole page width
% 	\footnotesize % Reduce font size
	
% 	\subsection*{Disclaimer}

% 	Lorem ipsum dolor sit amet, consectetur adipiscing elit. Praesent porttitor arcu luctus, imperdiet urna iaculis, mattis eros. Pellentesque iaculis odio vel nisl ullamcorper, nec faucibus ipsum molestie. Sed dictum nisl non aliquet porttitor. Etiam vulputate arcu dignissim, finibus sem et, viverra nisl. Aenean luctus congue massa, ut laoreet metus ornare in. Nunc fermentum nisi imperdiet lectus tincidunt vestibulum at ac elit.
	
% 	\subsection*{Copyright}
	
% 	\textcopyright~[2024] [ByteDance] 
	
% 	Copyright notice text\ldots In hac habitasse platea dictumst. Curabitur mattis elit sit amet justo luctus vestibulum. In hac habitasse platea dictumst. Pellentesque lobortis justo enim, a condimentum massa tempor eu. Ut quis nulla a quam pretium eleifend nec eu nisl. Nam cursus porttitor eros, sed luctus ligula convallis quis.
	
% 	\subsection*{Authors}
	
% 	Contact:
	
% 	\vfill % Push the following down to the bottom of the page
	
% 	\subsubsection*{Changelog}
	
% 	\scriptsize % Reduce font size further
	
% 	\begin{tabular}{@{} L{0.05\linewidth} L{0.15\linewidth} L{0.6\linewidth} @{}} % Column widths specified here, change as needed for your content
% 		\toprule
% 		v1.0 & 2024-11-09 & Initial report.\\
		
% 		\bottomrule
% 	\end{tabular}
% \end{twothirdswidth}

% \newpage

%----------------------------------------------------------------------------------------
%	TABLE OF CONTENTS
%----------------------------------------------------------------------------------------


\tableofcontents % Output the table of contents, automatically generated from the section commands used in the document


\newpage

%----------------------------------------------------------------------------------------
%	SECTIONS
%----------------------------------------------------------------------------------------


\section{Introduction}
\label{sec:intro}

\input{tex/intro}

\section{Results}
\label{sec:result}

\input{tex/results}

\section{Methods}
\label{sec:method}

\input{tex/method}

\section{Accessibility}

\input{tex/accessibility}

\section{Future Plan}

\input{tex/plan}


\newpage


%----------------------------------------------------------------------------------------
%	 REFERENCES/BIBLIOGRAPHY
%----------------------------------------------------------------------------------------

\newpage

\addcontentsline{toc}{section}{Reference List} % Add the bibliography to the table of contents


\printbibliography[title=Reference List] % Output the bibliography with a custom section title

%----------------------------------------------------------------------------------------
%	APPENDICES
%----------------------------------------------------------------------------------------

\newpage


\appendix


\section{Dataset}

% \subsection{Data Pipeline}
% \label{app:data-pipeline}
% Inclusion of various biomolecules creates exceptional challenges for data curation and featurization. We dive into the details of \AFShort{} supplementary materials to reproduce the data pipeline. Below we list the major differences in our implementation.

% \textbf{Parser.} 
% In selecting alternative locations, we use the first occupancy rather than the largest one. This is because we found that using the largest occupancy may result in some adjacent residues adopting different conformations, preventing the formation of covalent bonds and leading to chain breaks.

% \textbf{MSA.}
% We search MSAs by MMSEQS2~\cite{steinegger2017mmseqs2}, and use MSAs from the Uniref100~\cite{uniprot2023uniprot} database for pairing (species are identified with taxonomy IDs).

% \textbf{Templates.} 
% We do not use templates.

% \textbf{Cropping.} 
% In our contiguous cropping method, we discard metals and ions. According to the method described by \AFShort{}, metals and ions are easily selected in contiguous cropping. Predicting the coordinates of these atoms becomes challenging and may result in significant loss during training when the cropped metals and ions are not surrounded by other biomolecules. In all cropping methods, since ligands and non-standard amino acids are represented by a single atom per token, they may be cropped into fragments. Therefore we ensure that ligands and non-standard amino acids are not fragmented into incomplete segments in cropping. 

% \textbf{Featurization.} 
% \begin{itemize}
%     \item \textbf{Reference features.} We use the version of CCD~\cite{westbrook2015chemical} downloaded on 2024-06-08 and generate RDKit conformers using pdbeccdutils~\cite{kunnakkattu2023pdbe} v0.8.5. When RDKit conformer generation fails, we use the CCD ideal coordinates and do not use the representative coordinates to avoid potential data leakage
%     \item \textbf{Bond features.} Bond features between tokens only include bonds within ligands, bonds between ligands and polymers, and bonds within non-standard residues.
% \end{itemize}


\subsection{Training Dataset}
\label{app:training-dataset}

In addition to WeightedPDB, we also use AlphaFold 2 predictions to train our model. We use representative sequences from MGnify \cite{richardson2023mgnify} with greater than 200 residues as the distillation data source. Due to resource limitations, we run 5 models on a small part of the data and only 1 model (model-3) on the rest. As a supplement, we also use a subset of the OpenProteinSet~\cite{ahdritz2023openproteinset}. All distillation datasets are filtered based on pLDDT to obtain high-quality training data.


\begin{table}[h]
    \centering
    \caption{ \justifying \textbf{Datasets used to train \OurModelName{}.} \small We list the descriptions for the training datsets.}
    \label{tab:training-dataset}
    \begin{tabular}{l|m{5cm}|c|c}
    \toprule
     Name & Description & Sampl. strategy & Weight \\
    \midrule 
     Weighted PDB &  Ground truth PDB structures & weighted & 0.60 \\
     \midrule
     Protein monomer distillation (Set1) & Protein monomer predictions from MGnify using 1 model & uniform & 0.35 \\
     \midrule
     Protein monomer distillation (Set2) & Protein monomer predictions from MGnify using 5 models & uniform &  0.03 \\
     \midrule
     OpenProteinSet~\cite{ahdritz2023openproteinset} subset & Protein monomer predictions from Uniclust30 & uniform & 0.02 \\
    \bottomrule
    \end{tabular}
\end{table}

\subsection{Evaluation Dataset}
\label{app:evaluation-dataset}


% \begin{table}[h]
%     \centering
%     \caption{ \textbf{Datasets used to benchmark the model.} Two commonly-used benchmark dataset have been used. We list the details for these two datasets. `\#' denotes the number of the objects.
%     }
%     \label{tab:evaluation-set}
%     \begin{tabular}{l|cccc}
%     \toprule
%      Name & Structure & \#PDB entries & \#Tokens & \#Atoms \\
%     \midrule 
%     \PbShort{}~\cite{buttenschoen2024posebusters}  &  Asymmetric unit & 308 & No Limit & No Limit\\
%     LowH Recent PDB (\ref{app:recentpdb-dataset}) & First bioassembly &  & $\leqslant$2560 & No Limit \\
%     \bottomrule
%     \end{tabular}
% \end{table}


\subsubsection{PoseBusters V2}


We use the \PbFull{} (\PbShort{})~\cite{buttenschoen2024posebusters} for evaluating protein-ligand interactions. This dataset contains 308 structures.
To facilitate a fair comparison with \AFShort{}, we follow their procedure~\cite{abramson2024af3} to remove chains clashing with the ligand of interest (specifically, files: 7O1T, 7PUV, 7SCW, 7WJB, 7ZXV, 8AIE). Similarly, for PDB entry 8F4J, we retain only the protein chains within 20 \AA{} of the ligand.
We follow the \AFShort{} practice, in which the model inference is performed on the \textit{asymmetric unit} of each structure, to allow for direct comparability with both the numbers reported in \AFShort{} and their released predictions.

\subsubsection{Low Homology Recent PDB Set}
\label{app:recentpdb-dataset}
We follow the Supplementary Information of \AFShort{} to curate the Low Homology (LowH) Recent PDB set for evaluating the prediction of protein and nucleic acid interfaces. This set consists of PDB targets released between 2022-05-01 and 2023-01-12, filtered for sequence similarity below 40\% to better assess model generalization. We apply the same token cutoff to evaluate models on structures with fewer than 2560 tokens. However, due to some missing details in the paper and various possible differences related to clustering, low homology filtering, and other factors, our dataset still differs in PDB and cluster numbers compared to the \AFShort{} paper, despite our efforts with different tools and settings. The number of clusters used for each evaluation task is specified in the figures.

\subsubsection{Nucleic Acids Evaluation Set}
\label{app:nuc-eval-set}


\textbf{CASP15.} Eight targets that were publicly available as of December 1, 2023 are used for the evaluation: R1116/8S95, R1117/8FZA, R1126 (https://predictioncenter.org/casp15/TARGETS\_PDB/R1126.pdb), R1128/8BTZ, R1136/7ZJ4, R1138/[7PTK/7PTL], R1189/7YR7 and R1190/7YR6.

\textbf{Protein-Nucleic Acids Complexes.} The dataset used for the results in Figure~\ref{fig:summary} is curated by the following procedure. The nucleic acid evaluation set is screened from the RecentPDB dataset's PDB IDs provided in Table 14 of the \AFShort{} supplementary materials. RoseTTAFold2NA can only input up to 1000 residues, hence the dataset does not include systems with a total residue count exceeding 1000. Additionally, it does not contain systems with both DNA and RNA present simultaneously. Subsequently, we retain structures with only one RNA chain and one protein chain, or with two DNA chains and one protein chain. We then remove all systems containing non-standard residues in the DNA. The remaining structures coincided with the number in \AFShort{} Figure 1, 25 RNA-Protein structures and 38 dsDNA-Protein structures. Although the number here matches that in \AFShort{}, it does not necessarily mean the evaluation was conducted on an identical evaluation set, which may lead to discrepancies in the metrics. % In order to promote reproducibility, the detailed list of PDB IDs comprising this set can be accessed in the \href{https://github.com/bytedance/Protenix}{Protenix repository}.


\section{Evaluation Workflow }

\label{app:evaluation}

For each PDB entry, we perform the following steps:
\begin{enumerate}[left=0cm]
    \item \textbf{Inference and Sampling.} 
    
    Unless specified, we generate 25 samples using 5 model seeds. Each sample is exported as both a CIF file and a JSON file, with the JSON file containing confidence ranking scores. For targets in PoseBusters V2, we predict the asymmetric unit of the structure; for targets in all other datasets, we predict the first bioassembly of the structure. No cropping is applied during inference, and we generally use 10 recycles unless otherwise noted. \AFShort{} specifies its configuration for recycles only in Appendix 5.10, where inference times are reported using 10 trunk recycles on 16 GPUs. Therefore, we infer that \AFShort{} likely used at least 10 recycles in their report. 

    \item \textbf{Chain and Atom Permutation.} For each predicted structure (CIF), we perform chain mapping to establish a one-to-one correspondence between predicted and true entities. Once equivalent chains are identified, we permute the chains and atoms to better align the prediction with the true structure (details in Appendix~\ref{app:permutation}).
    \item \textbf{Metrics Computation.} Different types of interactions are evaluated with specific methods. For proteins, we use the DockQ Python package to calculate DockQ scores. For predictions on PoseBusters V2, we follow these steps:
    \begin{enumerate}[left=0cm]
        \item \textit{Exporting Structures}: 
    The ligand of interest is exported to an SDF file, while all other chains with at least one atom within 5 \AA{} of any atom in the predicted ligand are exported to a PDB file.
    \item \textit{PoseBusters Analysis}: Finally, we use the PoseBusters Python package v.0.2.7 to calculate the \RmsdShort{} of the ligands and evaluate any structural violations using the exported files.
    \end{enumerate}
    
    % We permute equivalent chains and atoms within each sample to optimize the accuracy of the predicted ligand and main chain. In some instances, we observe that the ligand and main chain align closely with the reference structure, while the remaining chains show significant deviations.
    % Similar discrepancies are observed in predictions released by \ (see Appendix \ref{} for details) \cy{do we still plan to have these case studies? }.
    % \item \textbf{Structural Alignment}: We align the predicted structure to the reference by minimizing the \RmsdShort{} of the backbone atoms in the pocket region.

\end{enumerate}


\section{Chain and Atom Permutation}
\label{app:permutation}


\begin{algorithm}[H]
    \centering
    \caption{Chain Permutation (Training)}\label{app:perm_train}
    \begin{algorithmic}[1]
        \State \textbf{Input:} $\{x_k^\text{pred}\}$,$\{x_l^\text{gt}\}$ \hfill $x_k^\text{pred}\in \mathbb{R}^{n_k\times3}, x_l^\text{gt} \in \mathbb{R}^{n_l\times3}$
        \State $a^\text{pred} \gets FindPredAnchorChain(\{x_k^\text{pred}\})$   \hfill $a^\text{pred} \in \mathbb{N}$ \label{alg:find_pred_anchor_train}
        \State $\{a^\text{gt}_m\} \gets FindGTAnchorChains(\{x_l^\text{gt}\})$ \hfill $a^\text{gt}_m \in \mathbb{N}$ \label{alg:find_gt_anchor_train}
        \State $T_{m} \gets GetTransform(x_{a^\text{gt}_m}^\text{gt},x_{a^\text{pred}}^\text{pred})$ \label{alg:transform_gt}
        \State $x_l^\text{gt} \gets T_m\cdot x_l^\text{gt}$
        \State $p_m$ $\gets GreedyMatchRemainingChains(\{x_k^\text{pred}\},\{x_l^\text{gt} \})$ \hfill $p_m \in \mathbb{N}^{N_\text{chains}}$ \label{alg:greedy_match}
        \State $e_m \gets sum_k(RMSD(x_k^\text{pred},x_{{p_{mk}}}^\text{gt}))$
        \State \textbf{Return}  $p_\text{argmin($e_m$)}$
    \end{algorithmic}
\end{algorithm}

\begin{algorithm}[H]
    \centering
    \caption{Chain Permutation (Inference)}\label{app:perm_test}
     \begin{algorithmic}[1]
        \State \textbf{Input:} $\{x_k^\text{pred}\}$,$\{x_l^\text{gt}\}$ \hfill $x_k^\text{pred}\in \mathbb{R}^{n_k\times3}, x_l^\text{gt} \in \mathbb{R}^{n_l\times3}$
        \State $\{a^\text{gt}_i\},\{a^\text{pred}_j\} \gets FindAnchorPairs(\{x_k^\text{pred}\},\{x_l^\text{gt}\})$ \hfill $a^\text{gt}_i,a^\text{pred}_j \in \mathbb{N}$ \label{alg:find_anchor_pairs}
        \State $T_{ij} \gets GetTransform(x_{a^\text{gt}_i}^\text{gt},x_{a^\text{pred}_j}^\text{pred})$
        \State $x_l^\text{gt} \gets T_{ij}\cdot x_l^\text{gt}$
        \State $p_{ij}$ $\gets GreedyMatchRemainingChains(\{x_k^\text{pred}\},\{x_l^\text{gt} \})$ \hfill $p_{ij} \in \mathbb{N}^{N_\text{chains}}$
        \State $e_{ij} \gets sum_{k}(RMSD(x_k^\text{pred},x_{{p_{ijk}}}^\text{gt}))$ 
        \State \textbf{Return}  $p_\text{argmin($e_{ij}$)}$
    \end{algorithmic}
\end{algorithm}

To address ambiguities in chain and atom naming for accurate loss and metric calculations, we introduce chain and atom permutations at various stages. The methods are detailed below.

\textbf{Training.} During training, we permute the chains and atoms of the ground truth structure to align with the mini-rollout predictions before the diffusion sampling stage. Our chain permutation approach largely builds on the heuristic algorithms used in \AFShort{} ~\cite{abramson2024af3} and \AFmultierShort{} ~\cite{evans2021protein}. Initially, anchor chains in the ground truth and predicted structures are identified for alignment. This is followed by a greedy matching process for the remaining chains (Algorithm~\ref{app:perm_train}, Line~\ref{alg:greedy_match}). The permutation that minimizes the average RMSD between the prediction and ground truth chains is selected. For better efficiency and applicability, we also introduce several adaptations: we only consider entities with at least four resolved tokens for anchor chains and prioritize polymer chains to reduce permutation complexity. Additionally, to account for cropping during training,  we first identify an anchor chain in the predicted structure, then enumerate candidate anchor chains in the ground truth (Algorithm~\ref{app:perm_train}, Line~\ref{alg:find_pred_anchor_train} and \ref{alg:find_gt_anchor_train}). Specifically, for samples using spatial cropping, we restrict the ground truth anchor chain to chains included in the cropped region, as this cropping method may exclude certain chains during training. In contrast, for contiguous cropping, we lift this restriction, as all chains, even if cropped, remain included throughout the process.

We then perform atom permutation on the ground-truth structure within each residue/ligand to correct atom-level computation. This process begin by first aligning the global structure of the prediction to that of the ground truth, followed by residue-level permutation to minimize within-residue RMSD.

    
    
\textbf{Inference.}
During inference, we permute the chains and atoms in the structure predicted by the diffusion module, aligning them to the ground truth independently for each diffusion sample. The chain permutation approach here differs slightly from the training stage. First, we identify anchor chains in both the predicted and ground truth structures, requiring a minimum of four resolved tokens (Algorithm~\ref{app:perm_test}, Line~\ref{alg:find_anchor_pairs}). We then enumerate all anchor chain pairs between the prediction and ground truth and greedily matched the remaining chains for each pair, selecting the permutation which minimizes the global RMSD. 
   
When applying chain permutation to ligand complexes (e.g., \PbShort{}), 
we generate all possible combinations of pockets and ligands, and then select the permutation resulted in the lowest ligand-specific RMSD. 
For atom permutation, we align the predicted structure to the ground truth based specifically on the coordinates of pockets and ligands, rather than using global coordinates.
%ToDo: compare different permutation algorithms.
\end{document}
