## Supplementary material for "Protenix - Advancing Structure Prediction Through a Comprehensive AlphaFold3 Reproduction": figures and result summary: accessibility.tex

\label{sec:access}


Source code and datasets are available at the \href{https://github.com/bytedance/Protenix}{Protenix repository}\footnote{\url{https://github.com/bytedance/Protenix}}. We encourage readers to refer to the source code for detailed use cases.


\textbf{Data release.}
Our data release aims to provide a comprehensive foundation for researchers to reproduce results, conduct further analysis, or leverage the datasets for new applications. We highlight our contributions as follows:
\begin{itemize}[left=0cm]
    \item \textbf{Significant computational efforts}: 
    The processed data includes time-consuming MSA (multiple sequence alignment) search results, covering all instances in the wwPDB. This is especially beneficial for researchers with limited access to computing resources, saving them considerable time and effort in data processing.
    \item \textbf{PDB ID to processed data mapping}: To facilitate ease of use, we curate the dataset and provide a mapping from Protein Data Bank (PDB) IDs to the processed data format. This allows users to quickly integrate the data into their pipelines without extensive adjustments, streamlining their research process.
    % \item \textbf{Release of Inference Results on Benchmark (\PbShort{})}: To enable direct comparison, we release inference results on the \PbShort{} benchmark. This allows users to evaluate the model’s performance and compare it with their own results, providing a standardized reference for benchmarking.
\end{itemize}

\textbf{Code release.}
Our code release is designed to maximize usability and encourage experimentation, offering researchers a robust set of tools for extending our work:
\begin{itemize}[left=0cm]
    \item \textbf{Inference code}: We provide code for model inference, along with a notebook that allows users to customize and model their inputs.
    \item \textbf{Trainable code}: We release trainable code, allowing users to retrain or fine-tune models on additional datasets. This is beneficial for those interested in adapting the model to other applications or improving model performance on specific tasks.
    \item \textbf{Documentation}: Comprehensive documentation is included in our codebase to guide users through setting up, running, and modifying the code, making it easy for users at all skill levels to get started.
\end{itemize}


\iffalse
\comment{
Follow github README.md\\
Who will write README?
}


\outline{data}

  - Outlined steps for data acquisition and preprocessing. 
  
    - List of clarifications and adjustments(Appendix)
  
  - Emphasized significant computational efforts involved.
  
  - Data curation for release, dataset input configs
  
    - PDB ID to proccessed data
  
  - Release inference results on benchmark(posebuster)
  
\outline{code}

- Trainable code

- Documentation [ReadME.md]
\fi
