## Supplementary material for "Protenix - Advancing Structure Prediction Through a Comprehensive AlphaFold3 Reproduction": figures and result summary: intro.tex

\justifying   

Understanding the three-dimensional structures of biological molecules has revolutionized our ability to study and manipulate  molecular functions. Over recent decades, significant progress in protein structure prediction has been made through various contributions \cite{senior2020alphafold, jumper2021af2, evans2021afmultier, abramson2024af3, baek2021rosettafold,du2021trrosetta, mirdita2022colabfold, wu2022high, lin2023esmfold, ahdritz2024openfold}. However, predicting complex structures that involve multiple biomolecules—such as protein-ligand, protein-nucleic acid, and antibody-antigen interactions—poses additional challenges.  Recent works like NeuralPLexer \cite{qiao2024state}, Umol \cite{bryant2024structure}, and RoseTTAFold-AA \cite{krishna2024rosettafoldAA} have targeted these complex prediction tasks. Among these, \AFFull{} (\AFShort{})~\cite{abramson2024af3} has set a new milestone, representing a significant leap forward in this domain.


Despite the impressive advances of \AFShort{}, limited accessibility has constrained its broader adoption within the research community. The absence of code and certain ambiguities and typographical errors in the \AFShort{} paper present additional challenges for machine learning and computational biology researchers seeking to reproduce or improve the model. Open-source initiatives like 
HelixFold3~\cite{helixfold3} and Chai-1~\cite{chai1} have made significant strides in democratizing access to these advanced models.  However, the lack of comprehensive training code and preprocessed data remains a barrier for researchers aiming to fully reconstruct and utilize these models. To address these challenges, we introduce \OurModelName{}, designed to lower these barriers and better support the interdisciplinary research community.


Below, we outline our key contributions:

\begin{itemize}[left=0em]

\item \textbf{Model performance.} We benchmark \OurModelName{} against \AFShort{}~\cite{abramson2024af3}, \AFmultierFull{} (\AFmultierShort{})~\cite{evans2021afmultier}, and \RFNAFull{} (\RFNAShort{})~\cite{baek2024accurate}. \OurModelName{} achieves strong performance in predicting structures across different molecular types (Figure~\ref{fig:summary}). As a fully open-source model, it empowers researchers to generate novel predictions and fine-tune the model for specialized applications.


\item \textbf{Methodology.}  We implement \OurModelName{} based on the descriptions provided in \AFShort{} as part of our reproduction effort. We refine several ambiguous steps, correct typographical errors, and make targeted adjustments based on our observations of the model's behavior. By sharing our reproduction experience, we aim to support the community’s efforts to build on these improvements and drive further advancements in the field.


\item \textbf{Accessibility.} We have open-sourced \OurModelName{}, providing model weights, inference code, and trainable code for research purposes, as detailed in Section~\ref{sec:access}.

\end{itemize}
