## Supplementary material for "Protenix - Advancing Structure Prediction Through a Comprehensive AlphaFold3 Reproduction": figures and result summary: method.tex

\subsection{Data Pipeline}
Inclusion of various biomolecules creates exceptional challenges for data curation and featurization. We dive into the details of \AFShort{} supplementary materials to reproduce the data pipeline. Below we list the major differences in our implementation.

\textbf{Parser.} 
In selecting alternative locations, we use the first occupancy rather than the largest one. This is because we found that using the largest occupancy may result in some adjacent residues adopting different conformations, preventing the formation of covalent bonds and leading to chain breaks.

\textbf{MSA.}
We search MSAs using MMSEQS2~\cite{steinegger2017mmseqs2} and ColabFold~\cite{mirdita2022colabfold} MSA pipeline, and use MSAs from the Uniref100~\cite{uniprot2023uniprot} database for pairing (species are identified with taxonomy IDs). We do not use MSA for nucleic chains.

\textbf{Templates.} 
We do not use templates.


\textbf{Featurization.} 
\begin{itemize}[left=0cm]
    \item \textbf{Reference features.} We use the version of CCD~\cite{westbrook2015chemical} downloaded on 2024-06-08 and generate RDKit conformers using pdbeccdutils~\cite{kunnakkattu2023pdbe} v0.8.5. When RDKit conformer generation fails, we use the CCD ideal coordinates and do not use the representative coordinates to avoid potential data leakage.
    \item \textbf{Bond features.} Bond features between tokens only include bonds within ligands, bonds between ligands and polymers, and bonds within non-standard residues.
\end{itemize}


\textbf{Cropping.} Our implementation of the cropping method takes the following aspects into consideration.
\begin{itemize}[left=0cm]
    \item In our contiguous cropping method, we discard metals and ions, as this approach cannot guarantee that cropped atoms remain close to each other. When metals or ions are cropped in isolation from other chains, it becomes challenging for the model to accurately predict their positions, potentially leading to large training losses. Therefore, we exclude metals and ions from contiguous cropping but retain them in spatial cropping.
    \item Since ligands and non-standard amino acids are represented by a single atom per token, they may be cropped into fragments. To prevent this, we ensure that ligands and non-standard amino acids remain intact and are not split into incomplete segments during cropping.
    \item We account for the impact of cropping on chain permutation during training. For spatial cropping, we restrict permutation to chains within the cropped region to ensure predictions focus on spatially close structures. In contrast, for contiguous cropping, where cropped chains may be distant but equivalent chains may be closer to each other, we allow equivalent permutations and adjust the cropped region based on model predictions. Details on permutation handling are provided in Appendix~\ref{app:permutation}.
\end{itemize}


\begin{table}[hbtp]
\centering
\caption{ \textbf{Corrections.} We provide a description and explanation of the introduced corrections in our implementation. The algorithm numbers correspond to those in the \AFShort{} Supplementary materials.
}
\label{tab:corrections}
\begin{tabular}{p{2.3cm}|p{4.8cm}|p{8.3cm}}
\toprule
\textbf{Algorithm} & \textbf{Correction} & \textbf{Explanation} \\ \midrule
Algorithm 1 
\newline Line 10 & 
\textcolor{darkgreen}{\st{$\{ \mathbf{z}_{ij} \} \mathrel{+}= \text{MsaModule}$}}
\newline 
$\{ \mathbf{z}_{ij} \} = \text{MsaModule}$
& 
The MsaModule itself contains residual connections for updating $\mathbf{z}$, hence adding an additional residual update would be redundant. \\ \midrule

Algorithm 3 \newline Line 8 & 
\textcolor{darkgreen}{\st{If not $b_{ij}^{\text{same\_chain}}$}}
\newline If $b_{ij}^{\text{same\_entity}}$ & 
Comparing the symmetry IDs ($\text{sym\_id}$) only makes sense for chains that share the same entity ID. If two chains belong to different entities, their symmetry IDs are irrelevant, and the original comparison would be logically flawed. \\ \midrule

Algorithm 18 \newline Line 9 
& 
\textcolor{darkgreen}{\st{$\vec{\delta}_l = ( \vec{x}_l - \vec{x}_l^{\text{denoised}} ) / \hat{t}$}}
\newline 
$\vec{\delta}_l=( \vec{x}_l^{\text{noisy}} - \vec{x}_l^{\text{denoised}}) / \hat{t}$ 
& 
The original form in Line 9 does not align with Line 11, 
$
        \vec{x}_l \gets \vec{x}_l^{\text{noisy}} + \eta \cdot dt \cdot \vec{\delta}_l,
$
which follows an Euler method for solving ODEs. The correction ensures mathematical consistency.
\\
\midrule

Algorithm 23 \newline Line 2
&
\textcolor{darkgreen}{\st{$
\{\mathbf{b}_i\}=\text{AttentionPairBias}
$}}
\newline
$
\{\mathbf{b}_i\}{\mathrel{+}=}\text{AttentionPairBias}
$
&
Empirically we find the model does not converge well if residual connection is missing in the DiffusionTransformer block.
\\
\midrule
Algorithm 28
\newline
Line 11
&
\textcolor{darkgreen}{\st{$\vec{\mathbf{x}}_l^{\text{align}}=R \vec{\mathbf{x}}_l + \vec{\mu} $}}
\newline
$\vec{\mathbf{x}}_l^{\text{align}}=R \vec{\mathbf{x}}_l + \vec{\mu}^{\text{GT}} $
&
We believe this is a typo. Otherwise, this is mathematically inconsistent.
\\
\midrule
Algorithm 31
\newline
Line 3
&
\textcolor{darkgreen}{\st{$[ 3 \frac{3}{8} \, \text{\AA{}}, \, 5 \frac{5}{8} \, \text{\AA{}}, \, \dots, \, 21 \frac{3}{8} \,\text{\AA{}} ]$}}
\newline
$[ 3 \frac{3}{8} \, \text{\AA{}}, \, 4 \frac{5}{8} \, \text{\AA{}}, \, \dots, \, 21 \frac{3}{8} \,\text{\AA{}} ]$
& 
We believe this is a typo. After fixing it, the distances can be discretized into 15 bins of equal width 1.25 \AA{}.
\\
\midrule
Supp. 3.7.1
\newline
Equation (6) & 
\textcolor{darkgreen}{\st{$(\hat{t}^2+\sigma_{\text{data}}^2)/(\hat{t}+\sigma_{\text{data}})^2$}}
\newline
$(\hat{t}^2+\sigma_{\text{data}}^2)/(\hat{t}\times\sigma_{\text{data}})^2$
& We believe this is a typo because the per-sample weighting of the diffusion loss, which is different from the original form in \AFShort{}, is carefully designed in a principled way by the authors of EDM~\cite{karras2022edm}. 
\\
 \bottomrule
\end{tabular}
\end{table}

\subsection{Model and Training}

We refer readers to the supplementary materials of \AFShort{} for detailed methodology, as our implementation is based on it. In this section, we primarily highlight the differences we have introduced in our implementation and explain the underlying reasons for making these modifications.


\textbf{Corrections and adjustments.}  In our analysis of the algorithms presented in the original paper, we identify several errors and ambiguities. Some of the adjustments are key to performance. These corrections and their corresponding explanations are shown in Table~\ref{tab:corrections}.  Additionally, we make slight modifications to the confidence head by incorporating LayerNorm and adding a few linear layers (see our released code). We find that the confidence loss does not converge as effectively when implemented exactly as described in the original paper.


\textbf{Parameter initialization.} We do not conduct an extensive analysis of the impact of different initialization strategies. However, we apply zero-initialization to several modules. Specifically, some linear layers are zero-initialized to ensure that each residual layer behaves as an identity operation at initialization. Following recent work on conditional diffusion models~\cite{peebles2023dit, esser2024sd3}, we zero-initialize two linear layers in the AdaptiveLayerNorm. Additionally, we apply zero initialization to the N-cycle loop connection blocks in the Pairformer (see Algorithm 1, line 8 in \AFShort{} report). Empirically, we find that these zero-initializations help prevent dimensional collapse in the network weights and mitigate the explosion of hidden values.


\textbf{Scalability.} 
Despite having only about 386 million parameters, 
\AFShort{} requires an enormous amount of computation.
Especially, its Pairformer blocks lead to numerous intermediate activations. 
Additionally, during training, the diffusion module has a large diffusion batch size and applies activations for each atom pair, thereby creating memory bottlenecks for both training and inference. 
To tackle these challenges and enhance training efficiency, we have introduced several approaches as follows:
\begin{itemize}[left=0cm]    

    \item  \textbf{BF16 mixed precision training.} Compared to FP32 training, BF16 training significantly reduces peak memory usage and nearly doubles the actual training speed. However, not all GPU architectures support BF16, with compatibility limited to specific models like NVIDIA's Ampere~\cite{krashinsky2020nvidiaampere} and Hopper~\cite{harris2022nvidiahopper}.

    \item \textbf{Custom CUDA Kernels.} A naive implementation of LayerNorm in PyTorch \cite{paszke2019pytorch} often encounters memory limitations. Building on optimizations from FastFold \cite{cheng2022fastfold} and OneFlow \cite{yuan2021oneflow}, we develop a custom LayerNorm implementation that adaptively adjusts block and grid sizes while leveraging shared memory to optimize GPU utilization. This approach yields a 30\%-50\% improvement in end-to-end training speed compared to PyTorch's native LayerNorm across various training stages.
    
    \item \textbf{DS4Sci\_EvoformerAttention.} By utilizing the DS4Sci\_EvoformerAttention kernels from DeepSpeed4Science \cite{DS4Sci_EvoformerAttention}, a 10\%-20\% improvement in end-to-end training speed across various training stages is achieved.

    
    \item \textbf{Other Considerations.} A direct implementation of the model update steps, as outlined in the \AFShort{} paper, leads to the creation of unnecessarily large tensors. To avoid this, we use alternative implementations that prevent the formation of these large tensors. Additionally, to address memory bottlenecks during both training and inference, we employ several optimization techniques, including gradient checkpointing, in-place operations, tensor offloading, and chunking, similar to the approach used in OpenFold~\cite{ahdritz2024openfold}.

    
\end{itemize}


\textbf{Training.}  \OurModelName{} is trained on 192 GPUs  over approximately two weeks. Table~\ref{tab:training_stage_config} provides a summary of our training stages. We also simplified the hyperparameters considerably, as our training utilizes the Protein Monomer Distillation dataset only, excluding other distillation sets.
Although we implement a multi-stage training setup similar to \AFShort{}, our model is trained with significantly fewer steps.  We train the model for 75K steps during the initial stage and 15K steps during fine-tuning stage 1. We run only 4K steps for fine-tuning stage 2, which may be insufficient for adequately training the confidence head.


\begin{table}[t]
\centering
\caption{\textbf{Training Stages Configurations.} We list the key hyper-parameters about the multiple training stages. 
}
\label{tab:training_stage_config}
\begin{tabular}{l|ccc}
\hline
Details & Initial Training & Fine-tuning Stage 1 & Fine-tuning Stage 2 \\
\hline
Training Crop Size & 384 & 640 & 640 \\
Train PAE Head & \xmark & \xmark & \cmark \\
Train Structure and Distogram & \cmark & \cmark & \xmark \\
Diffusion Batch Size & 48 & 32 & - \\
Train on Distillation Sets & \cmark & \cmark & \xmark \\
Bond Loss Weight & 0 & 1 & - \\
\hline
\end{tabular}
\end{table}
