## Supplementary material for "Protenix - Advancing Structure Prediction Through a Comprehensive AlphaFold3 Reproduction": figures and result summary: results.tex

\begin{figure}[tb]
    \centering
    \begin{tabular}{llll}
A. ~Ligands & B. ~~~~~~~~~~~Protein Interfaces & C. ~~~~~Nucleic Acids &
    D. ~~~~~RNA  \\
    \adjustbox{valign=t}{\includegraphics[height=0.24\textwidth]{figs/result/pb-summary.pdf}} &
    \adjustbox{valign=t}{\includegraphics[height=0.24\textwidth]{figs/result/prot-summary.pdf}}  &
    \adjustbox{valign=t}{\includegraphics[height=0.24\textwidth]{figs/result/nuc_prot.pdf}}  & 
 \multirow{2}{*}{\vspace{10em} \includegraphics[height=0.24\textwidth]{figs/result/casp15.pdf}} \\ %& \hspace{-8.5em}\raisebox{-1.8em}{\includegraphics[height=0.0675 \textwidth]{figs/result/figure_one_d_legend.pdf}} \\
    ~~~~~~~~\adjustbox{valign=t}{\includegraphics[width=0.085\textwidth]{figs/result/figure_one_a_legend.pdf}} & 
    ~~~~~~~~~~~~~~~~~~~~~~~~\adjustbox{valign=t}{\includegraphics[width=0.085\textwidth]{figs/result/figure_one_b_legend.pdf}} &
    \multicolumn{2}{c}{~~\adjustbox{valign=t}{ \includegraphics[width=0.21\textwidth]{figs/result/figure_one_c_legend.pdf}}~~~\adjustbox{valign=t}{\includegraphics[width=0.21\textwidth]{figs/result/figure_one_d_legend.pdf}}  }  \\
    % \multicolumn{4}{c}{ \includegraphics[width=0.95\textwidth]{figs/result/summary_legend.pdf} } \\
    \end{tabular}
    \caption{ \justifying \textbf{Model performance across several benchmarks.} \small Unless specified otherwise, performances are evaluated on the top-ranked prediction of each model based on its ranking score. \textbf{A.} Performance of \OurModelName{} and \AFShort{} on \PbShort{}. The RMSD success rate is the percentage of predictions where the pocket-aligned ligand RMSD is 2 \AA{} or lower. The PB-Valid success rate is the percentage of predictions that meet all 18 validity criteria specified by the PoseBusters test-suite.  $N$ is the number of structures. 
    \textbf{B \& C.} Performance of \OurModelName{}, \RFNAShort{}, \AFmultierShort{}, and \AFShort{} on filtered subsets of the Low Homology Recent PDB Set across different interface types. For protein interfaces, the success rate is defined as the percentage of interfaces with a DockQ score above 0.23, while for protein-nucleotide interfaces, the interface LDDT score is reported. Metrics are averaged within and across interface clusters; $N$ represents the number of clusters. "\AFShort{} in \AFShort{}" and "\RFNAShort{} in \AFShort{}" indicate metrics of \AFShort{} and \RFNAShort{} as reported by \AFShort{}, respectively.
    \textbf{D.} Performance of \OurModelName{}, AIChemy\_RNA2, \RFNAShort{}, and AF-3 on CASP15 RNA. $N$ represents the number of structures.
    }
    \label{fig:summary}
\end{figure}

\subsection{Overview}

\OurModelName{} is trained using experimental structures curated from the Protein Data Bank (PDB)~\cite{berman2000pdb} with a cutoff date of September 30, 2021 and predicted structures of protein monomers with AlphaFold2 \cite{jumper2021af2} and OpenFold \cite{ahdritz2024openfold} (See details in Table~\ref{tab:training-dataset} and Appendix~\ref{app:training-dataset}). We benchmark \OurModelName{} on different evaluation datasets to assess its performance on different types of molecules. Details of the evaluation datasets are provided in Appendix~\ref{app:evaluation-dataset}. 
We compare \OurModelName{}'s performance with that of \AFShort{}, \AFmultierShort{}, and \RFNAShort{}.


% In contrast to \AFShort{}, which trained two separate models—one using data with a cutoff of September 30, 2019, for evaluating the Posebusters benchmark set, and another with a cutoff of September 30, 2021, for all other evaluations—we opted to train a single model due to resource limitations. 
In contrast to \AFShort{}, which trains two separate models with different data cutoffs (September 30, 2019, for Posebusters, and September 30, 2021, for other datasets), we opt to train a single model due to resource limitations.
Targets from \PbFull{} (\PbShort{}) are excluded to avoid data leakage. We acknowledge the differences in the cutoff dates and present an analysis in Section~\ref{sec:ligands} to distinguish genuine improvements on Posebusters from potential data overlap.


For each PDB entry, we follow \AFShort{}'s inference setup, generating 25 predictions using 5 model seeds, with each seed producing 5 diffusion samples. The predictions are ranked using confidence scores. % We report the performance of the best, worst, and top-ranked predictions to provide a full view of model capabilities. Details of some hyperparameters such as the number of recycles, the size of MSA sequences used during inference are described in Appendix~\ref{app:evaluation}.

In the following sections, we present detailed benchmark results, along with in-depth analysis of low-performing cases to highlight potential directions for further improvement.


\begin{figure}[htbp]
\centering
\begin{tabular}{ll}
A. ~~~~~~~~~~~Overall Performance &
B. ~~~~~~~~Success rates with different RMSD thresholds
\\
\adjustbox{valign=t}{\includegraphics[width=0.375\textwidth]{figs/result/pb/overall_performance.pdf}} &
\adjustbox{valign=t}{\includegraphics[width=0.58\textwidth]{figs/result/pb/rmsd_with_curve.pdf}}
\\ %\vspace{2mm} \\
C. ~~~~Common ligands and others &
D. ~~~~~~~~~~~~~~~~Targets with different homology
\\
\adjustbox{valign=t}{\includegraphics[width=0.375\textwidth]{figs/result/pb/common-ligands.pdf}} &
\adjustbox{valign=t}{\includegraphics[width=0.58\textwidth]{figs/result/pb/prot-homology-detail.pdf}}
\\ %\vspace{2mm} \\
\multicolumn{2}{l}{E. ~~~~~~~~~~~~~~~~~~~~~~~~~~~~~~~~~~~~~~~~~~~~~~Detailed validity check comparison}
\\
\multicolumn{2}{c}{
\includegraphics[width=0.975\textwidth]{figs/result/pb/pb_valid_detailed.pdf}
} 
\end{tabular}
\caption{\justifying \textbf{PoseBusters evaluation.} \small \textbf{A.} Comparison of  protein-ligand binding success rate and the structure validity between \OurModelName{} and \AFShort{}. The metrics of \AFShort{} are computed using the PoseBusters test suite and the predictions provided by \AFShort{}. These metrics are consistent with the results reported by \AFShort{}. \textbf{B.} A comparison of RMSD success rates under different thresholds. In addition to the results for \AFShort{}, we present the corresponding success rates of our model under various sample selection methods. "Oracle" refers to the samples with the lowest RMSD; "all sample" presents the metrics computed using all 25 samples generated by \OurModelName{}; and "confidence ranked" provides the metrics for the samples chosen based on the confidence head. \textbf{C.} Comparison of results on ligands split into different categories. We follow \AFShort{}'s definition~\cite{abramson2024af3} to identify 50 \AFShort{} defined  Natural Common Ligands (NCL), which occur over 100 times in PDB and are visually identified as natural, and call them "old natural common ligands". Furthermore, we select other 75 NCL, referred as "new natural common ligands", 
%\hy{Add to appendix list} 
in \PbShort{} that meet the same criteria within our training set.  \textbf{D.} Comparison of results on targets with low, moderate, and high protein sequence homology. We display the homology computed with the 2019-cutoff training set (left) and the 2021-cutoff training set (right). Note that the sample size in each homology group is slightly different from \AFShort{} report ~\cite{abramson2024af3}, due to the potential differences in alignment methods used. %. More details can be found in appendix \ref{app:eval-PoseBusters} \cy{which section?}.
\textbf{E.} Comparison of detailed validity check on \PbShort{}. }
\label{fig:pb-main}
\end{figure}


\subsection{Ligands}
\label{sec:ligands}


% In this section, 
We evaluate \OurModelName{} on the PoseBusters Version 2 benchmark set~\cite{buttenschoen2024posebusters} and compare its performance to that of \AFShort{}~\cite{abramson2024af3}.  We follow the evaluation procedures outlined in \AFShort{} paper to ensure consistency and fairness of comparisons. Details of the evaluation procedure are provided in Appendix~\ref{app:evaluation}. 


\textbf{RMSD success rate.} The success rate is defined as the percentage of predictions for which the pocket aligned ligand \RmsdFull{} (\RmsdShort{}), as defined in ~\cite{abramson2024af3},
between the ground truth and the prediction is no greater than 2 \AA{}. As shown in Figure~\ref{fig:pb-main}[A], \OurModelName{} outperforms \AFShort{}-2019 in terms of success rate, achieving slightly higher rates for both RMSD and PB-Valid metrics. This suggests that \OurModelName{} represents the current state-of-the-art (SOTA) model for the protein-ligand cofolding task.

Figure~\ref{fig:pb-main}[B] shows success rates across varying RMSD thresholds, offering a more detailed view of model performance.
The \OurModelName{} - oracle configuration (red dashed line) achieves the highest success rate at all RMSD thresholds, consistently outperforming the \AFShort{}-2019 confidence ranked (black solid line). 
The \OurModelName{} - confidence ranked and \OurModelName{} - all sample configurations closely match \AFShort{}'s performance, highlighting \OurModelName{}'s ability to generate high-quality protein-ligand cofolding predictions. It's worth noting that the selected results still lag behind the best candidates among all samples, suggesting room for improvement with a better sample ranker. 

\textbf{Similarity analysis.} We note that \OurModelName{} is trained with PDB cut-off 2021-09-30, hence the improvement of performance may be attributed to additional training data. To avoid potential data leakage issue, we remove identical PDB IDs from our training set. Additionally, we perform a similarity analysis as shown in Figure \ref{fig:pb-main}[C] and [D]. These figures illustrate that compared to \AFShort{}-2019, \OurModelName{} performs better on common ligands, likely benefiting from the additional training data. However, since they exhibit the same performance on non-common ligands, we can conclude that our model generalizes as well as \AFShort{}-2019. On the target side, \OurModelName{} outperforms \AFShort{}-2019 on high homology group but performs slightly worse on other targets.


\begin{figure}[h]
\centering
\begin{tabular}{ll}
% A. & B. \\
\raisebox{0.em}{A.}~~~~~~~~~~\adjustbox{valign=t}{\includegraphics[width=0.35\textwidth]{figs/result/case_study/ligand_good_case_7wux_v3.png}}~~~~~~ &
\raisebox{0.em}{B.}~~~~~~~~~~\adjustbox{valign=t}{\includegraphics[width=0.35\textwidth]{figs/result/case_study/ligand_good_case_7urd_v2.png}}~~~~~~~~
\\ 
\end{tabular}
\caption{\justifying \textbf{Novel protein-ligand interactions.} \small Predicted protein chains are shown in {\color{myblue}{blue}}, predicted ligands show in {\color{myturquoise}{turquoise}}, and the ground truth is shown in {\color{mygray}{gray}}. \textbf{A.} AziU3/U2 complexed with (5S,6S)-O7-sulfo DADH from Streptomyces sahachiroi (PDB: 7WUX). \textbf{B.} Human PORCN in complex with LGK974, a clinical-stage inhibitor with 30 heavy atoms (PDB: 7URD).}
\label{fig:ligand_good_case}
\end{figure}


\textbf{Physical plausibility.} We also conduct the \PbShort{} plausibility test, using the PoseBusters test-suite version 0.2.7 \cite{buttenschoen2024posebusters}. Apart from \RmsdShort{} metrics, PB-valid reflects the ability of a model to generate physically valid structures. As shown in Figure \ref{fig:pb-main}[E], we check on 18 validity criteria, which can primarily be categorized into intramolecular validity, intermolecular validity, chemical validity and consistency, etc.  Similar to \AFShort{}, the model cannot consistently generate samples with correct chirality and without steric clashes, hence we follow \AFShort{} and incorporate a penalty term to mitigate such stereochemistry violations in our score function when we perform sample ranking.  This penalty term has successfully improved the PB-valid success rate by over 10\%, though there still exists a roughly 5\% chance that we fail to predict the correct chirality.


\begin{figure}[!t]
\centering
\begin{tabular}{ll}
\multicolumn{2}{l}{
\raisebox{10em}{A.}~~~~~ \includegraphics[width=0.85\textwidth]{figs/result/case_study/ligand_bad_case_7loe_cropped.png}} \\
\raisebox{0.em}{B.}  \adjustbox{valign=t}{\includegraphics[width=0.46\textwidth]{figs/result/case_study/ligand_bad_case_5sak_1_v2.png}} &
\adjustbox{valign=t}{\includegraphics[width=0.46\textwidth]{figs/result/case_study/ligand_bad_case_5sak_2_v2.jpg}}
\\ 
\end{tabular}
\caption{\justifying \textbf{Sensible failure cases.} \small Predicted protein chains are shown in {\color{myblue}{blue}}, predicted ligands show in {\color{myturquoise}{turquoise}}, and the ground truth is shown in {\color{mygray}{gray}}. \textbf{A.} T4 lysozyme mutant L99A in complex with 1-fluoranylnaphthalene alternative ligand conformations (PDB: 7LOE). \textbf{B.} Endothiapepsin in complex with compound FU5-1 (PDB: 5SAK) and similar fragment in the PDB for the same target (PDB: 5SAN). }
\label{fig:ligand_fail_case}
\end{figure}

\textbf{Ligand case studies.} We inspect some challenging cases to assess \OurModelName{}’s ability to predict protein-ligand interactions. In cases where there are no close structural neighbors in the training set, \OurModelName{} demonstrates highly accurate predictions, as shown in Figure \ref{fig:ligand_good_case}. Specifically, Figure \ref{fig:ligand_good_case}[A] shows precise binding interactions with endogenous ligands like sulfooxy-heptanoic acid (7WUX), while Figure \ref{fig:ligand_good_case}[B] illustrates accurate predictions for novel inhibitors (7URD), even when the ligands are larger and have tight binding conformations. These examples highlight \OurModelName{}’s ability to predict diverse and complex interactions with high accuracy.
% a novel inhibitors (),  (7URD) even when ligand size is large (30 heavy atoms) and binding conformation is tight.  

We also inspect the cases where the RMSD of predicted ligands exceeds 2 \AA{} and find that, in some instances, the predictions are reasonable. For example, in Figure \ref{fig:ligand_fail_case}[A], \OurModelName{} predicts an alternative ligand conformation with the fluorine atom oriented oppositely (7LOE), though both conformations share similar occupancies (42\% vs. 58\%). In Figure \ref{fig:ligand_fail_case}[B], \OurModelName{}’s prediction results in a larger RMSD, but the predicted binding mode still maintains a key interaction between the ligand and two aspartates (5SAK). Interestingly, the predicted binding mode resembles another fragment in the PDB for the same target (5SAN). These cases reveal areas for improvement in the evaluation metrics. They also show that \OurModelName{} is able to capture key binding features even in more challenging predictions.


\subsection{Proteins}


We benchmark our model against \AFmultierShort{}, the top-performing open-source model available for comparison. Both models predict the first bioassembly of each structure. For \AFmultierShort{}, non-protein chains are excluded, and non-standard residues are mapped to standard ones, as the model does not support these components. Our model predicts the complete complex, though we do not assess whether including additional chains in the input influences performance.  The results are summarized in Figure~\ref{fig:summary}[B]. We report the DockQ success rate for different types of interfaces: all protein-protein interfaces and protein-antibody interfaces.
We report results from our model with 10 recycles and from \AFmultierShort{}(5x5) with 20 recycles, including separate scores for best (oracle), median, and top-1 ranked predictions. For \OurModelName{}, the ranker used for protein interfaces is the "chain pair ipTM" confidence. Overall, our model achieves higher DockQ success rate compared to \AFmultierShort{}, indicating improved prediction accuracy. The only exception occurs in protein-antibody cases, where our top-ranked predictions are on par with those of \AFmultierShort{}, indicating potential for improvement in the sample ranker.


\textbf{Case study.} Beyond the performance metrics, we also identify an interesting case, as shown in Figure~\ref{fig:protein_application}. It  highlights the potential applicability of \OurModelName{} as a tool for biology and pharmacology research, helping with hypothesis testing and uncovering novel mechanisms which can serve as the starting point for the development of new therapeutics. 

\begin{figure}[H]
\centering
\vspace{0.5cm}
\begin{tabular}{ll}
     \adjustbox{valign=t}{\includegraphics[width=0.48\textwidth]{figs/result/case_study/protein_good_case_7xva_v3.jpg}} & 
     \adjustbox{valign=t}{\includegraphics[width=0.48\textwidth]{figs/result/case_study/protein_dna_good_case_7xv6_v2.png}} \\
\end{tabular}
\caption{\justifying \textbf{Potential applications.} \textbf{Left}: \small \OurModelName{} prediction ({\color{myblue}{blue}} and {\color{myturquoise}{turquoise}}) on a novel protein-protein interaction from PDB: 7XVA ({\color{mygray}{gray}}), only very recently characterized as novel regulation mechanism for the {\color{myblue}{TR4 transcription factor}} binding to {\color{myturquoise}{JAZF1}}, a regulation mechanism involved in different cancer types, whose understanding might lead to novel therapeutic developments. \textbf{Right}: Notably, \OurModelName{} also predicts correctly the TR4 DBD binding to DNA as homodimer from PDB: 7XV8, highlighting integrative modeling capabilities and ability to model complex biological mechanisms ~\cite{liu2023structures}.
% [[cite: https://doi.org/10.1093/nar/gkac1259]]
}
\label{fig:protein_application}
\end{figure}


\subsection{Nucleic Acids}


We extend our evaluations to include RNA and DNA targets, where \OurModelName{} demonstrates performance comparable to \AFShort{} while achieving higher accuracy than \RFNAShort{}. \OurModelName{} does not use MSA for nucleic chains.

\begin{figure}[t!]
    \centering
    \begin{tabular}{ll}
        A.~~~~~~~~~~~~~~~~~~~~~~~~~~~LDDT & B.~~~~~~~~~~~~~~~~~~~~~~~~~~~TM-score \\
        \includegraphics[width=0.48\linewidth]{figs/summary/casp_lddt_compare_summary.pdf} &  \includegraphics[width=0.45\linewidth]{figs/summary/casp_tm_compare_summary.pdf} \\
    \end{tabular}
    \caption{ \justifying \textbf{CASP15 RNA prediction accuracy.} \small The results from AlChemy\_RNA2, \RFNAShort{}, \AFShort{}, and \OurModelName{} are reported.}
    \label{fig:casp-rna}
\end{figure}


\textbf{CASP15.} We assess \OurModelName{} on CASP15 RNA targets, following the approach in the \AFShort{} paper, focusing on eight targets that were publicly available as of December 1, 2023.  We compare the results of \OurModelName{}, \AFShort{}, AIchemy\_RNA2, and \RFNAShort{}. The AIchemy\_RNA2 predicted structures are obtained from the CASP website, with those exhibiting the highest LDDT scores selected for analysis.  The \RFNAShort{} predicted structures are downloaded from the Zenodo repository~\cite{baek23casp15}, alongside the original paper.  The metrics for \AFShort{} are taken from the preprint by Bernard et al.~\cite{bernard24hasalphafold3}, where the authors systematically evaluated \AFShort{}'s RNA prediction performance and reported LDDT and TM-score values computed using OpenStructure~\cite{mariani2013lddt,biasini2013openstructure}. Since the authors evaluted \AFShort{}'s performance based on a single seed, we also report the performance of \OurModelName{} using a single seed instead of five seeds. The top-ranked sample of each structure, selected based on pLDDT, is used for evaluation. 
For \RFNAShort{}, AIchemy\_RNA2, and \OurModelName{}, the LDDT and TM-score metrics are recomputed consistently using OpenStructure~\cite{mariani2013lddt,biasini2013openstructure}. As shown in Figure~\ref{fig:summary}[C] and Figure~\ref{fig:casp-rna}, the average LDDT and TM-score of \OurModelName{} are similar to those of \AFShort{}, significantly outperforming \RFNAShort{}, but still lagging behind AIchemy\_RNA2~\cite{chen2023rna}, which benefits from human input.
\begin{wrapfigure}{r}{0.35\textwidth}
\centering
%\vspace{-0.5cm}
\adjustbox{valign=t}{\includegraphics[width=0.35\textwidth]{figs/result/case_study/nuc_good_case_7r6r_v2.png}} 
\caption{\justifying \textbf{Novel protein-nucleic acid complex.} \small Accurate prediction ({\color{myblue}{blue}} and {\color{myturquoise}{turquoise}}) by \OurModelName{} on a novel protein-nucleic acid complex (mycobacteriophage immunity repressor protein bound to double stranded DNA) without close homologs in the training set. (PDB: 7R6R ({\color{mygray}{gray}})) }
\label{fig:nuc_good_case}
\vspace{-1cm}
\end{wrapfigure}


\textbf{Protein-nucleic acid complexes.}  As illustrated in Figure~\ref{fig:summary}[C][D], \OurModelName{} predicts protein-nucleic complexes with higher accuracy than \RFNAShort{}. To ensure a fair comparison, we apply the same evaluation approach to both  \OurModelName{} and \RFNAShort{} (details in Appendix~\ref{app:evaluation}). For protein-dsDNA interfaces, the two DNA chains in the double-stranded DNA (dsDNA) are treated as a single entity. The interface LDDT is then calculated between this entity and the protein chain.   
The dataset used for our comparison is curated based on the description provided in \AFShort{} (See Appendix~\ref{app:evaluation-dataset} for details). The number of structures in our curated dataset matches the count reported in \AFShort{}'s Figure 1.c, although we cannot guarantee the datasets are identical. Notably, the performance of \RFNAShort{} differs from that reported in \AFShort{}, which may be attributed to discrepancies in the dataset or the evaluation tools used. For each structure, the top-ranked sample based on the complex ranking score ($0.8 \cdot \text{ipTM} + 0.2 \cdot \text{pTM} - 100 \cdot \text{has\_clash}$)
is selected for evaluation. This is because, for dsDNA-protein interfaces, three chains are involved, making it difficult to use chain pair scores directly.  

%The \AFShort{} report does not directly describe the calculation method for dsDNA-Protein iLDDT, so this discrepancy may lead to differences in the metrics. 


\textbf{Case study.} We evaluate \OurModelName{}’s ability to predict protein-nucleic acid complexes, with a particular focus on cases with novel structures. One such example is shown in Figure~\ref{fig:nuc_good_case}, where \OurModelName{} successfully predicts a novel protein-DNA complex that reveals the mechanism by which a monomeric repressor promotes transcriptional silencing.


\subsection{Discussion}


\paragraph{Confidence score.}
We make slight modifications to the confidence head architecture in contrast to \AFShort{}, which will be discussed in Section \ref{sec:method}.
After training the confidence head, we observe that predicted scores can be used to select interfaces on low homology recentPDB dataset. 
As illustrated in Figure \ref{fig:chain_iptm_dockq}, a positive Pearson correlation ($\rho$>0.5) is observed between predicted scores and interface DockQ scores for diverse interface types, including protein-protein and protein-nucleic acid interactions.
Given that our PAE head has been trained for only 4K steps, there is still huge potential for further improvement.

\begin{figure}[h]
    \centering
    \begin{tabular}{lll}
        A. ~~~~~~~~~~~~~All Interface & B.~~~~~~~~~~~Protein-Protein & C.~~~~~~~Protein-Nucleic Acid \\
         % \raisebox{4em}{\rotatebox{90}{\tiny LDDT}}  
         \includegraphics[width=0.3\linewidth]{figs/dockq_iptm/all_chain_iptm_dockq.pdf}  &  \includegraphics[width=0.3\linewidth]{figs/dockq_iptm/prot_prot_chain_iptm_dockq.pdf} &  \includegraphics[width=0.3\linewidth]{figs/dockq_iptm/nuc_prot_chain_iptm_dockq.pdf}  \\
    \end{tabular}
    \caption{\justifying \textbf{Relationship between predicted chain pair ipTM and the DockQ score.} \small \textbf{A}. A box-and-whisker plot of DockQ against binned chain pair ipTM on both protein-protein and protein-nucleic acid interfaces.
    \textbf{B}. The subset of figure \textbf{A} on the protein-protein interfaces. \textbf{C}. The protein-nucleic acid subset of figure A.}
    \label{fig:chain_iptm_dockq}
\end{figure}


\paragraph{Limitations.}

We review the showcase examples presented in the \AFShort{} paper to evaluate how our model performs on these cases. We find that our model also predicts these structures with high accuracy. Upon observing similar results, we further investigated the training set and noted that some of these complexes share significant similarities with certain training samples. For instance, as shown in Figure~\ref{fig:similar_case}, the protein-DNA complex 7PZB has a close counterpart in the training set (3MZH), as does the protein-ligand complex 7XFA, which is similar to 5OAX. This suggests that the accuracy of both \OurModelName{} and \AFShort{} may partially result from memorization, indicating the need for more out-of-distribution (OOD) test sets to better assess its generalizability.


\begin{figure}[h]
\centering
\begin{tabular}{llll}
A. \adjustbox{valign=t}{\includegraphics[width=0.25\textwidth]{figs/result/case_study/limitation_na_7pzb_1.png}} & \hspace{-10pt}
\adjustbox{valign=t}{\includegraphics[width=0.24\textwidth]{figs/result/case_study/limitation_na_7pzb_2.png}}
& 
B. \adjustbox{valign=t}{\includegraphics[width=0.17\textwidth]{figs/result/case_study/limitation_pl_7xfa_2_v3.png}} &
\adjustbox{valign=t}{\includegraphics[width=0.185\textwidth]{figs/result/case_study/limitation_pl_7xfa_1_v3.png}}
\\ 
\end{tabular}
\caption{\justifying \textbf{} \small \textbf{A.} \textbf{Left}: Example of good prediction of protein-nucleic acids interactions by \OurModelName{} ({\color{myblue}{blue}}) and PDB crystal structure 7PZB ({\color{mygray}{gray}}). \textbf{Right}: Close sample in the training set (3MZH) in {\color{mysalmon}{salmon}}. \textbf{B.} \textbf{Left}: Example of good prediction of protein-ligand interactions by \OurModelName{} in PoseBusters and PDB crystal structure (7XFA). \textbf{Right}: Close sample in the training set (5OAX). }
\label{fig:similar_case}
\end{figure}
