## Supplementary figures and images for "Protenix - Advancing Structure Prediction Through a Comprehensive AlphaFold3 Reproduction"

### all_chain_iptm_dockq.pdf

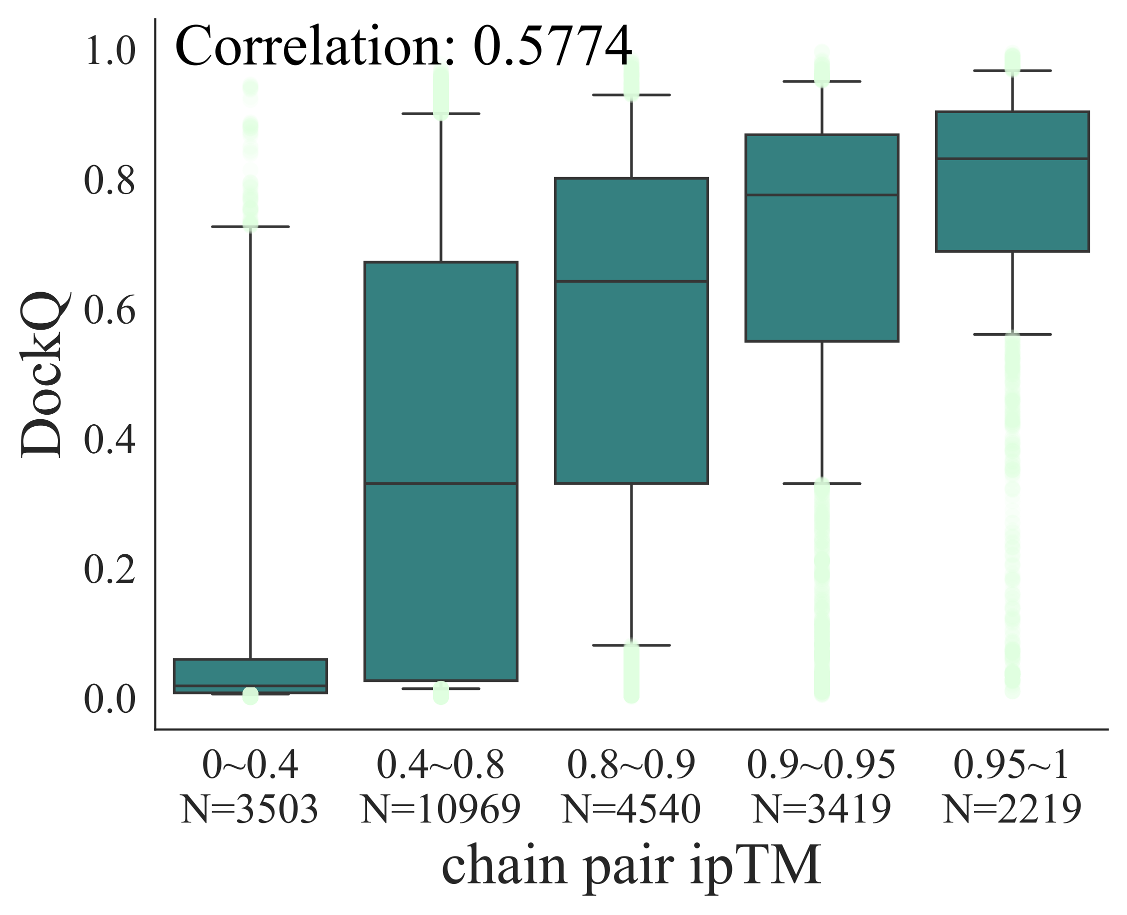

### bytedance_logo.png

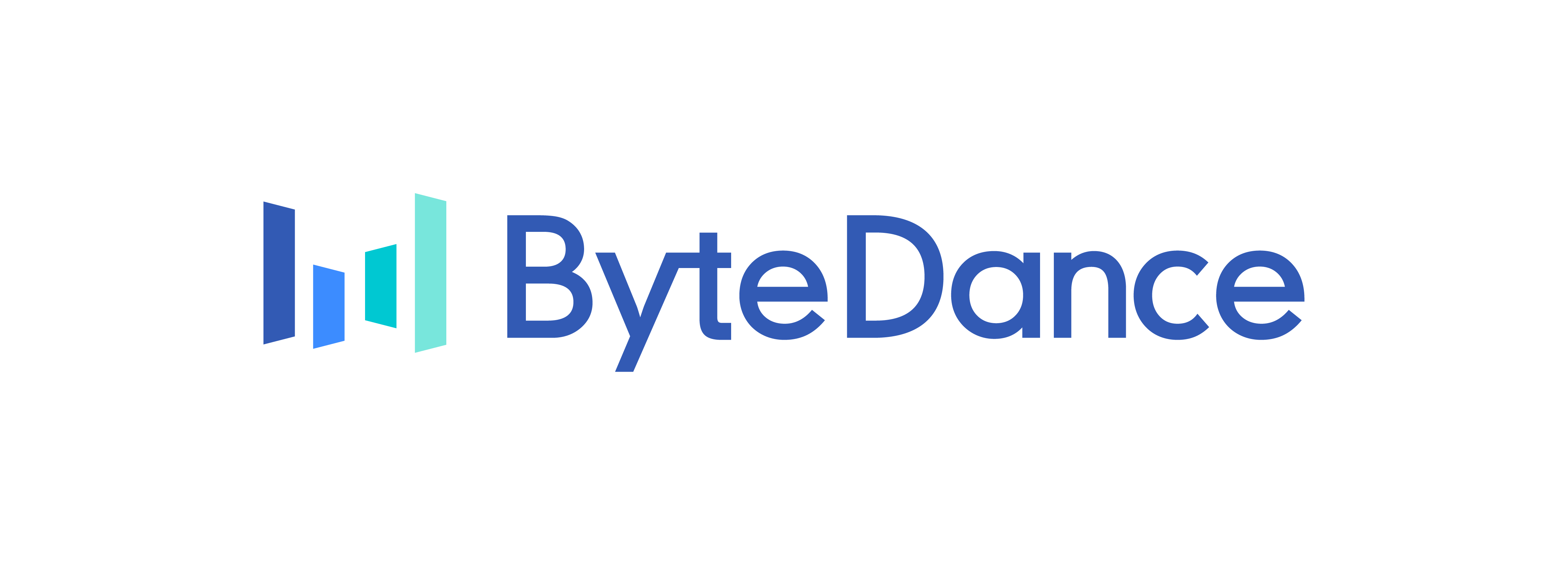

### bytedance_logo_crop.png

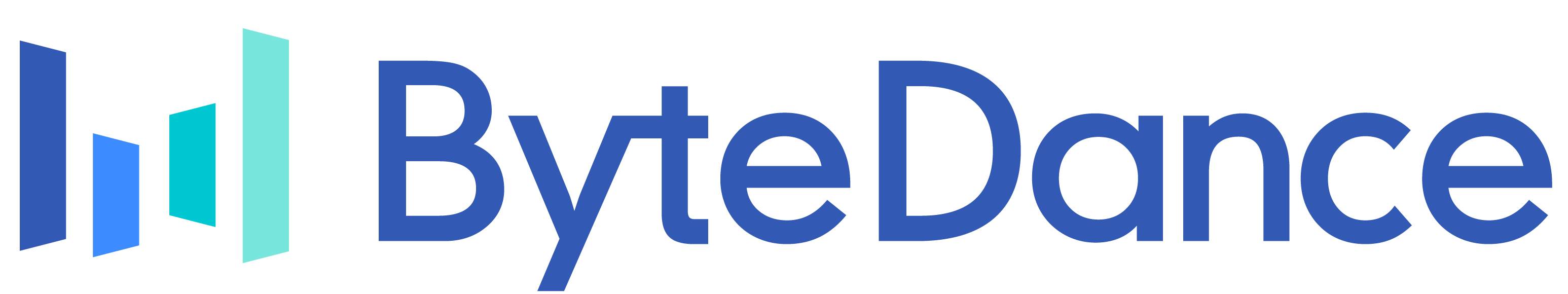

### casp15.pdf

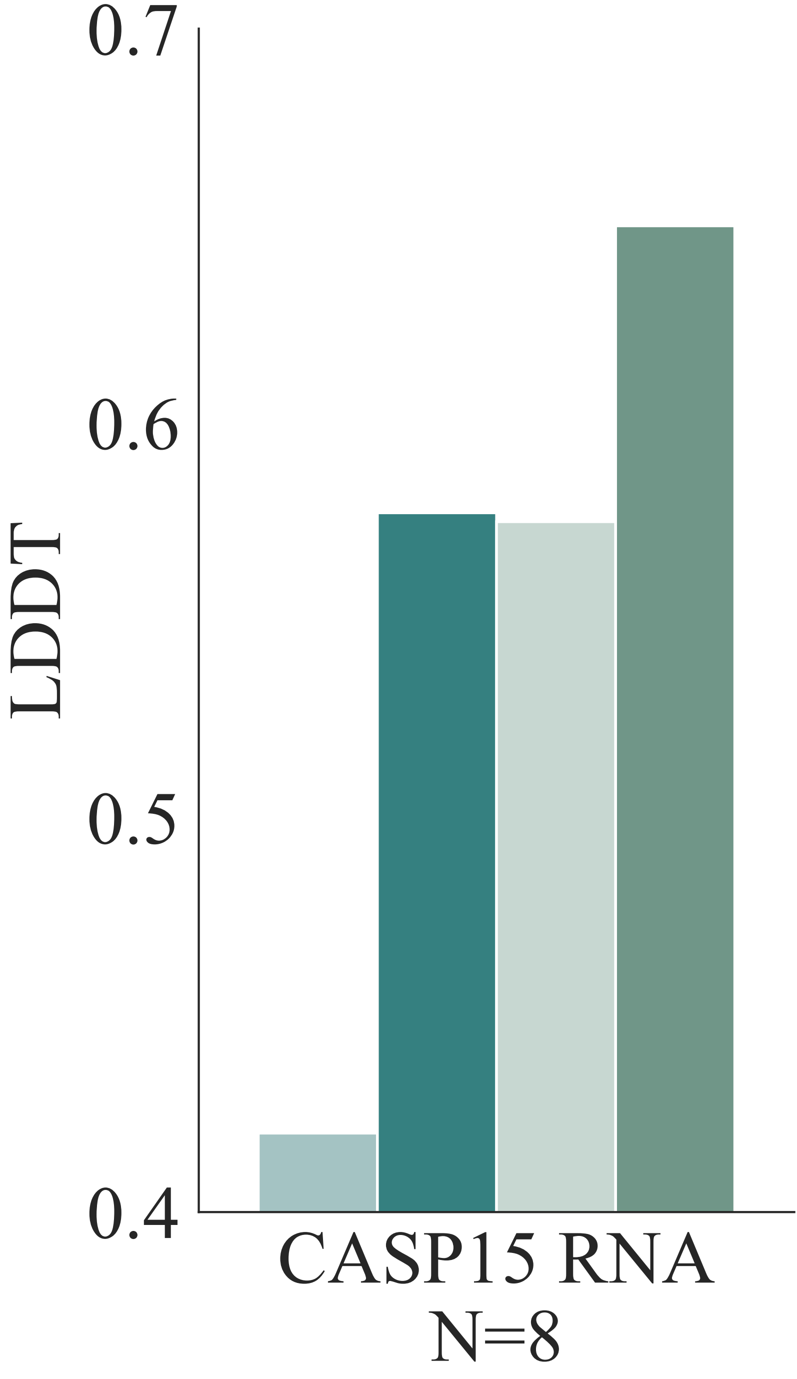

### casp_lddt_compare_summary.pdf

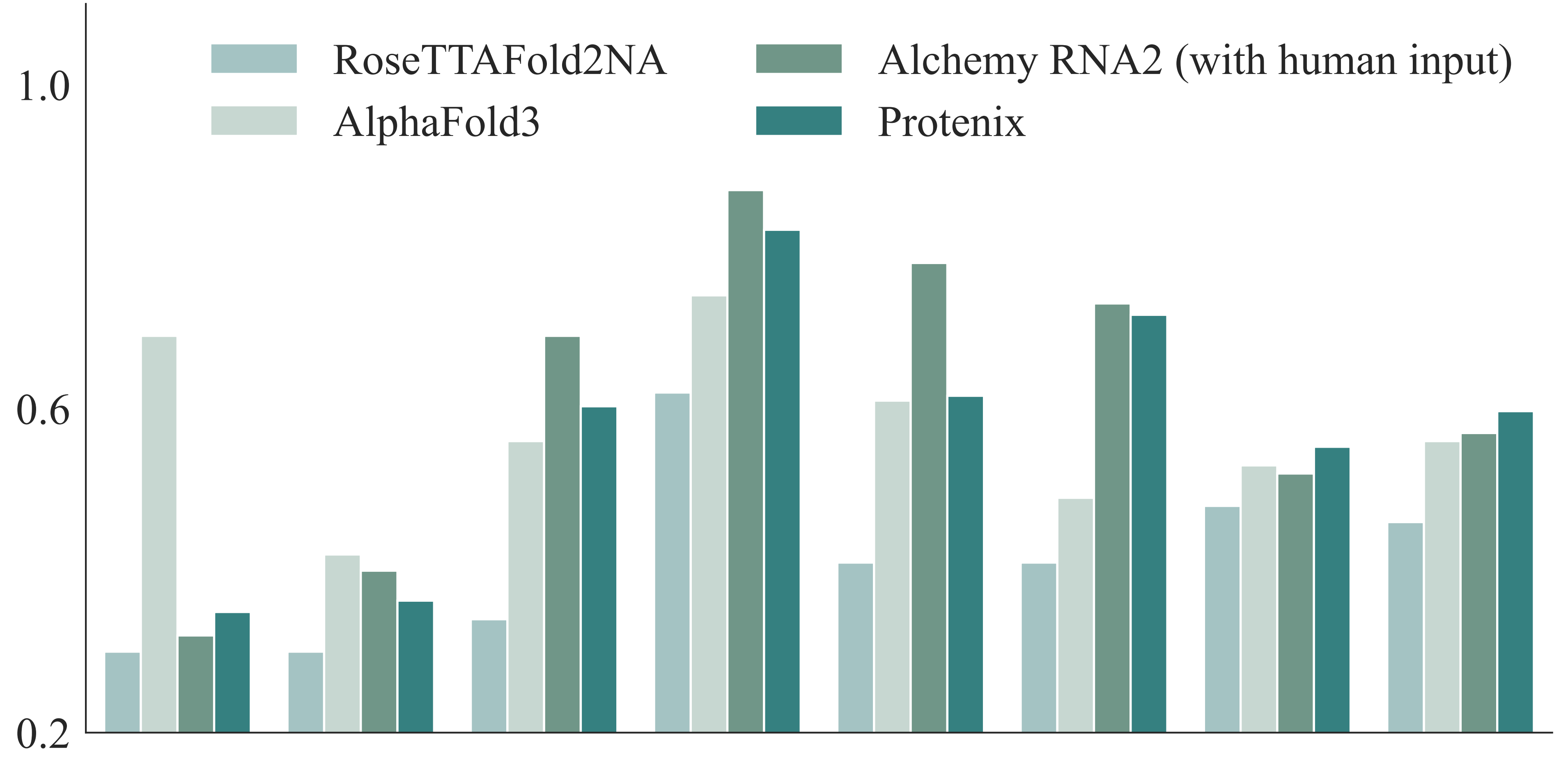

### casp_tm_compare_summary.pdf

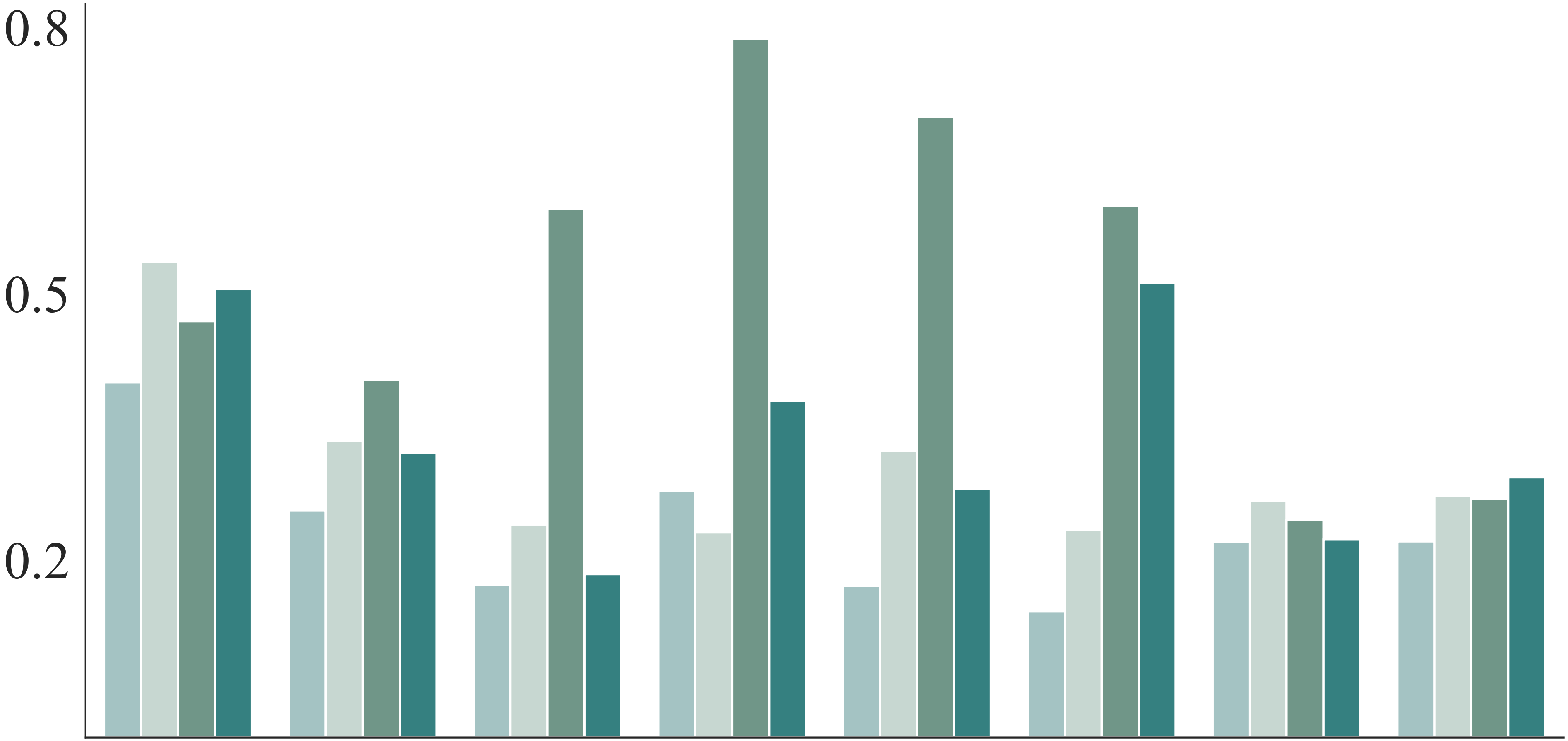

### common-ligands.pdf

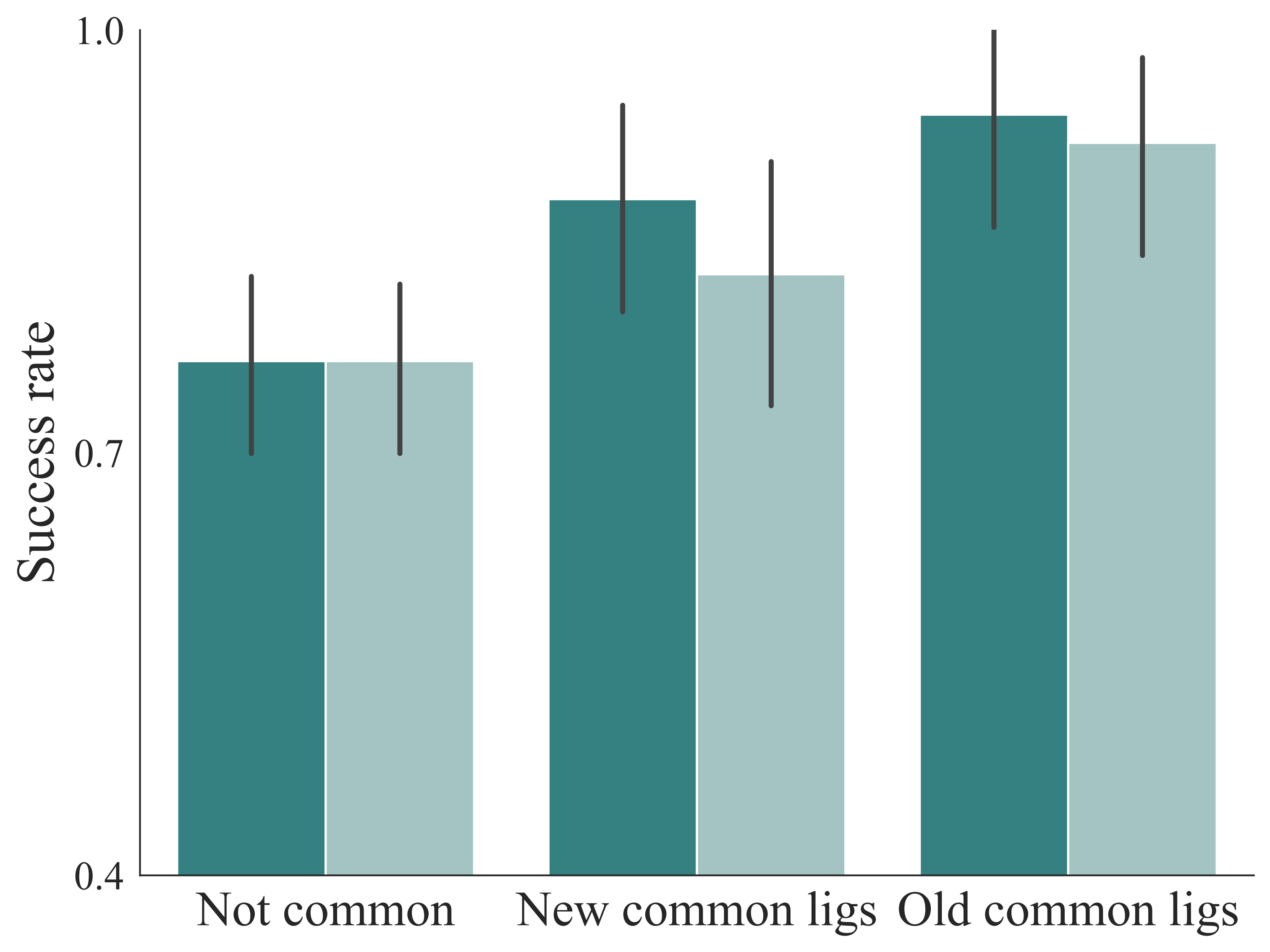

### dockq_compare_summary.pdf

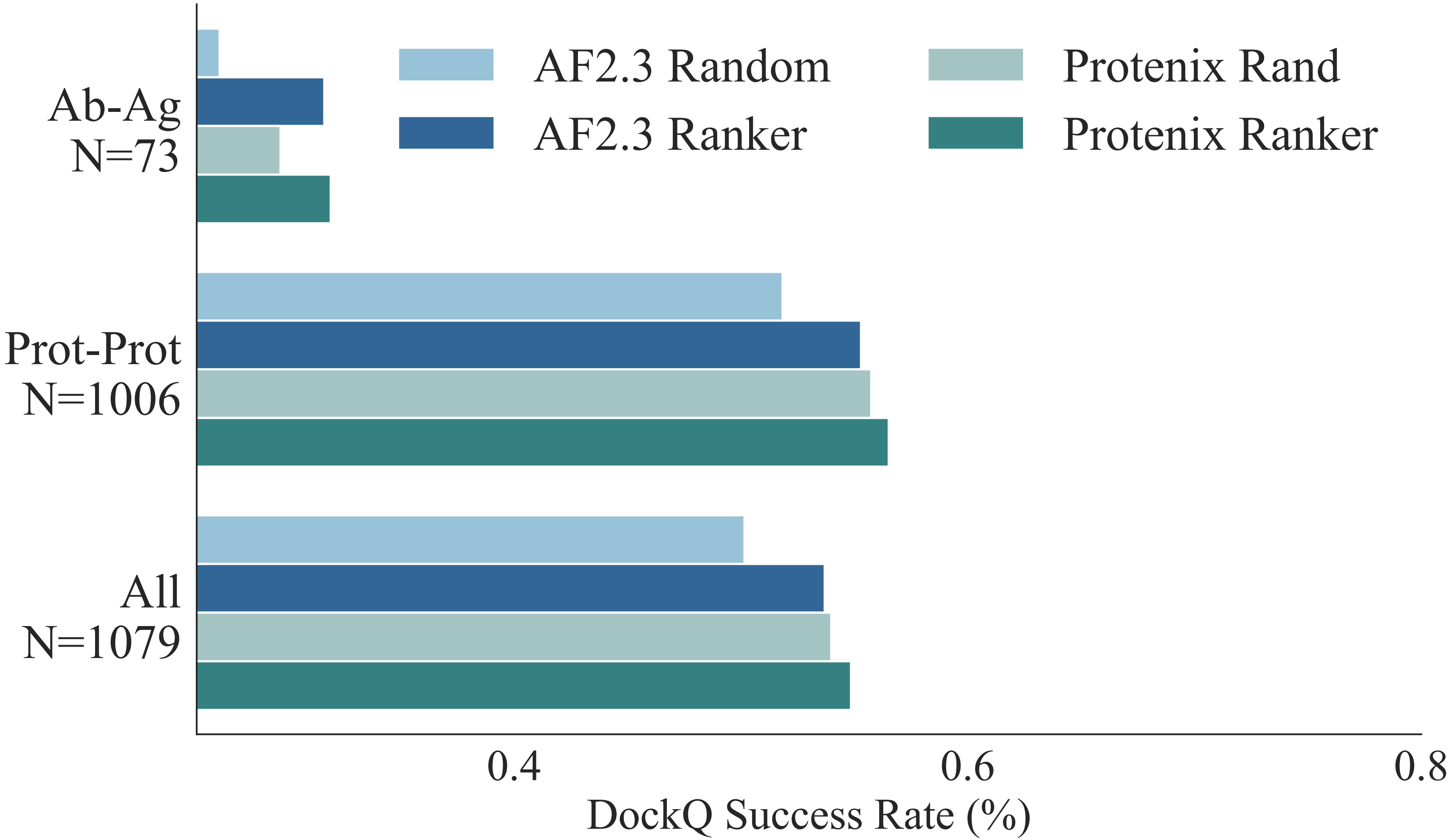

### figure_one_a_legend.pdf

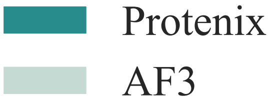

### figure_one_b_legend.pdf

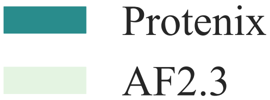

### figure_one_c_legend.pdf

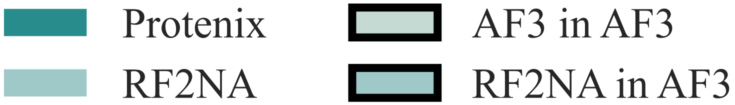

### figure_one_d_legend.pdf

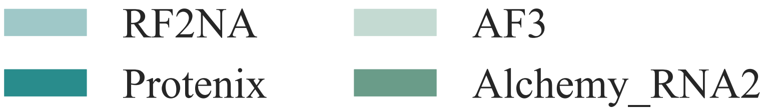

### ligand_bad_case_5sak_1_v2.png

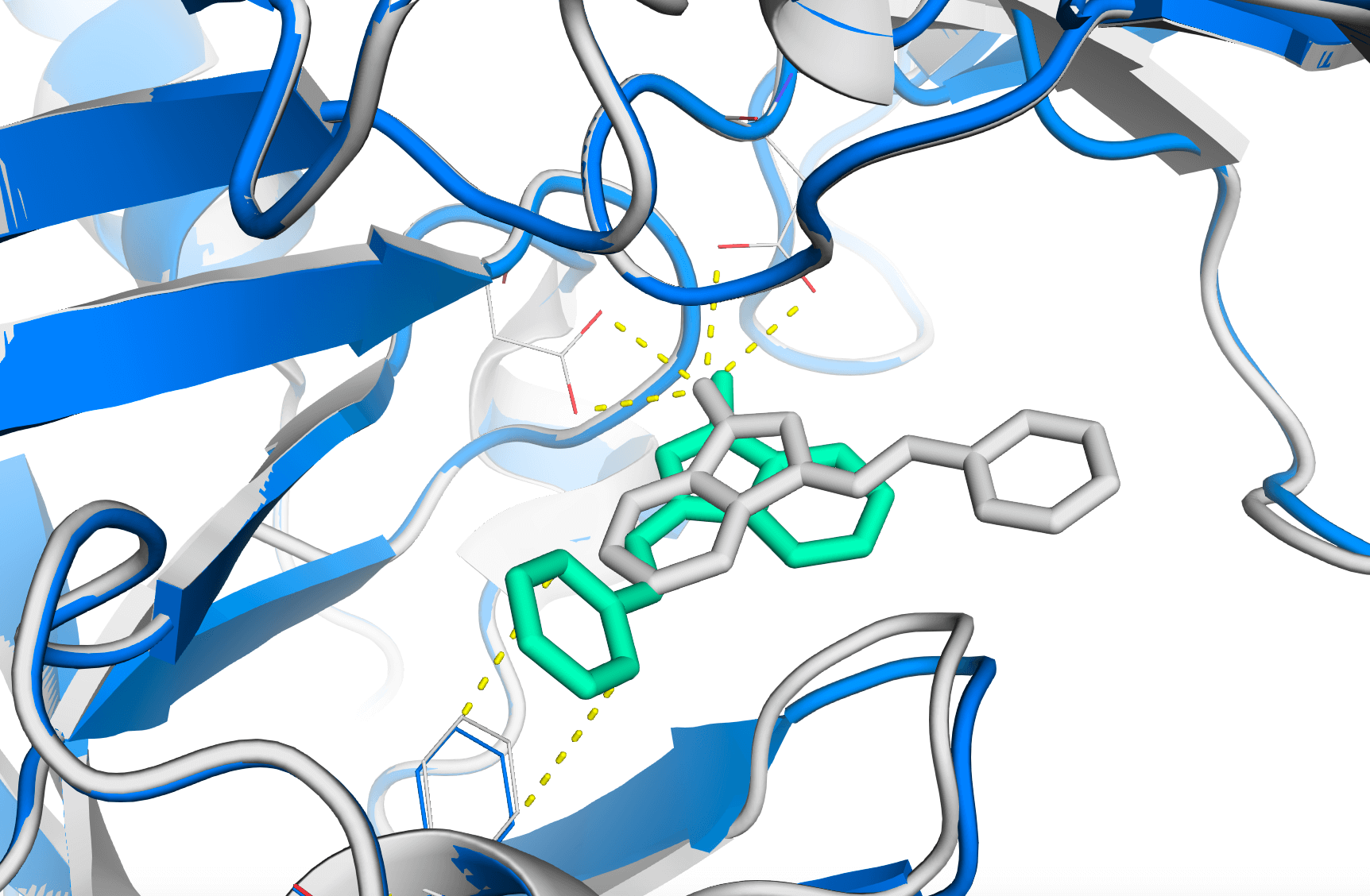

### ligand_bad_case_5sak_2_v2.jpg

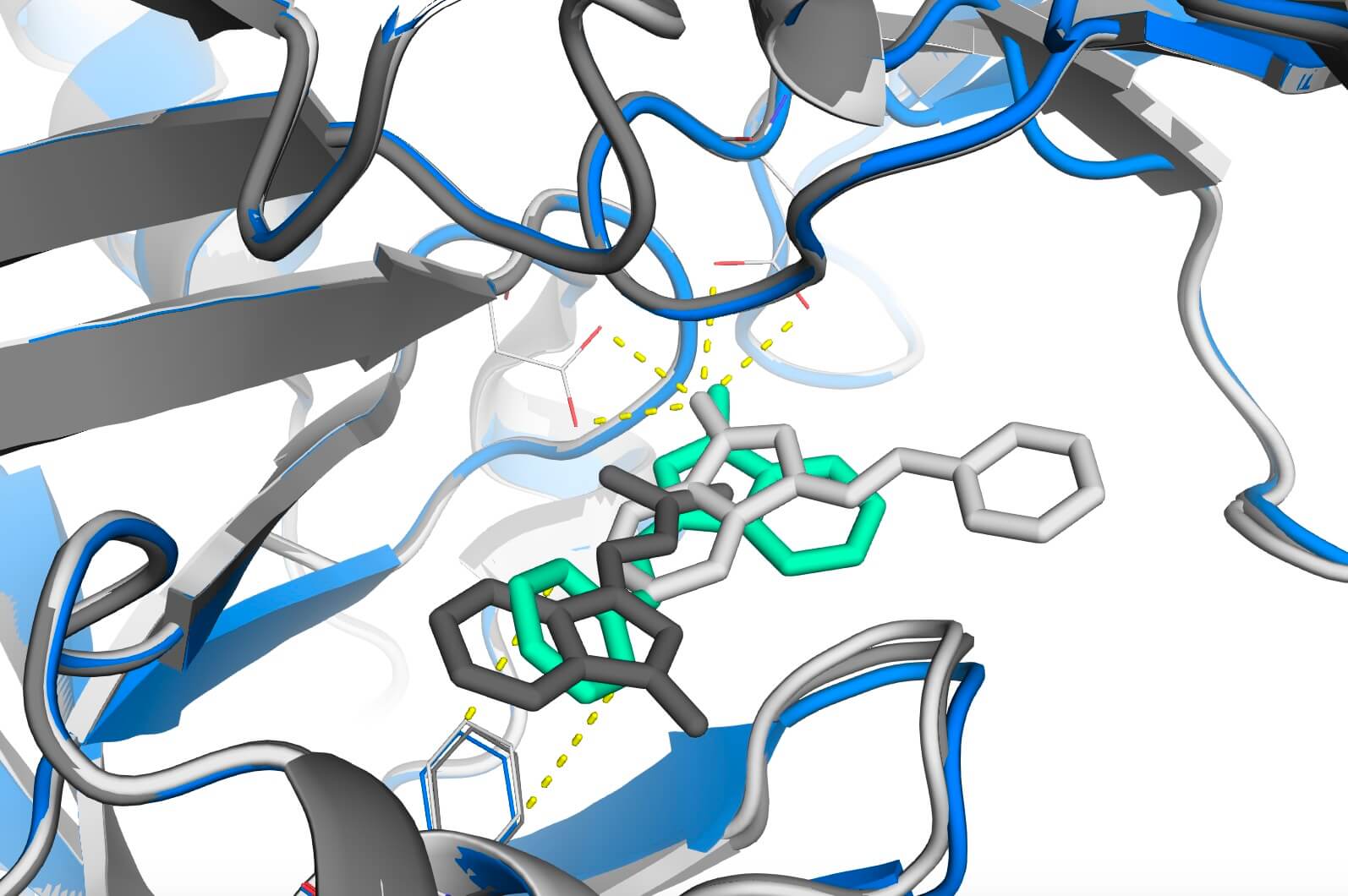

### ligand_bad_case_7loe_cropped.png

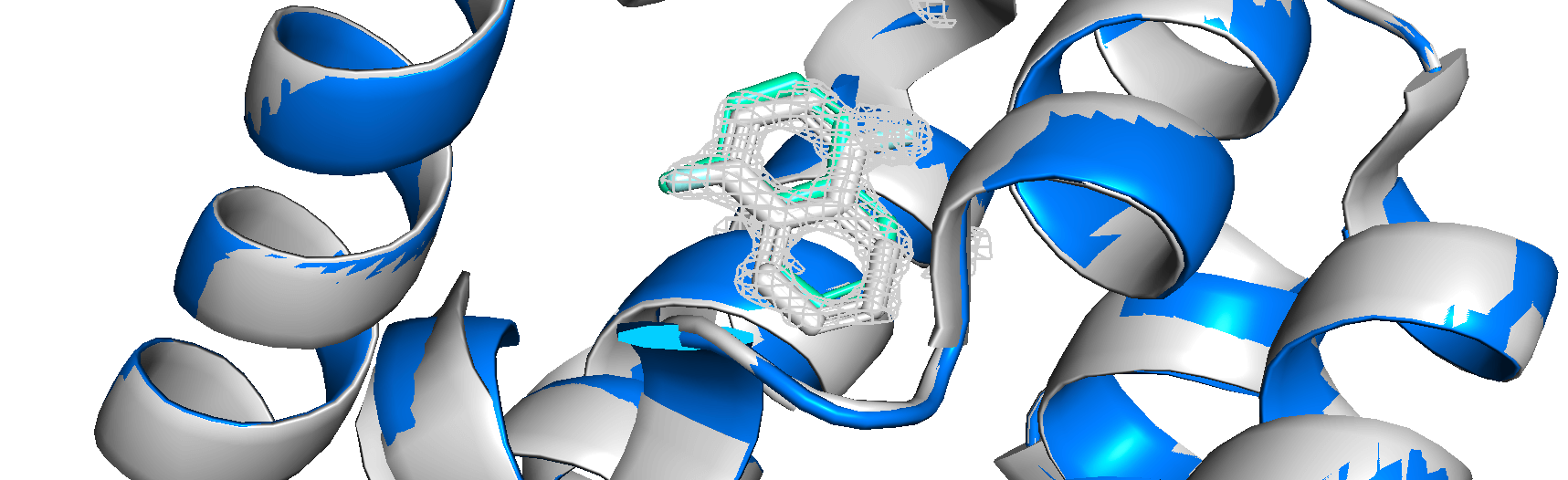

### ligand_good_case_7urd_v2.png

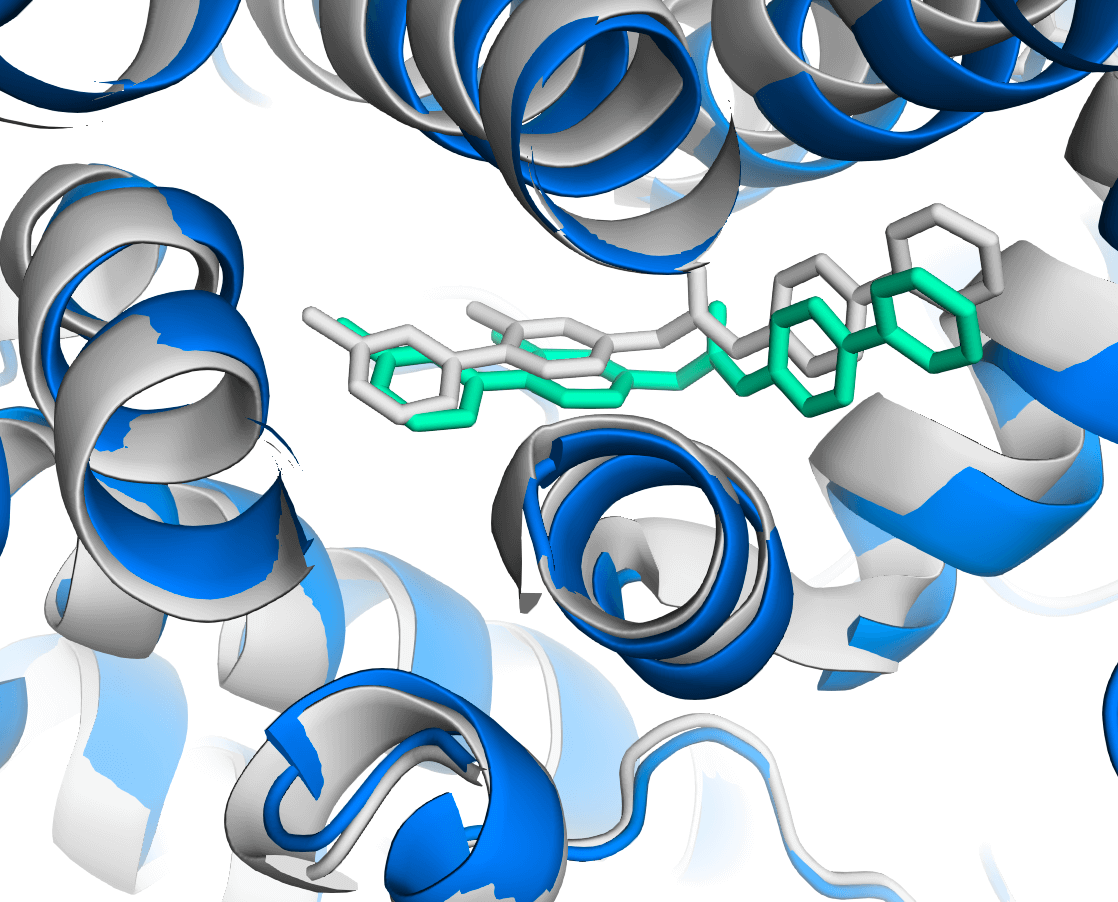

### ligand_good_case_7wux_v3.png

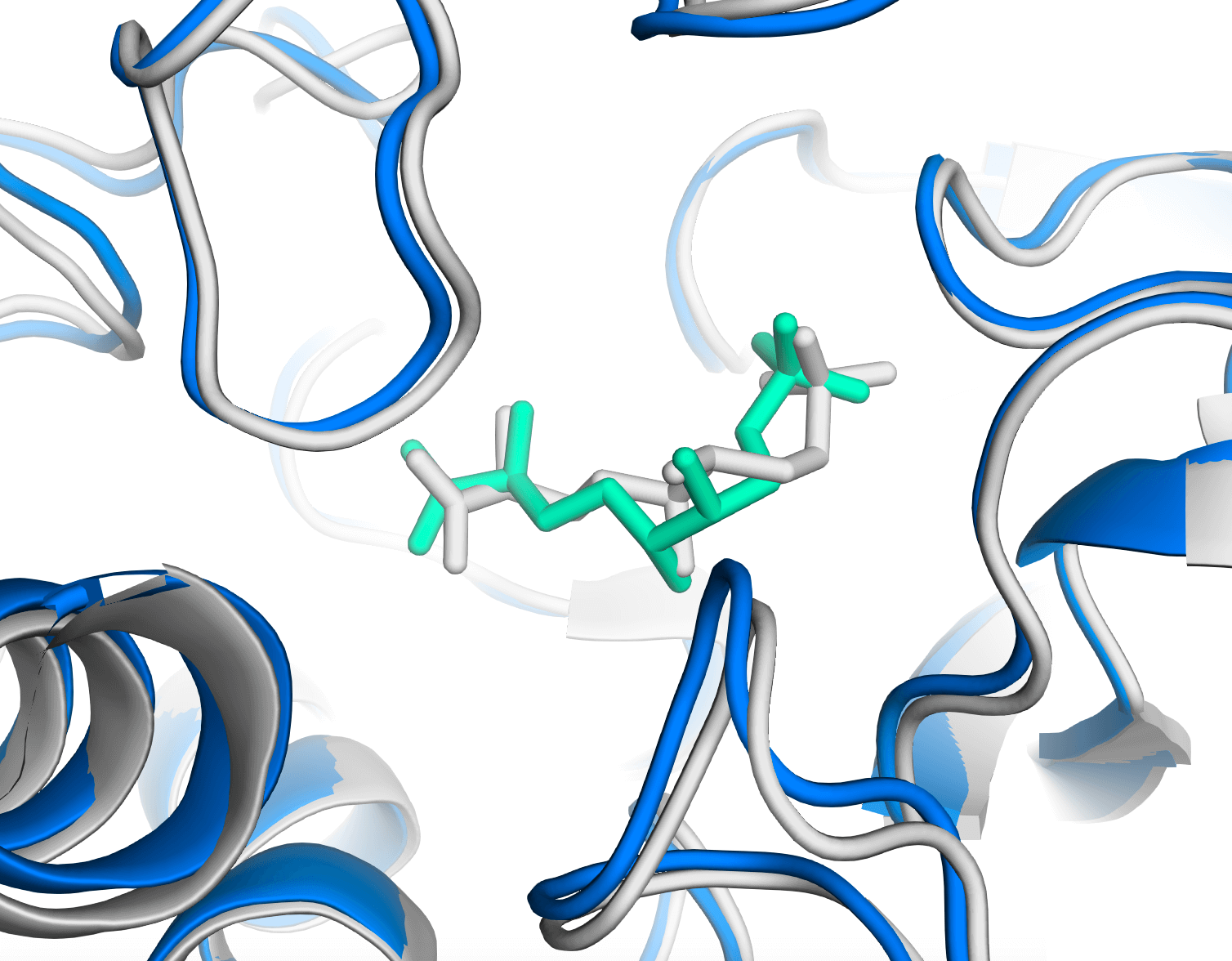

### limitation_na_7pzb_1.png

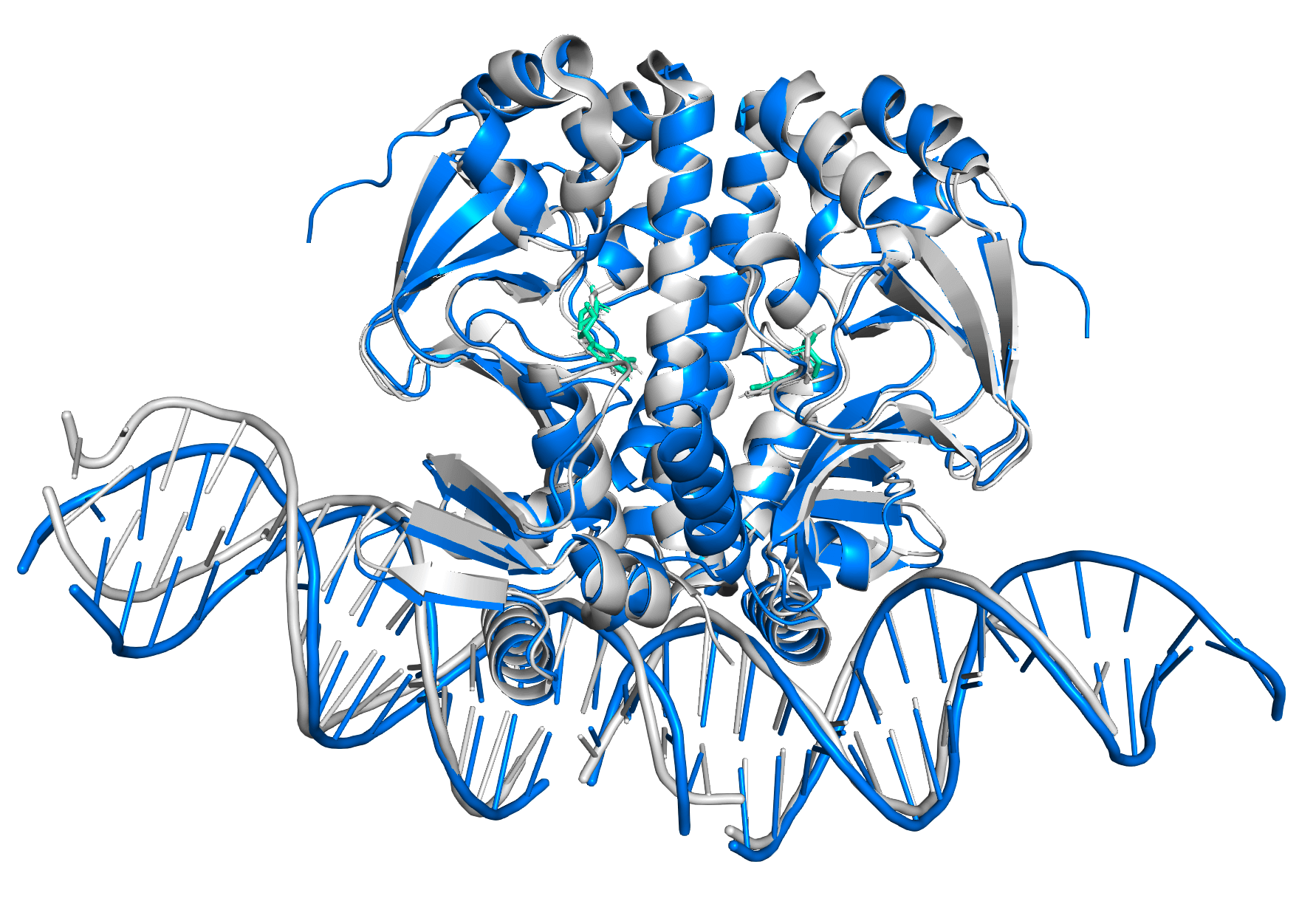

### limitation_na_7pzb_2.png

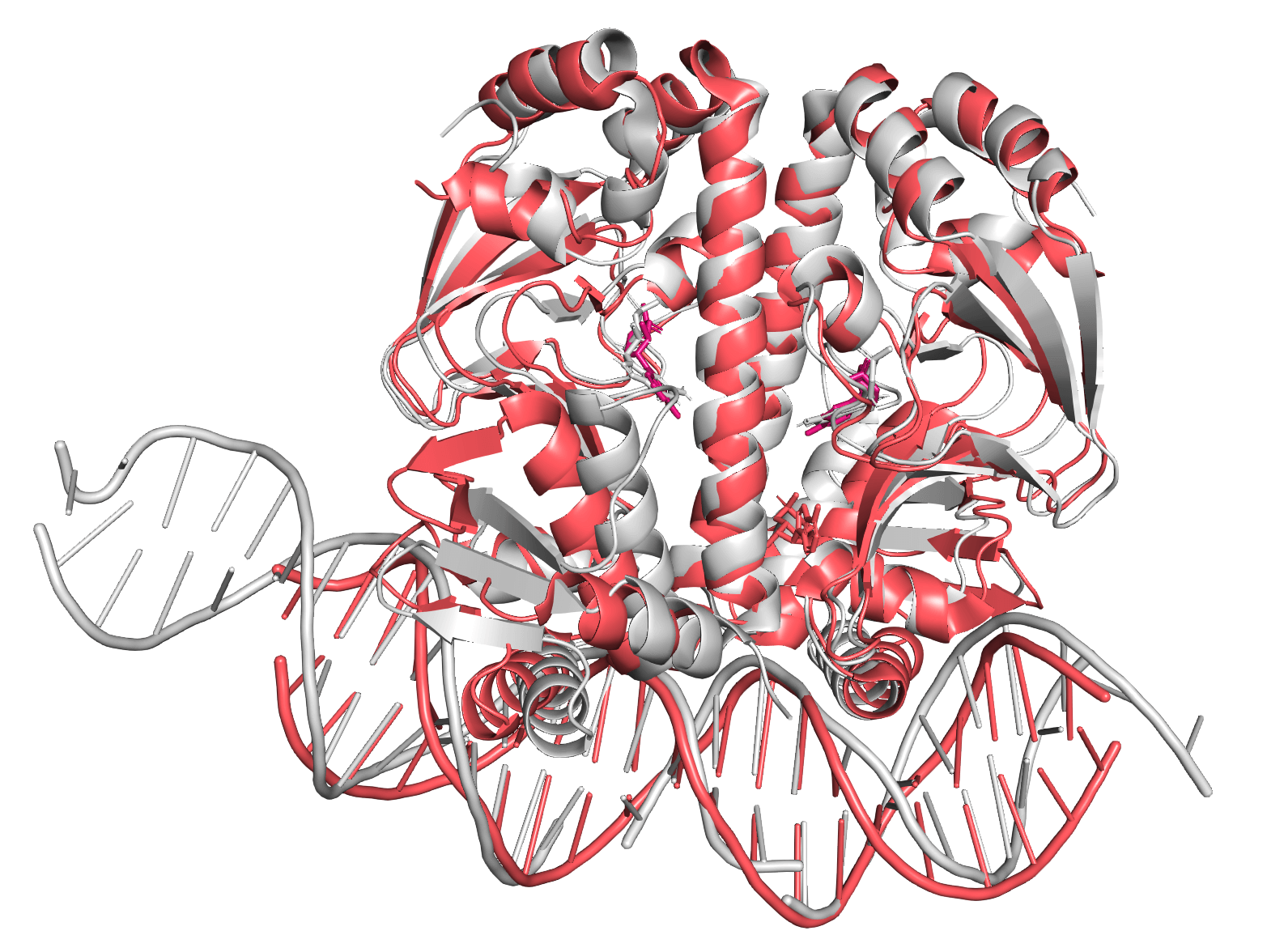

### limitation_pl_7xfa_1_v3.png

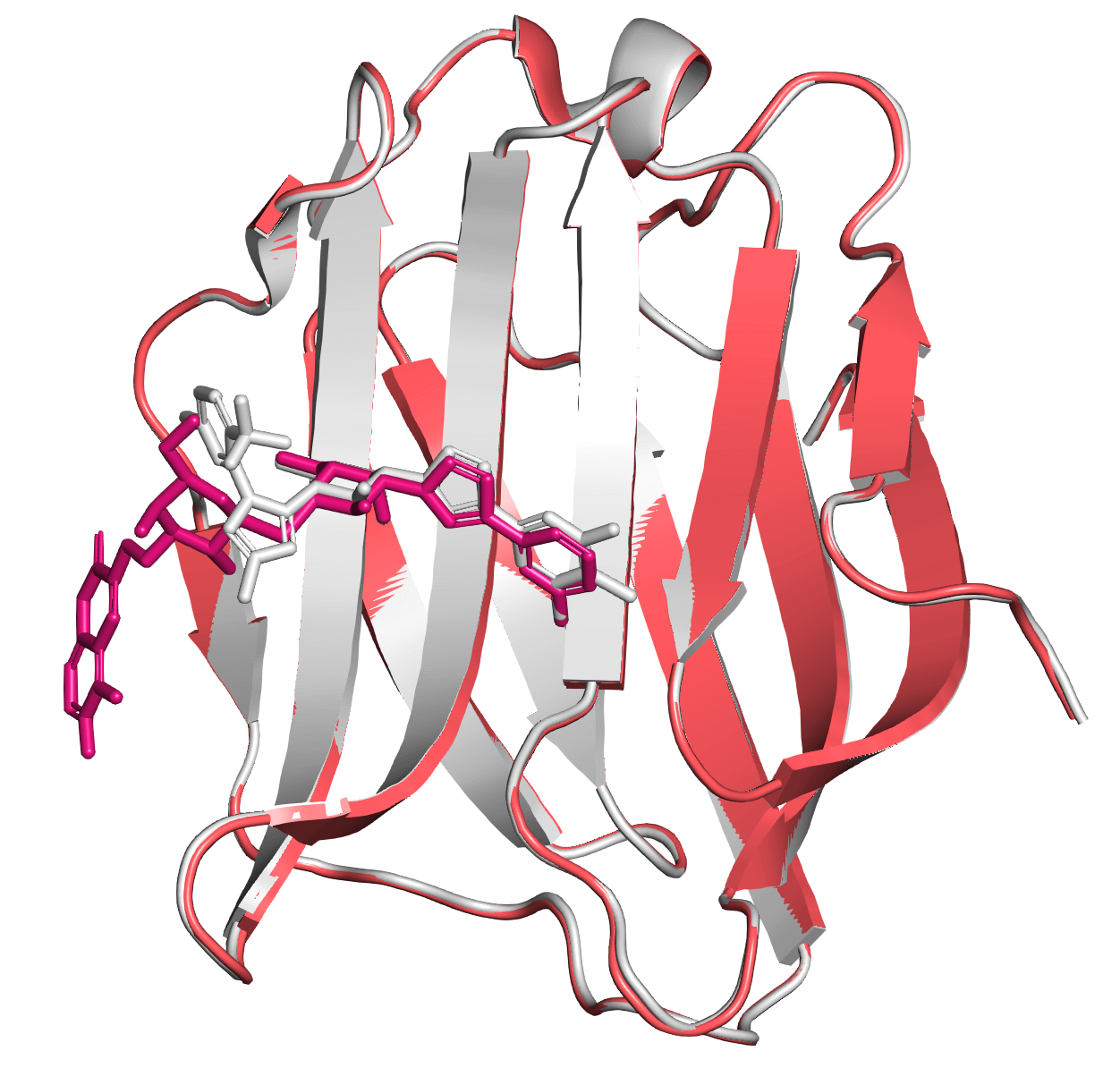

### limitation_pl_7xfa_2_v3.png

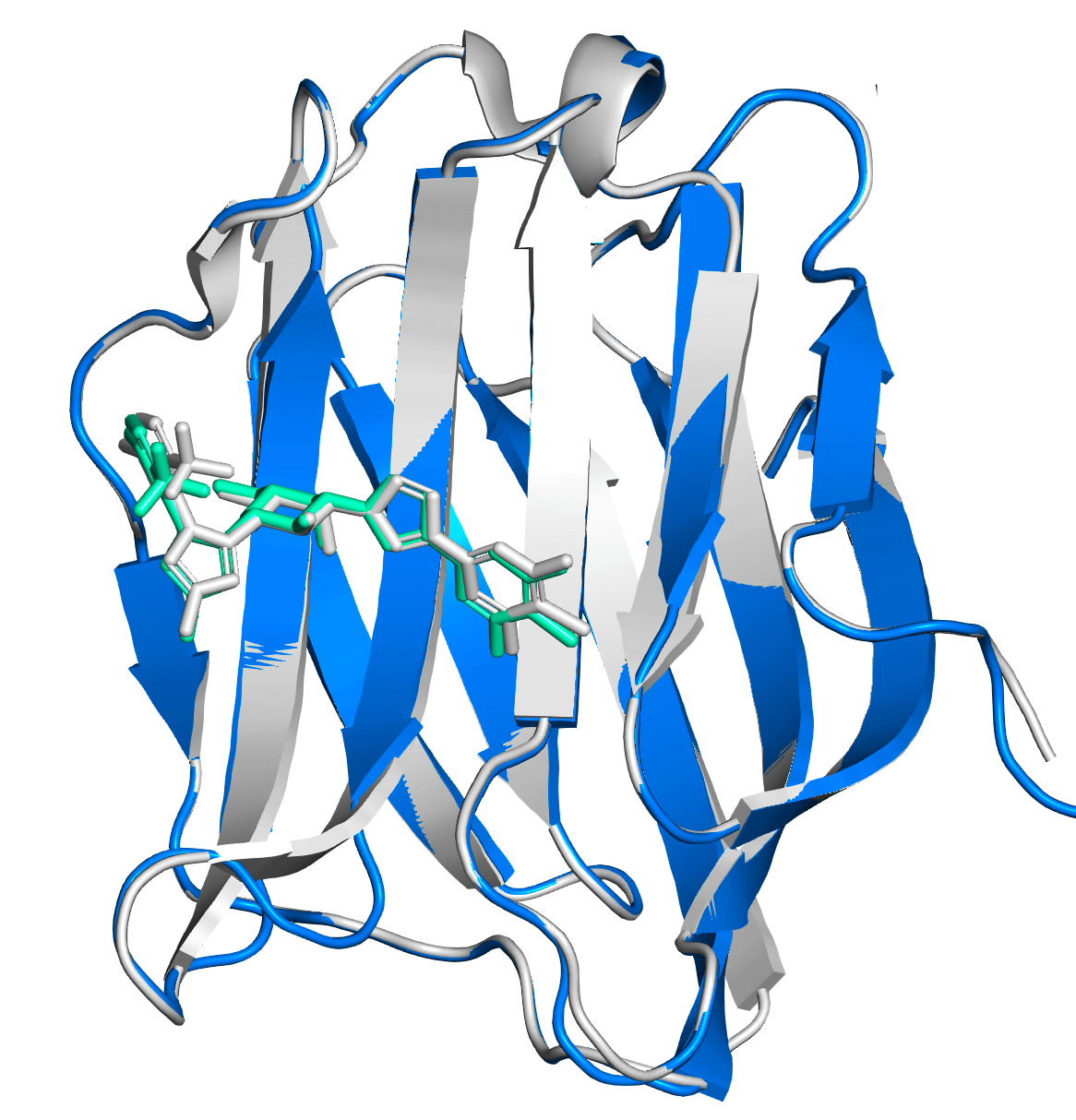

### nuc_compare_summary.pdf

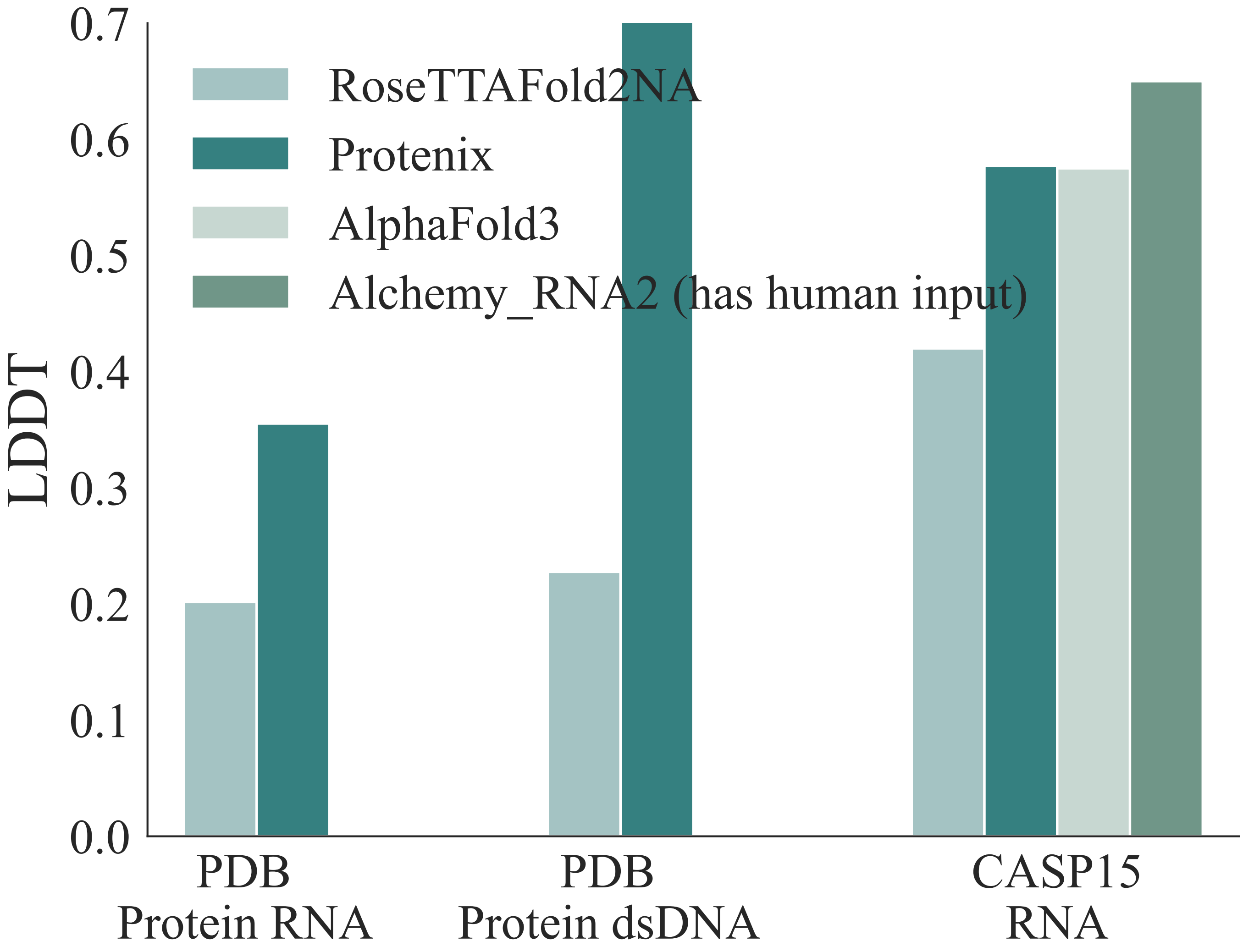

### nuc_good_case_7r6r_v2.png

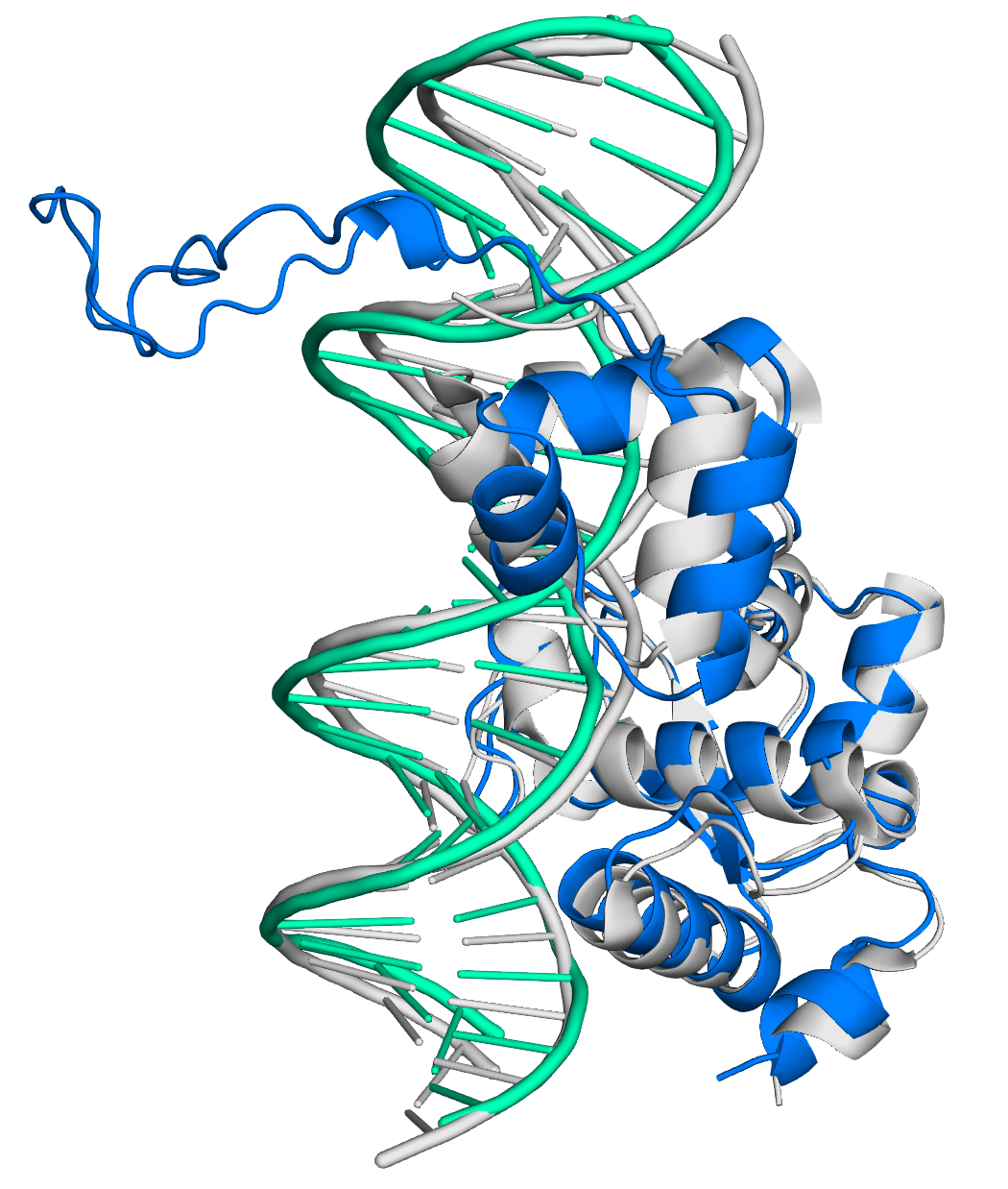

### nuc_prot.pdf

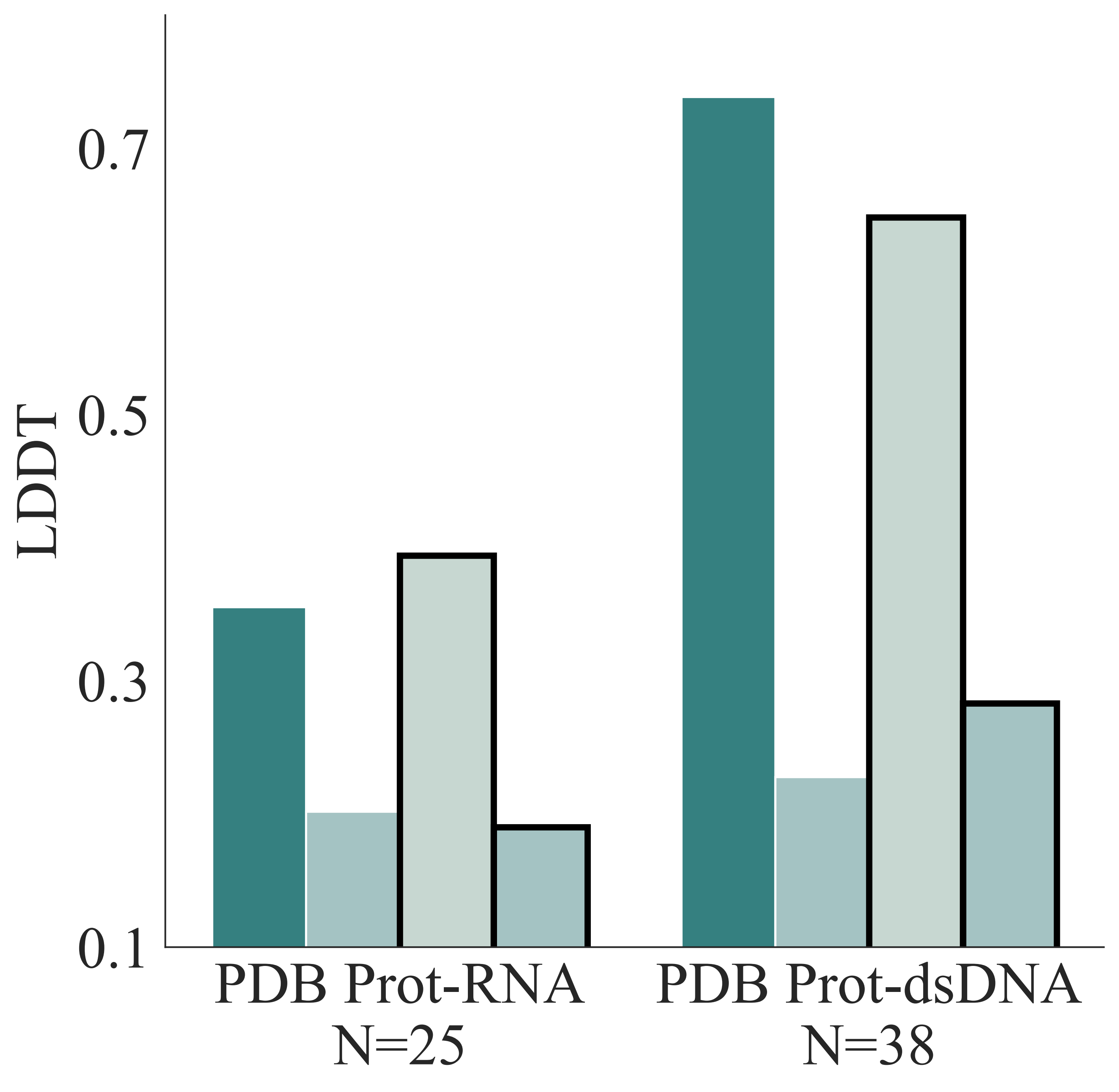

### nuc_prot_chain_iptm_dockq.pdf

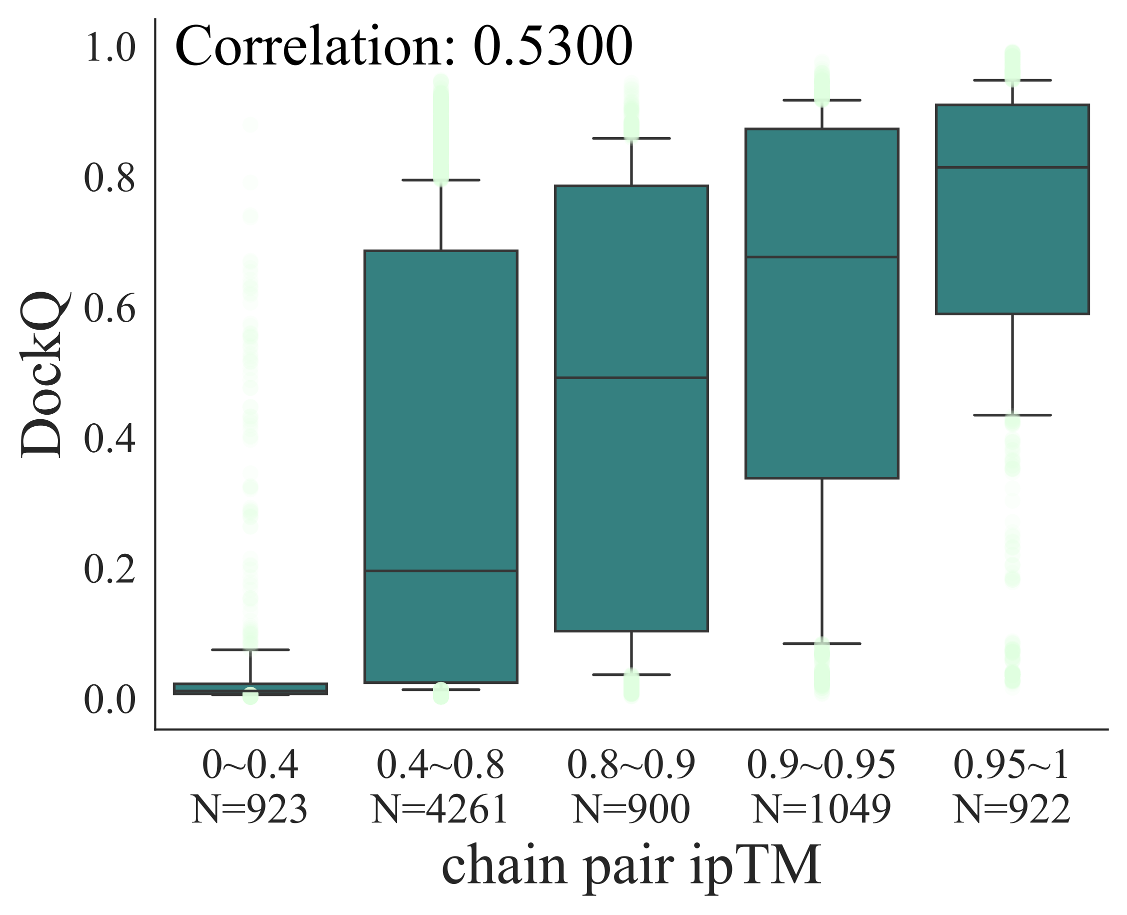

### overall_performance.pdf

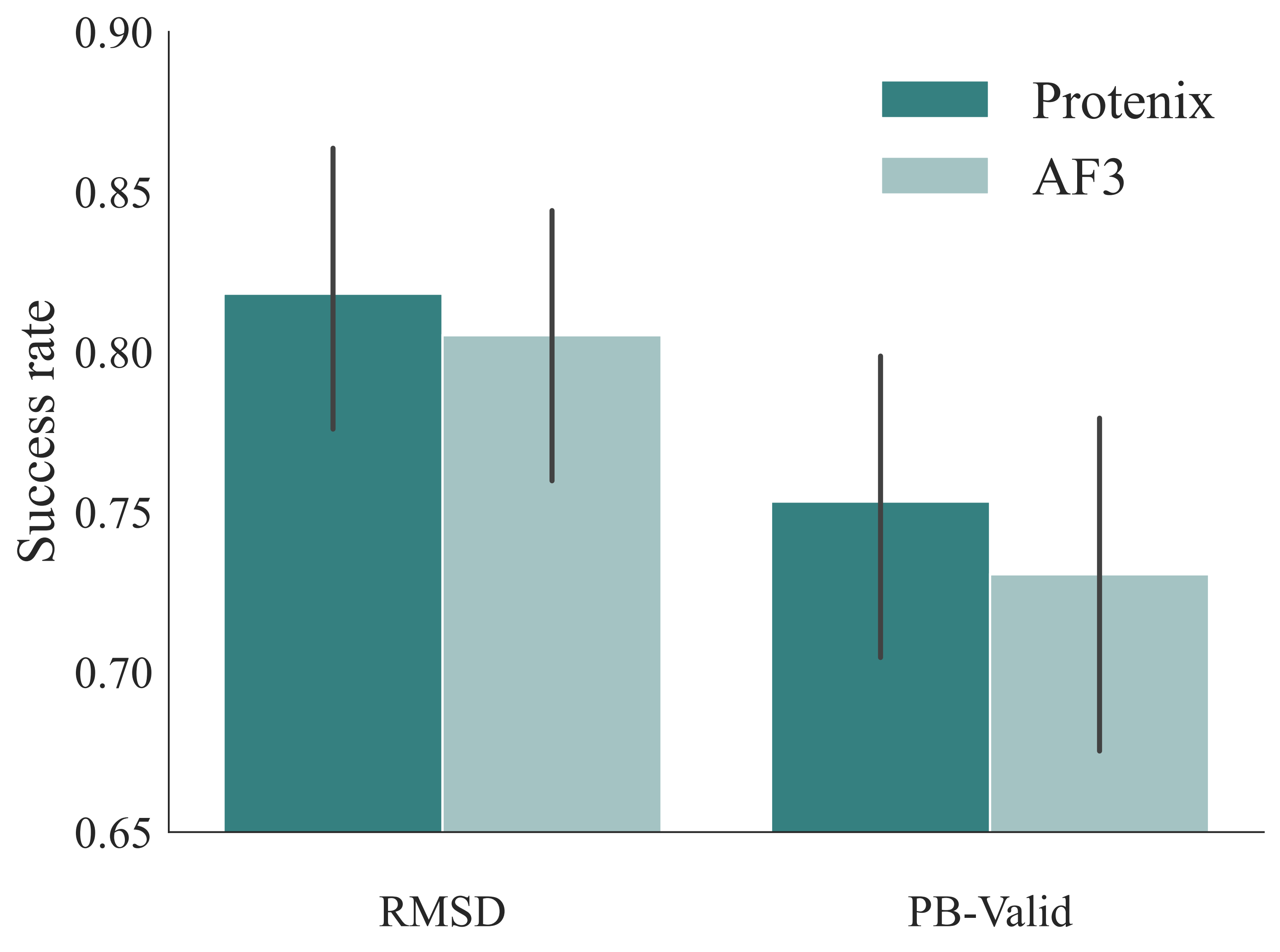

### pb-summary.pdf

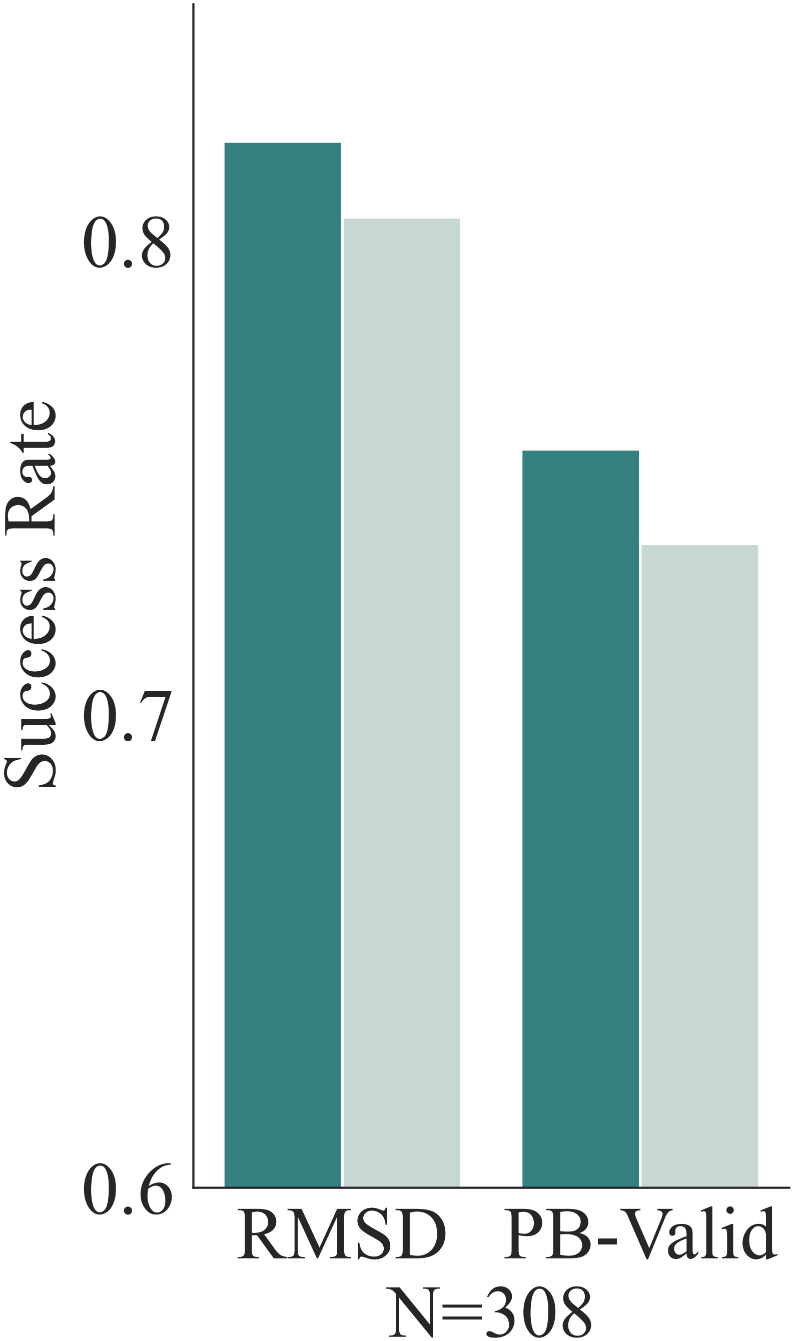

### pb_valid_detailed.pdf

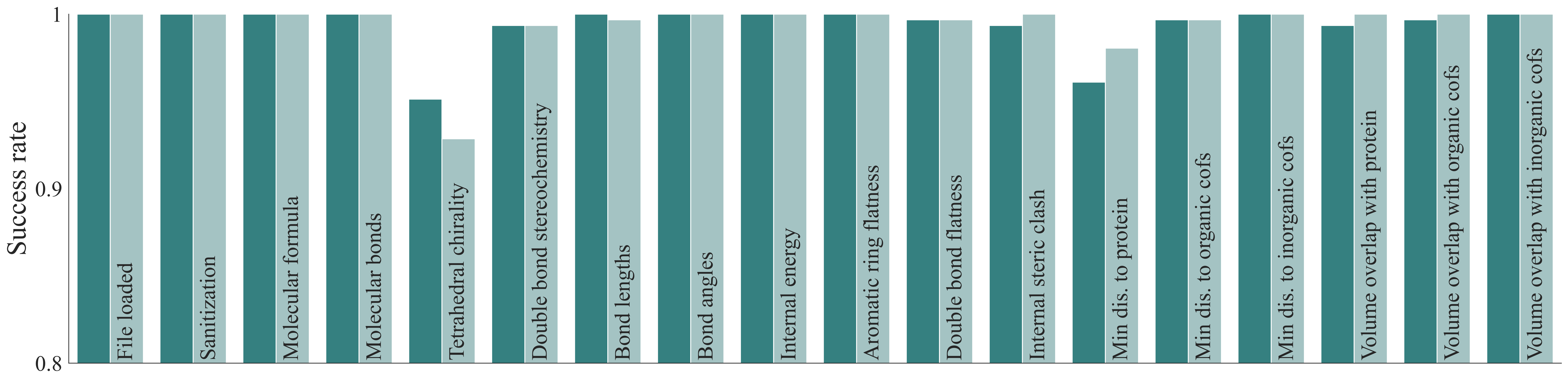

### prot-homology-detail.pdf

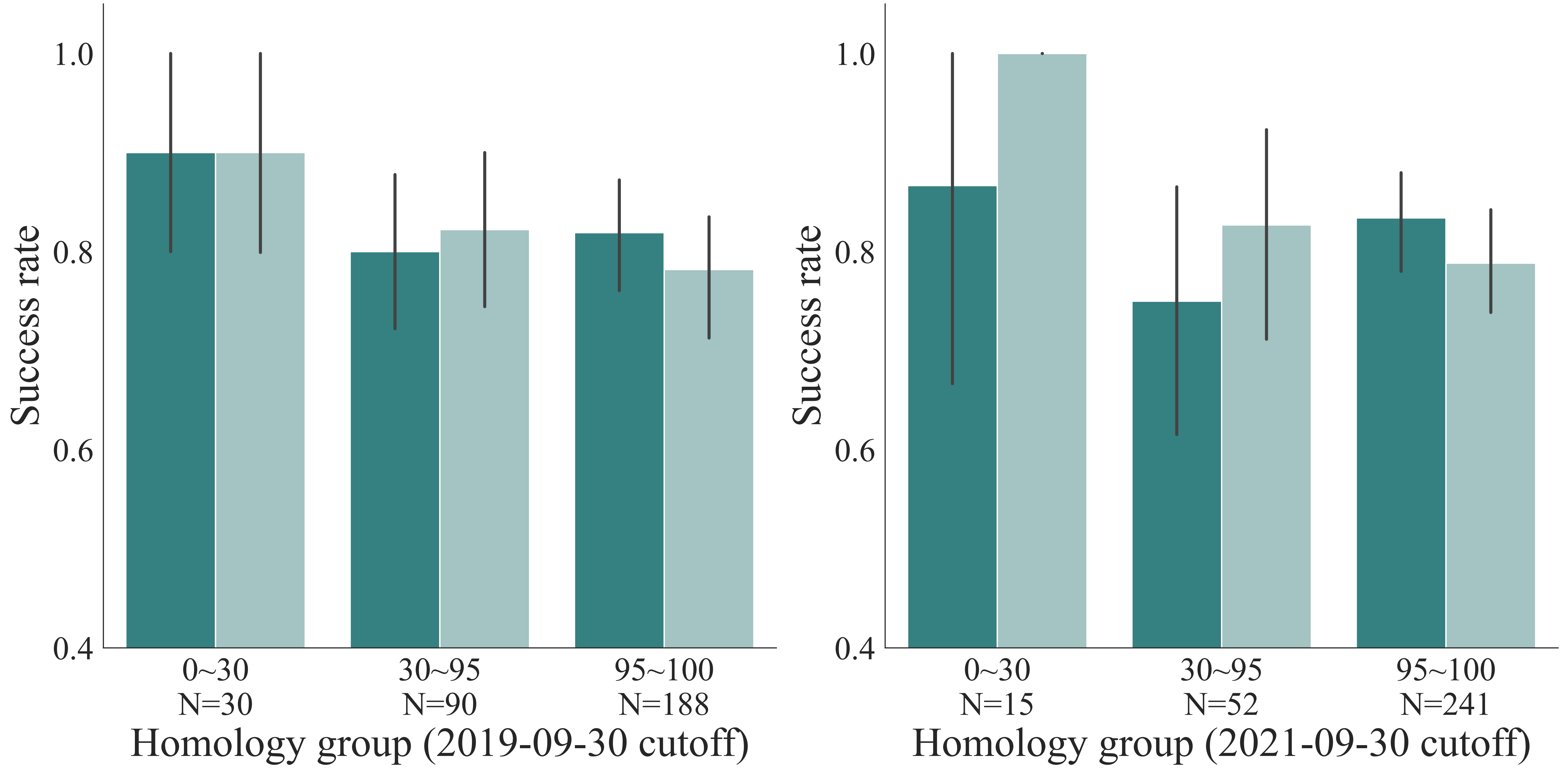

### prot-summary.pdf

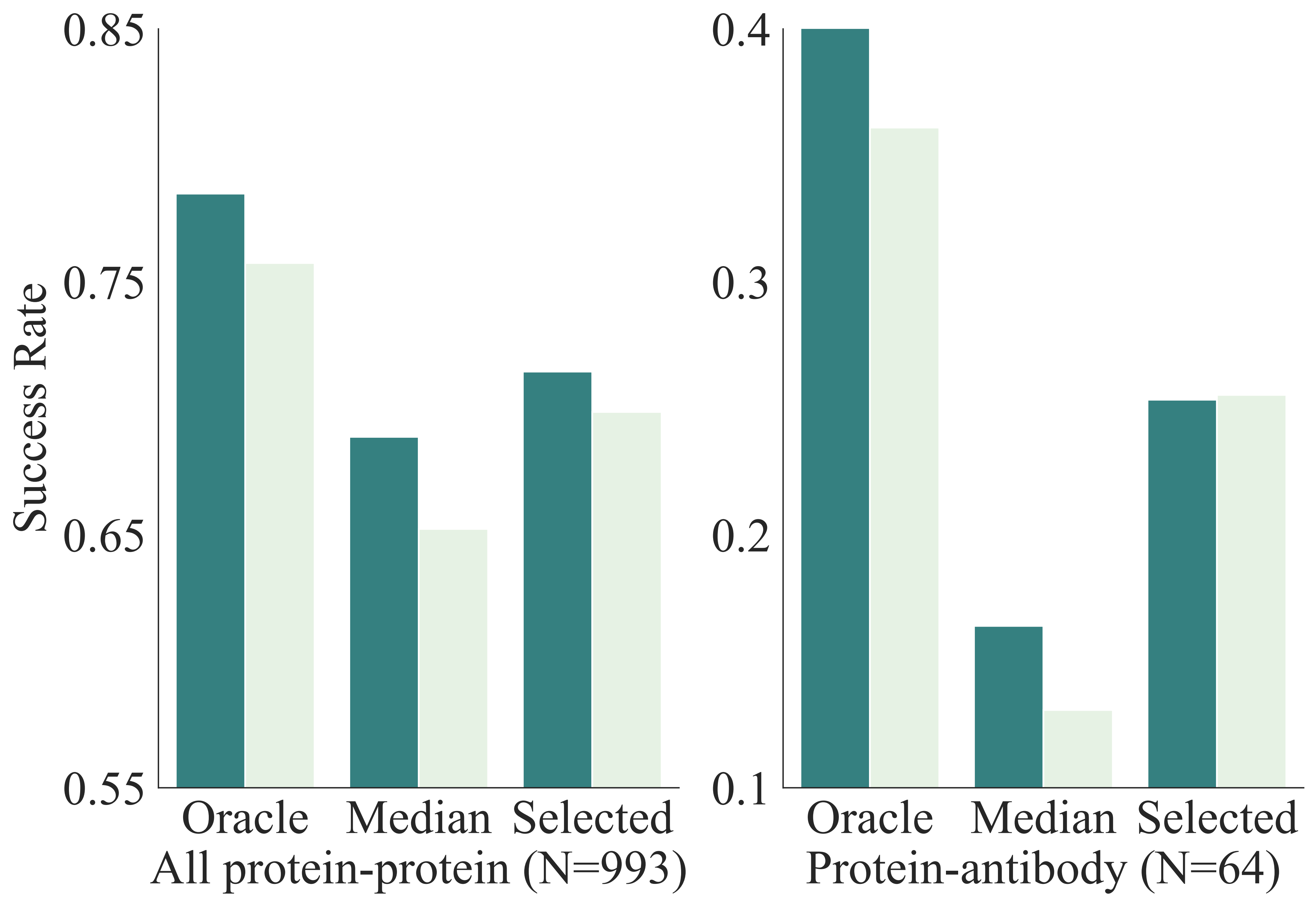

### prot_prot_chain_iptm_dockq.pdf

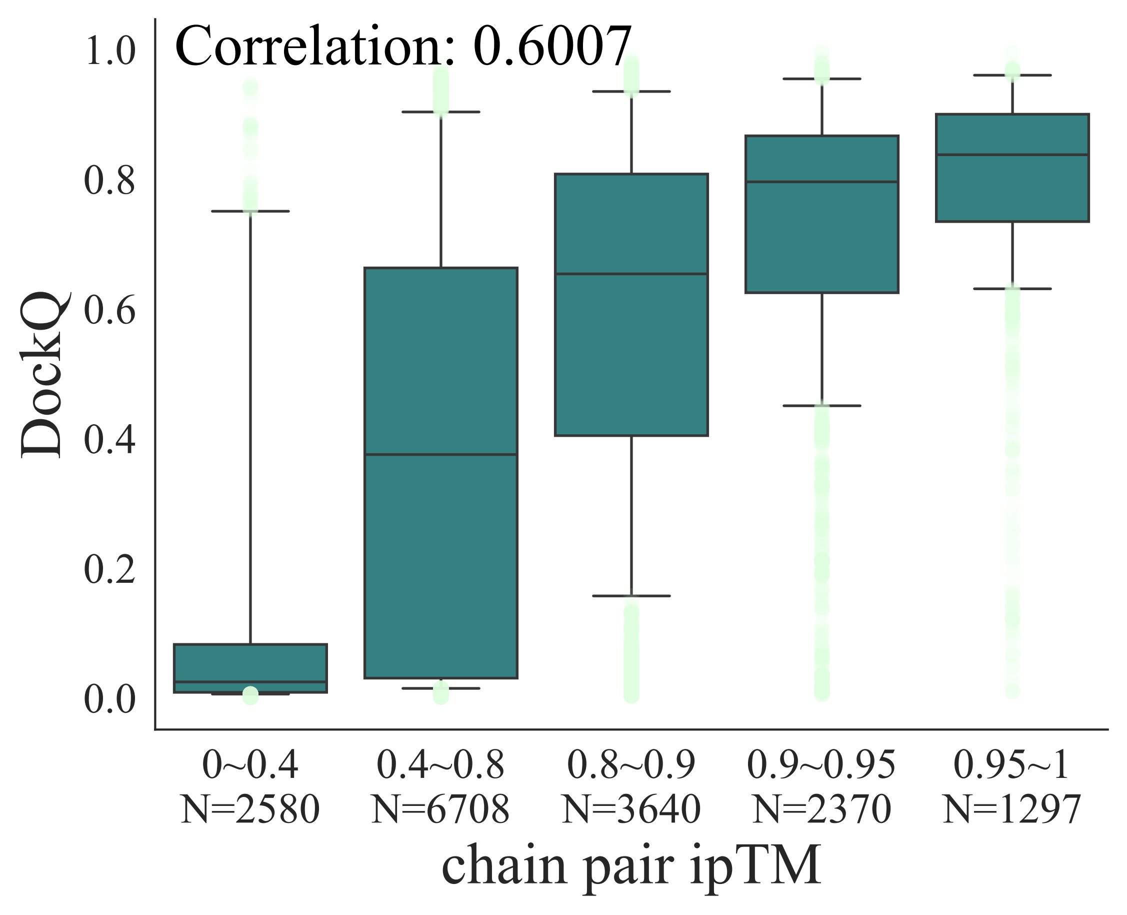

### protein_dna_good_case_7xv6_v2.png

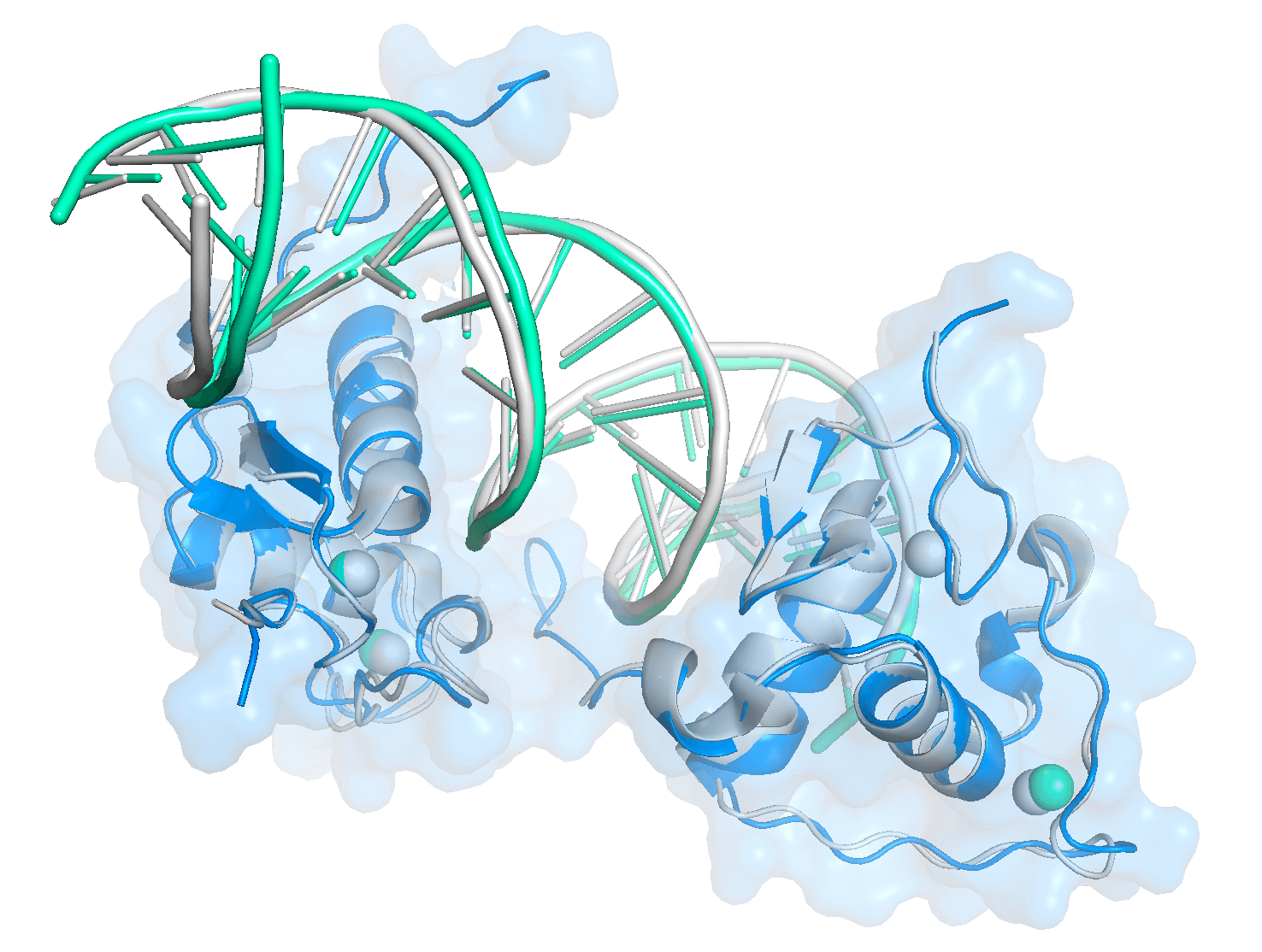

### protein_good_case_7xva_v3.jpg

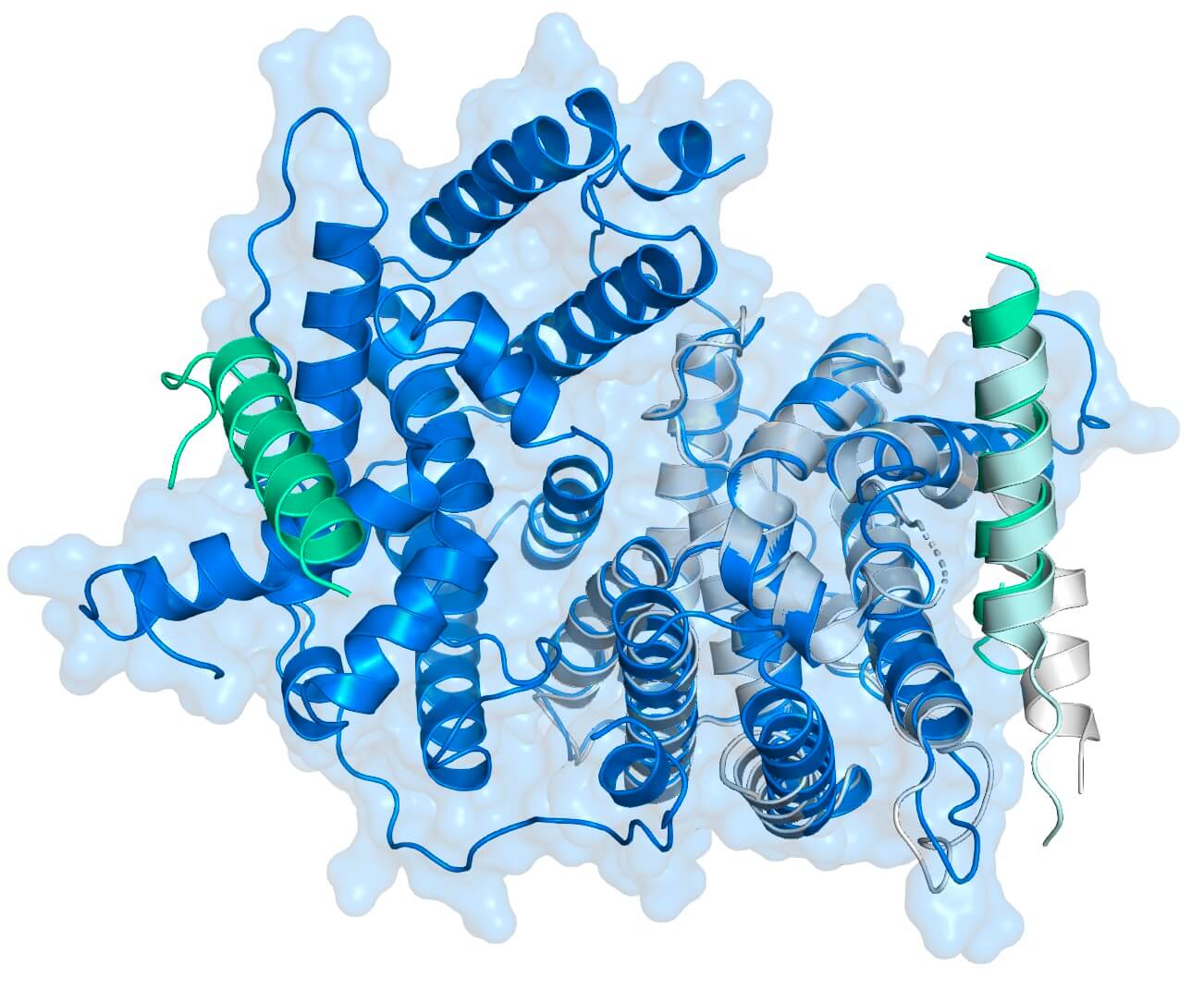

### rmsd_with_curve.pdf

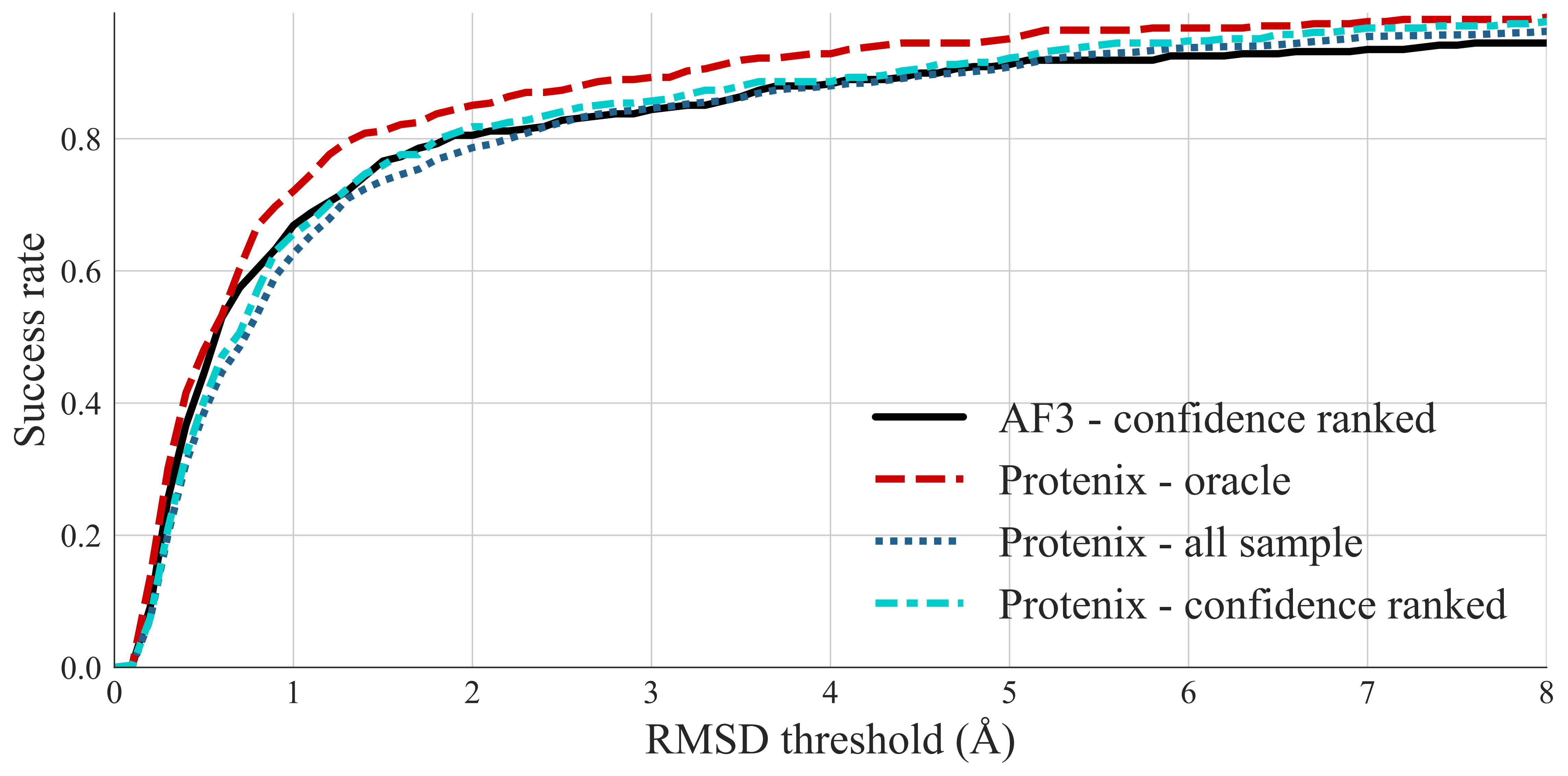
